# Extensive genetic variation of leaf specialized metabolites in sessile oak (*Quercus petraea*) populations

**DOI:** 10.1101/2023.04.07.536008

**Authors:** Domitille Coq--Etchegaray, Stéphane Bernillon, Grégoire Le-Provost, Antoine Kremer, Alexis Ducousso, Céline Lalanne, Fabrice Bonne, Annick Moing, Christophe Plomion, Benjamin Brachi

## Abstract

Specialized or secondary metabolites play a key role in plant resistance against abiotic stresses and defences against bioaggressors. For example, in sessile oaks *Quercus petraea*, phenolics contribute to reduce herbivore damage and improve drought resistance. Here, we explored the natural variation of specialized metabolites in nine European provenances of sessile oaks and aimed to detect its underlying genetic bases. We sampled mature leaves from high and low branches on 225 sessile oak trees located in a common garden and used untargeted metabolomics to characterise the variation of 217 specialized metabolites. In addition, we used whole genome low-depth sequencing to genotype individuals for 1.4M genetic markers. We found that leaf specialized metabolites displayed extensive within-provenance variation, but very little differentiation between provenances. In addition, a genome-wide association study allowed detecting significant associations for 42% of these metabolites. Hence, our results suggest that genetic variation for most leaf specialized metabolites is unlikely to be locally adaptive, however lack of differentiation among populations suggests selection acts locally to maintain diversity at loci associated with leaf specialized metabolites variation.

## Introduction

One of the crucial mechanisms developed by plants to interact with their environment is the production of molecules, known as specialized or secondary metabolites. Each plant species produces thousands of different molecules, which range from extremely specific to shared across the plant kingdom (De Luca & St Pierre 2000). In leaves, certain compounds, including tannins, flavonoids and glucosinolates were shown to have defensive effects against herbivores, including insects but also vertebrates (see Dearing, Foley & McLean 2005 for a review). Some compounds have become cues for specialist herbivores to find host plants (Wink 2018) and others attract the predators of herbivores, hence reducing damage to the plant (McCormick, Unsicker & Gershenzon 2012). Beyond interactions with herbivores, some specialized metabolites have antimicrobial properties and influence the leaf microbial community (Bailey *et al*. 2005) like certain flavonoids in carnation (Galeotti, Barile, Curir, Dolci & Lanzotti 2008) and tomato (Vargas *et al*. 2013). In addition, specialized metabolites production is also tightly linked to the plant immune system and phytohormones (Bednarek 2012), making them key players in plant pathogen interactions and good candidates for quantitative resistance and even exapted resistance (Newcombe 1998; Bartholomé *et al*. 2020). Beyond the leaf, in root exudates, specialized metabolites including flavonoids help attract beneficial microbes and promote mycorrhizal symbiosis (reviewed in (Walker, Bais, Grotewold & Vivanco 2003; Sebastiana *et al*. 2021), hence enhancing growth and water-use efficiency.

The effects of specialized metabolites even reach beyond the plant boundaries and influence the local biotic environment. Differences were detected in insect communities in the canopies of tree species producing different tannins (Forkner, Marquis & Lill 2004), molecules which were also found to impact the litter microbial community (Schweitzer *et al*. 2008). Finally, plant specialized metabolites play important roles for the plant protection against abiotic stresses (Sardans, Peñuelas & Rivas-Ubach 2011). Many specialized compounds have *in vitro* antioxidant properties and may help scavenge reactive oxygen species (ROS) *in vivo* (Nakabayashi *et al*. 2014). Thus, certain specialized metabolites, in particular anthocyanins, protect plants from the negative effects of UV light (Rozema *et al*. 2002), but can also enhance drought, heat and cold tolerance (Obata *et al*. 2015).

Individuals within a plant species do not produce a homogeneous set of compounds and different chemotypes can emerge. Multiple studies have shown that the quantities and structure of specialized metabolites vary within species, either because they are plastic and respond to the environment, or because of genetic differences between individuals. For example, in Arabidopsis and multiple members of the *Brassicaceae* family, glucosinolates and their metabolic pathways have been broadly investigated. Extensive genetic variation for specialized metabolites, such as glucosinolates, is present within species, with some compounds displaying broad geographical clines, suggesting adaptation to local herbivore communities (Züst *et al*. 2012; Brachi *et al*. 2015).

Differences in metabolic profiles were found among natural pedunculate oak (*Quercus robur*) stands in Germany, but interestingly, all stands included resistant or susceptible oak trees to herbivory (Bertić *et al*. 2021). In fact, oak trees from a single forest could be categorised as resistant or susceptible to defoliation by *Tortrix viridana*, based on the presence/concentration of specific compounds in their leaves (Kersten et al., 2013). Leaves from resistant oak trees were enriched in defence-related polyphenolic compounds, while leaves from the susceptible oaks were enriched in growth-related substances such as carbohydrates and amino-acid derivatives. These results are therefore consistent with both the adaptation of populations to local environmental conditions, but also the maintenance, within stands, of variation in the relative investment of trees in defence and growth.

European white oaks cover a large portion of European temperate forests and the decline of many populations, likely accelerated by climate change, could have dramatic consequences for the ecosystem services they provide including biodiversity. Indeed, a recent study showed that oaks are associated with 2,300 species in the UK, including birds, mammals, bryophytes, fungi, invertebrates and lichens, with 326 obligate species, including fungi, invertebrates and lichens (Mitchell *et al*. 2019). Many of these interactions, and the central role of oak trees in forest ecosystems, could be partly mediated by specialized metabolites. With oak populations suffering from decline related to both abiotic and biotic stresses imposed by global change, studying the natural variation and genetic bases of leaf specialized metabolites is essential to better understand the evolutionary history of specialized metabolites variation in oak populations. In particular, quantifying how much of this variation is locally adaptive and how much is maintained within population, is an important addition to the study of traits adaptive to climate, such as phenology or drought tolerance (Sáenz-Romero *et al*. 2017; Torres-Ruiz *et al*. 2019), to estimate the adaptive potential of oak populations.

In this study, we investigated the natural variation and the genetic bases of non-volatile leaf specialized metabolites in nine European sessile oak (*Quercus petraea* (Matt.) Liebl.) provenances growing in a common garden, using untargeted metabolomics. We started by characterising the genetic structure of oak populations and examined the differentiation between provenances for leaf specialized metabolites. We then performed a genome-wide association study to identify the genetic bases and architectures of leaf specialized metabolites. We investigated candidate genes and molecular structures for metabolites presenting strong associations and interesting patterns of variation. Our analyses revealed very little differentiation among provenances of the metabolites quantified, with the vast majority of the phenotypic variation being present within provenances. Our genome-wide association study identified significant associations for 42% of the 217 metabolites we quantified, suggesting that leaf specialized metabolite variation was, to a large extent,genetically determined, and that the variation of individual molecules displayed mono- to oligogenic architectures. The fact that the vast majority of the molecules with significant associations did not display differentiation between populations suggests that most molecules did not contribute to local adaptation among provenances, at a European geographical scale.

## Materials and Methods

### Sampling sessile oak leaves

We sampled sessile oak trees grown in a common garden experiment located in Eastern France (Sillegny, 48°59′ 24″ N 06°07′ 56″ E). This common garden is one of the four long-term common gardens including trees from 106 populations of sessile oak *Quercus petraea* Matt. Liebl. (Ducousso *et al*. 2022).

From the 106 populations represented in the common garden, we selected nine populations along a latitudinal gradient spanning from the South-West of France to the North of Germany (**Figure 1a**, Table **S1**). We collected leaves in the common garden on September 7^th^ and 8^th^, 2016 on 225 trees (22 to 28 trees per population). At the time of sampling, trees were between 29-35 years old. For each tree, we sampled four to six fully developed leaves from branches at two heights: low branches, mostly protected from direct sunlight by the canopy, and high branches exposed to sunlight. Leaves were collected either using a pole pruner when possible, or with a shotgun whenever the branches were too high to reach. Four to six leaves per branch were stored in 20 mL plastic vials and frozen on dry-ice upon harvest (see Supplementary Material and Methods S1). We collected a total of 582 samples. These samples were used to generate three datasets: genome-wide SNP genotypes of oak trees, a “GWA dataset” and an “annotation dataset”. The production and analysis of these three datasets is described in the following and illustrated in Figure S1.

**Figure 1.**
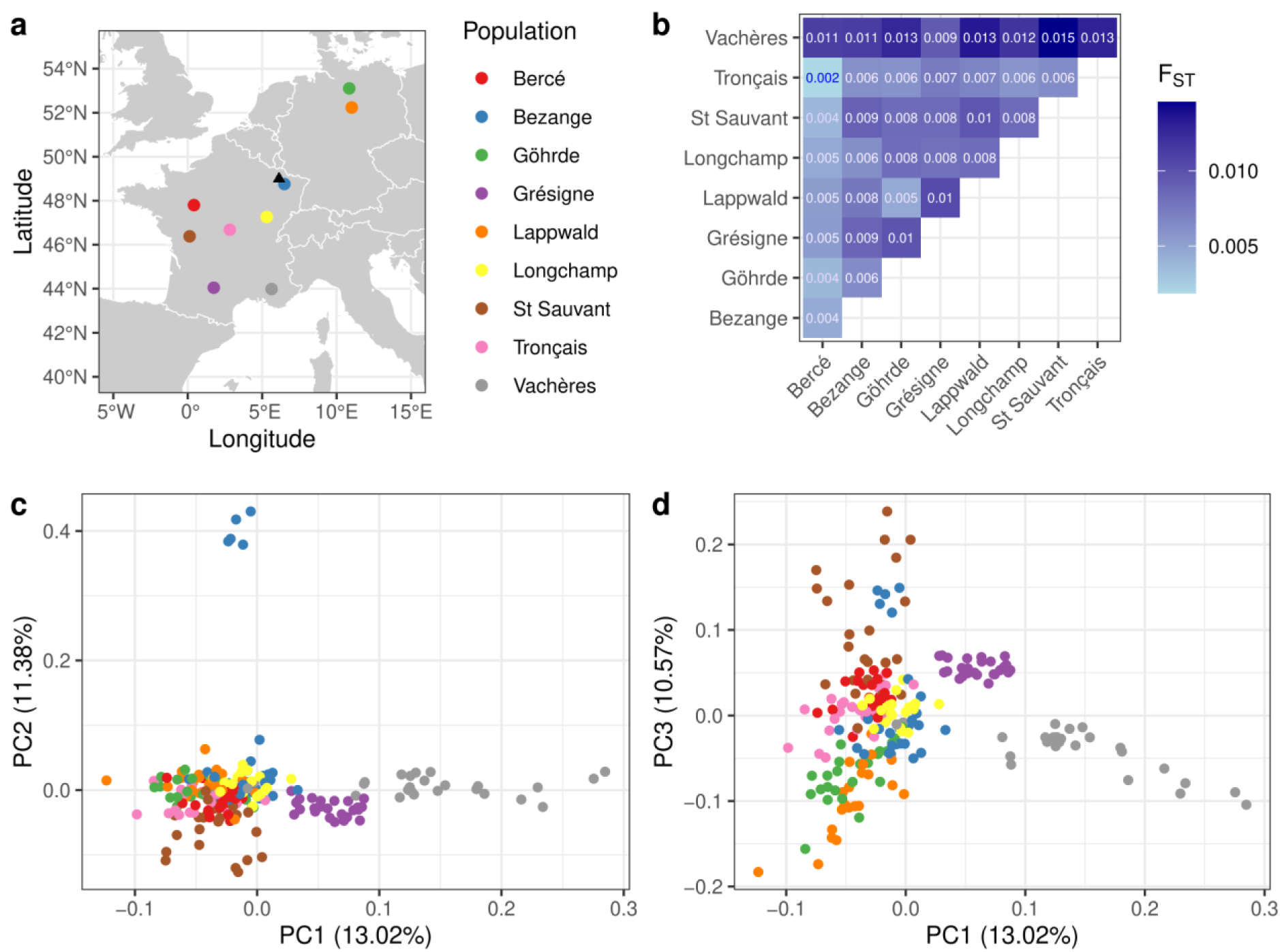
Structure and differentiation of sessile oak populations. **(a)** Map of sessile oak populations included in the study. The coloured dots on the map correspond to the geographical origin of the 9 populations we sampled. The location of the common garden in which the trees are installed is marked by the black triangle. **(b)** Hudson pairwise FST between populations. Populations are indicated along the x and y axes. FST between pairs of populations are indicated in the half-matrix. The cells in the matrix are coloured in darker shades of blue for high FST values. **(c-d)** Genetic variation of oak populations visualised using a PCA. Each point corresponds to a tree (N=218), projected in the plane defined by the first three components **(c)** PC1, PC2 and **(d)** PC1, PC3 of a principal component analysis computed using 356,413 un-correlated (r^2^ < 0.2) SNPs. Points are coloured according to populations as in **(a)**.

### Sequencing and genotyping of sessile oak individuals

Briefly, we extracted DNA from freeze dried leaf material and sequenced 224 individuals to a depth of ∼10X using the Illumina NovaSeq system (details are provided in the Supplementary Material and Methods S2 and detailed sample counts in Table **S2a**).

We used a home-made bioinformatic pipeline, developed with Snakemake v5.8.1 workflow manager (Köster *et al*. 2021) and Singularity containers (Kurtzer, Sochat & Bauer 2017). We removed Illumina TrueSeq adapters and trimmed reads using cutadapt v1.18 (Martin 2011) and sickle v1.33 (Joshi & Fass 2011). We aligned paired-end reads with bwa-mem2 v2.2.1 (Li 2013) on *Quercus robur* reference genome (Plomion *et al*. 2018). We marked duplicated reads using GATK v4.2.4.0 and clipped overlapping read pairs using BamUtils v1.0.15 (Jun, Wing, Abecasis & Kang 2015), following recommendation in GATK best practices (Van der Auwera *et al*. 2013). We created mpileup files of the 224 individuals using samtools v1.9 (Danecek *et al*. 2021) and converted them into pro files using sam2pro v0.8. From the pro files, we summarized nucleotide read quartets at each position of the reference genome (Plomion *et al*. 2018) and detected bi-allelic SNPs using a genotype-frequency estimator (GFE) (Maruki & Lynch 2015). Then, we called genotypes using the Bayesian genotype caller (BGC) described in (Maruki & Lynch 2017). We discarded SNPs with a minor allele frequency (MAF) lower than 10%, with more than 10% of missing calls, and located in regions annotated as transposable elements in the Q. *robur* reference genome. These filters were achieved with a combination of awk command lines and PLINK v1.9 (Chang *et al*. 2015). We discarded three pairs of individuals with genome-wide genetic similarity above 95% (i.e 95% of SNPs with identical genotypes between the two individuals) as they may have corresponded to the same individual sequenced twice (Table **S2a**).

### Population genomics analysis of sessile oak populations

We estimated linkage disequilibrium (LD) decay along the oak genome using the r² function available in PLINK v1.9 (Chang *et al*. 2015) with window size of 50kb.

To study the genetic structure of *Q. petraea* populations we pruned the SNPs according to LD previously estimated and removed highly correlated markers (r² > 0.2) using the “--indep-pairwise” function of PLINK v1.9 (Chang *et al*. 2015) with a window size of 2 kb and a step-size of 1. We computed an average Hudson fixation index (F_ST_) (Bhatia, Patterson, Sankararaman & Price 2013) between pairs of populations, estimated using all SNPs in the pruned dataset using the “--fst” PLINK v2.0 (Chang *et al*. 2015) with default parameters. To investigate global patterns of population structure, we performed a principal component analysis using the pruned dataset and the “--pca” function in PLINK v1.9 (Chang *et al*. 2015). Finally, we computed F_ST_ ^(^Weir & Cockerham 1984^)^ across populations using the “--fst” function in PLINK v2.0 (Chang *et al*. 2015) for all 1.4M markers.

### Metabolomics: extraction, data acquisition, processing and filtering

#### Metabolomics for annotation

#### Extraction and LC-MS analyses

Specialized metabolites were extracted using 70% methanol (see Supplementary Material and Methods S3). After the extraction, Quality Controls (QCs) were prepared by combining 2µl of methanolic extract from each sample into a single tube (excluding blanks and technical replicates).

This extraction included a total of 259 leaf samples. These 259 samples corresponded to: 1) leaves from high and low branches from a subsample of 54 trees, including all nine provenances, collected in 2016, plus five technical replicates extracted for five of the samples, 2) samples from high and low branches from the same 54 trees collected during a second sampling in 2021, plus five technical replicates extracted from four random samples. Quality Controls (QCs) were prepared by combining 2µl of methanolic extract from each sample into a single tube (excluding blanks and technical replicates). These samples were randomised across three 96-well plates, which also included 26 extractions blanks and 3 QCs (Table **S2b**). The goal was to obtain fresh extracts for a subset of the samples collected in 2016 and compare leaf specialized metabolites of the same trees, sampled over two years. In the present study, we only used samples collected in 2016 from this extraction for annotation purposes (see below).

#### Annotation dataset

This dataset was generated from 2016 leaf samples and used to perform automated annotations of specialized metabolites. The separation of extracts was achieved using reverse phase liquid chromatography with a C18 column over a 10 min gradient (see Supplementary Material and Methods S3). The chromatography flow was then directed to a LTQ-Orbitrap Elite equipped with a HESI probe (ThermoScientific, Bremen, Germany).

Data was acquired in data dependent mode to generate MS² spectra for selected ions and each extract was injected twice: once with the ESI set to positive mode and once with the ESI set to negative mode. (Parameter details provided in Supplementary Material and Methods S4).

#### Data treatment and filtering of the “annotation dataset”

We used the “annotation dataset” to annotate pseudo-molecules using an integrated metabolomic workflow (Fraisier- Vannier *et al*. 2020). First, we performed data alignment on a QC file injected in the middle of the injection sequence and spectral deconvolution using MS-DIAL v4.60 (Tsugawa *et al*. 2015) for the positive and negative ion modes. We used the following parameters: centroid mode, MS1 and MS2 tolerances at 0.01 and 0.025 Da, RT tolerance of 0.1 min and a mass tolerance of 0.01. Then, we filtered MS-DIAL data output with MS-CleanR v1.0 with default parameters (Fraisier-Vannier *et al*. 2020). We annotated filtered data with MS-FINDER v3.50 with MS1 and MS2 tolerances at 5 and 15 Da (Tsugawa *et al*. 2016). We searched formulas with C, H, O, N, P and S atoms from PlantCyc (https://plantcyc.org/) and KNApSAck (http://www.knapsackfamily.com/) databases. To obtain putative annotations of pseudo-molecules quantified in the “GWA dataset” (see below), we matched *m/z* and retention time (RT) with the molecules in the “annotation dataset” with a *m/z* tolerance of 0.03 and an RT tolerance of 0.6 min (Dataset S2). The raw data files for the annotation dataset were also used for manual verification of putative annotations for pseudo-molecules of interest by comparing parent ions *m/z* to databases and inspecting fragmentation patterns.

#### Metabolomics for GWA

#### Extraction and LC-MS analyses

An extraction of specialized metabolites was performed on 661 leaf samples and 99 extraction blanks (no leaf material) in 2016 (Table **S2c**), following the same protocol that was presented above. Leaf samples included one sample per tree and per height, as well as biological replicates (additional leaf samples from random trees, not exploited in this study), samples taken on the same trees at different times during the field sampling (not used in this study) and technical replicates. Technical replicates consist of 20 random leaf samples that were extracted five to six times. In total, this extraction included 760 samples randomised across eight 96-well plates (Table **S2c**). Note that the randomisation across plates was designed to include all provenances and heights combinations in all plates. Specialized metabolites were extracted using 70% methanol (see Supplementary Material and Methods S3). After the extraction, Quality Controls (QCs) were prepared by combining 2µl of methanolic extract from each sample into a single tube (excluding blanks and technical replicates).

#### GWA dataset

This dataset was used to screen specialized metabolite variation and perform genome-wide associations analyses and will be designated hereafter as the “GWA dataset”. The liquid chromatography protocol was identical to the liquid chromatography performed for the annotation dataset (see “Liquid Chromatography” in Supplementary material and methods S3). For the “GWA dataset”, the chromatography flow was directed to an ESI probe set to positive mode (-500 V endplate offset, +3.5 kV capillary voltage, 2.4 bar nebulizer, dry gas flow of 8.0 L/min at 190° C) into a hybrid quadrupole time-of-flight (QTOF) mass spectrometer (Bruker, Bremen, Germany). The mass-to-charge ratio ion scan was from *m/z* 50 to *m/z* 1500 with an acquisition frequency of 2 Hz. The mass spectrometer was *m/z*- calibrated with a 10 mM lithium formate solution injected at the begin and the end of each chromatogram. Using positive mode allowed detecting more metabolites, with better sensitivity. A QC was injected every 12 samples.

#### Data treatment and filtering of the GWA dataset

We converted raw proprietary files from the instrument to the mzML format using the ProteoWizard program (Chambers *et al*. 2012). We filtered resulting files to keep only data between 80 and 570 seconds using msConvert (Chambers *et al*. 2012) in order to remove calibration peaks and column wash offs at the beginning and the end of the runs. We removed 45 files from the eighth plate which had no calibration peaks. Past these initial processing steps, all analyses were carried out in a home- made bioinformatic pipeline using Snakemake workflow manager (Köster *et al*. 2021), Singularity containers (Kurtzer *et al*. 2017) and R v4.1.1 (R Core Team 2021). We used the IPO R package v1.18 (Libiseller *et al*. 2015) to optimise parameters for retention time (RT) correction and peak picking implemented in the R package XCMS v3.14 (Smith, Want, O’Maille, Abagyan & Siuzdak 2006) (Table **S3**) and added a noise threshold at 800. We ran the “findChromPeaks” function to detect chromatographic peaks. Then, we ran the “adjustRtime” function to correct RT deviation and calculated adjusted RTs using a subset-based alignment based on the QC samples injected at regular intervals. Finally, we ran the “PeakDensityParam” and “groupChromPeaks” functions to match all the detected peaks between samples. We produced a table of peak intensities and identified pseudo-molecules, based on RT and intensity correlations among peak groups using the “groupFWHM” and “groupCorr” functions of the CAMERA R package v1.48.0 (Kuhl, Tautenhahn, Böttcher, Larson & Neumann 2012).

After this step, we replaced NA values by zeros (as the fact that a molecule was undetected in a sample means that it was not present or at very low concentration and is biologically meaningful) and we filtered the dataset to remove outlying samples and peaks. First, we used the median peak intensity across all peak groups to remove samples that had values consistent with blanks and removed blanks that had median peak heights similar to regular samples.

Second, we filtered the peak groups defined by XCMS using QCs and blanks, with the rationale that peaks had to be significantly higher in the QCs than in the blanks to be retained. We used a one-way variance analysis (ANOVA, p-*value*<0.05) on log-transformed intensity data for each peak to determine the average difference in intensity between blanks and QCs and assess significance. Third we also removed all peaks with coefficients of variation above 30% in the QCs. Finally, for each injection, we checked that the internal standard included in the extraction buffer was still clearly present and that its intensity remained stable along the injection sequence (Figure **S2)**.

We then corrected peak intensities to account for two potentially confounding effects. First, in LC-MS analyses including many samples over long series, it is frequent to observe intensity drift. To correct the effect of intensity drift (although it was minimal in our analyses, see Figure **S2**) we divided the intensity values of each peak by the intensity of the corresponding peak in the closest QC injected after the sample along the injection sequence (a QC was injected every twelve samples in the injection sequence, see Supplementary Material and Methods S3). Second, the weight of dry material used in the extraction influences peak intensity. We therefore divided the peak intensities (relative to the values measured in the QCs) in each sample by the weight of material used in the extraction.

### Statistical analysis of leaf specialized metabolites variation

#### Specialized metabolite variation analysis

We studied peak intensities variation between low and high leaves and patterns of leaf metabolic profiles among populations measured in the “GWA dataset”. We transformed peak intensities using a cube root transformation and used a Pareto scaling and performed Principal Component Analysis (PCA) and a Partial Least-Square Discriminant Analysis (PLS-DA) on using MetaboAnalystR R package v4.0.0 (Chong & Xia 2018).

### Genome-wide association analysis

#### Genome-wide association studies

The genome-wide association analyses were performed on the transformed signal intensities of individual pseudo-molecules using a fixed and random model circulating probability unification method (farmCPU) (Liu, Huang, Fan, Buckler & Zhang 2016) implemented in the rMVP R package v1.0.6 (Yin *et al*. 2021). We used the “MVP” function and used the following parameters: maxLoop=10, method.bin= “static”, bin.size=c(5e2,5e3,5e4), permutation.threshold=TRUE, permutation.rep=1000 and threshold=0.05.

#### SNPs clusters

We clustered SNPs by regions based on the estimation of the linkage disequilibrium (∼3,000 bases). We considered significant associations identified by farmCPU (Liu *et al*. 2016), across all metabolites, and, walking along the genome, we grouped consecutive SNPs with significant associations in the same cluster if they were located less than 3000 bases apart. If the distance between consecutive significant SNPs was larger than 3000 bases a new cluster was created.

#### Gene annotation of highly associated SNPs

We searched genes within regions near SNPs highly associated with metabolite variation. Based on the estimated linkage disequilibrium decay distance (∼3,000 bases), we searched regions of 6,000 bases around SNPs with significant associations for candidate genes on the *Quercus robur* genome annotations (Plomion *et al*. 2018). When a candidate gene was identified, we extracted the gene sequence using samtools faidx v1.6 and used the blastx function of BLAST+ v2.12.0 (Camacho *et al*. 2009) with default parameters to find similarity between the candidate gene translated sequence and the *Arabidopsis* proteome. When no candidate was found, we extracted the entire 6000 bp) sequences and searched for similarities between the whole translated region sequence and the *Arabidopsis* proteome (Berardini *et al*. 2015).

## Results

### Population genomics of sessile oak populations

In this study we sampled 225 sessile oak trees originating from nine populations (**Figure 1a**) and sequenced 224 of these trees to a depth of 10X (on average 36.5M paired-end reads per tree). After filtering, we were able to genotype trees for 1,408,029 SNPs with a minor allele frequency above 10%, a missing genotype rate of 10% and after excluding regions annotated as transposable elements on the *Q. robur* reference genome (Plomion *et al*. 2018). We obtained an average SNP density of 170 SNPs per window of 100 kb, ranging from 0 to 923 SNPs, and 90% of 100 kb windows included over 9 SNPs. Three pairs of individuals displayed genetic similarity above 95% and were removed from the dataset, bringing the total number of individuals genotyped to 218.

Using these 1.4 million markers we estimated that linkage disequilibrium (LD), r², decreased to values below 0.2 over 2.9 kb in our collection of 9 populations spanning a large fraction of the species distribution range (Figure **S3**).

We then pruned the dataset down to 356,413 un-correlated SNPs (r²<0.2) in the 218 individuals to investigate population differentiation and genetic structure. First, we investigated the genetic differentiation between pairs of provenances using the Hudson fixation index F_ST_. We found that F_ST_ between pairs of populations was on average 0.008, ranging from 0.002 to 0.014 (**Figure 1b**). We observed the highest differentiation (F_ST_=0.0146) between two populations from France: Vachères and Saint-Sauvant (**Figure 1b**). Second, we investigated patterns of population structure using a PCA. The first principal component (PC) explained 13.02% of total variance and separated the two populations from the South-East of France (Vachères and Grésigne) from the other provenances (**Figure 1c**).

The second PC explained 11.38% of total variance and mostly captured the difference between a group of individuals from Bezange from the other individuals included in the study (**Figure 1c**). Inspection of kinship values shows that these individuals were likely half-sibs.

Overall, the PCA showed clustering of individuals according to their population of origin (**Figure 1c**). The third principal component explained 10.57% of the variance, and captured differences between the two most northern populations, from Germany, from the two most western populations in France (**Figure 1d).**

### Leaf specialized metabolites annotation and natural variation

We used high-throughput LC-MS analysis to measure the relative concentrations of leaf specialized metabolites in the leaves of the pedunculate oak trees from nine populations that we genotyped. The trees sampled are ∼25 years old, grow in a single common garden, and were sampled over two days at the end of the summer 2016. Specialized metabolites were extracted using a methanolic extraction and LC-MS analyses were run to produce two distinct datasets.

The first, dedicated to automated annotations of pseudo-molecules and referred to as the “annotation dataset”, was produced using a HPLC coupled to a LTQ-Orbitrap and samples were analysed in both positive and negative modes. The second dataset was generated using a HPLC coupled to a QTOF (quadrupole time-of-flight) mass spectrometer in positive mode.

Yet the same column and chromatographic conditions were used. This dataset was produced to screen variation of leaf specialized metabolites and is referred to as the “GWA dataset”. It includes representative peak intensities for 217 pseudo-molecules in 417 samples. Given our sampling and LC-MS protocols, the specialized metabolites investigated in this study were non-volatile and moderately polar.

#### Annotation of leaf specialized metabolites

Using the “annotation dataset”, which includes MS2 data in both positive and negative modes, we performed automatic annotation of specialized metabolites. We then matched molecules annotated in the “annotation dataset”, with the pseudo-molecules detected in the “GWA dataset” according to their retention time and *m/z*. We obtained putative annotations for 112 pseudo-molecules of the “GWA dataset” (out of 217). These putative annotations were grouped in ontology classes, and included mostly flavonoids, terpenes, quinic acids and derivatives, hydrolysable tannins, in addition to other rarer classes (Dataset S2).

### Variation of leaf specialized metabolites

Using the “GWA dataset”, we visualised leaf specialized metabolites variation of individuals using a principal component analysis (PCA) on the signal intensities of the representative peaks of 217 pseudo-molecules measured in 417 leaf samples, 206 and 211 for the low and high branches, respectively. PC1 explained 19.42% of total variance and clearly separated leaves sampled from high branches from leaves sampled from low branches (**Figure 2a**). We then performed a partial least-square discriminant analysis (PLS-DA) to discriminate between leaves from high and low branches. The first component resulting from this analysis explained 19.39% of the variance (Q2=0.59 with one component), confirming that branch height, which is related to leaf exposure to sunlight, is a major factor driving leaf content in specialized metabolites (**Figure 2d**).

**Figure 2.**
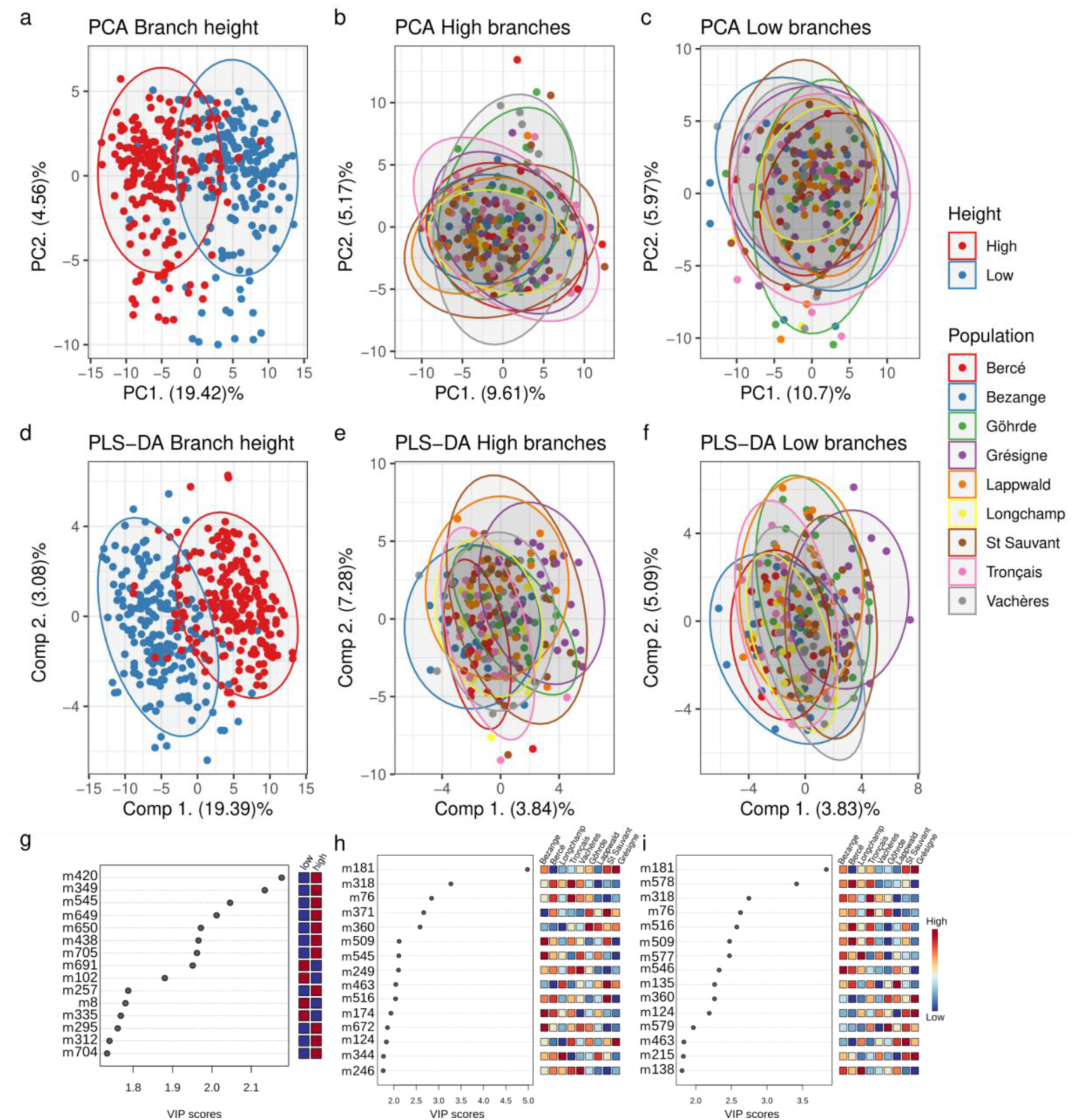
Variation of leaf specialized metabolites in oak trees from nine European populations. a-c. Principal component analysis of leaf specialized metabolites variation with 217 pseudo-molecules. **a.** Each point corresponds to a sample in our study (N=417) projected in the PC1 x PC2 plane (scores plot) of a principal component analysis of cube-root transformed and scaled (Pareto scaling) peak intensities of 217 pseudo-molecules. The points are coloured according to height (high and low) at which leaves were collected in the trees. **b.** Same as **a** but only includes samples collected from high branches (N=211) and coloured by populations. **c.** Same as **b** but only includes samples collected from low branches (N=206). **d-f.** Partial least-square discriminant analysis of leaf specialized metabolites variation with 217 pseudo-molecules. **d.** x- and y-axis correspond to the first and second components resulting from a discriminant analysis between high and low branches. **e.** The plot only includes samples collected on high tree branches (N=211), but x- and y-axis correspond to the first and second components resulting from a discriminant analysis between populations. **f.** Like **e.** But this time the plot only includes samples collected from low tree branches (N=206). In **a. to f.**, ellipses are coloured according to the grouping factor of interest (branch height in **a.** and **d.**, population in the others) and each include 95% of the samples in the corresponding group. **g, h and i:** Variable importance in the projections along component 1 of the discriminant analyses presented in **d**, **e** and **f**, respectively. For each plot, the x-axis represents the VIP (VIP scores > 1.5) of the 15 pseudo-molecules contributing the most to the component (y-axis). The coloured squares on the right of each plot represent the average transformed intensity value per pseudo-molecule, in each category of the grouping factor (branch heights in **g**, populations in **h** and **i**).

While the effect of branch height was evident from the PCA analysis, the population of origin of the trees did not have much influence on leaf specialized metabolite content (**Figure 2b,2c**). This observation is confirmed by the limited ability of the PLS-DA to discriminate between populations (Q2=0.07 and 0.04 with one component for high and low branches, respectively).

Indeed, the first component of the PLS-DA only explained 3.84% of the variance while the first component of the PCA explained 9.61%, confirming that the population of origin is not a major factor shaping specialized metabolite leaf content (**Figure 2d,2e**). While the first component of the PLS-DA discriminating populations did not explain much variation, it was correlated with the variation of intensity of the pseudo-molecule m181, which displayed the highest “variable importance in projection” (VIP) (**Figure 2h,2i**). Closer inspection of m181 variation showed that trees from the Grésigne population displayed, on average, higher concentrations (Figure **S4**).

## Genetic basis of leaf specialized metabolites

To investigate the genetic basis underlying the variation of leaf specialized metabolites in sessile oaks, we performed a genome-wide association study, using the “GWA dataset” and the farmCPU method (Liu *et al*. 2016). We performed association analysis separately for each of the 217 pseudo-molecules and for the two branch heights (217 molecules * 2 branch heights = 434 scans). All trees were genotyped for 1,408,029 SNPs (MAF>10% and a maximum genotype missingness at 10%) and we performed association analyses in panels of 200 and 203 trees for low and high branches, respectively (**Figure 3**). Considering the results from both branch heights together, we identified a total of 850 significant associations, corresponding to 752 individual SNPs associated with at least one of 93 pseudo-molecules (42% of pseudo-molecules).

**Figure 3.**
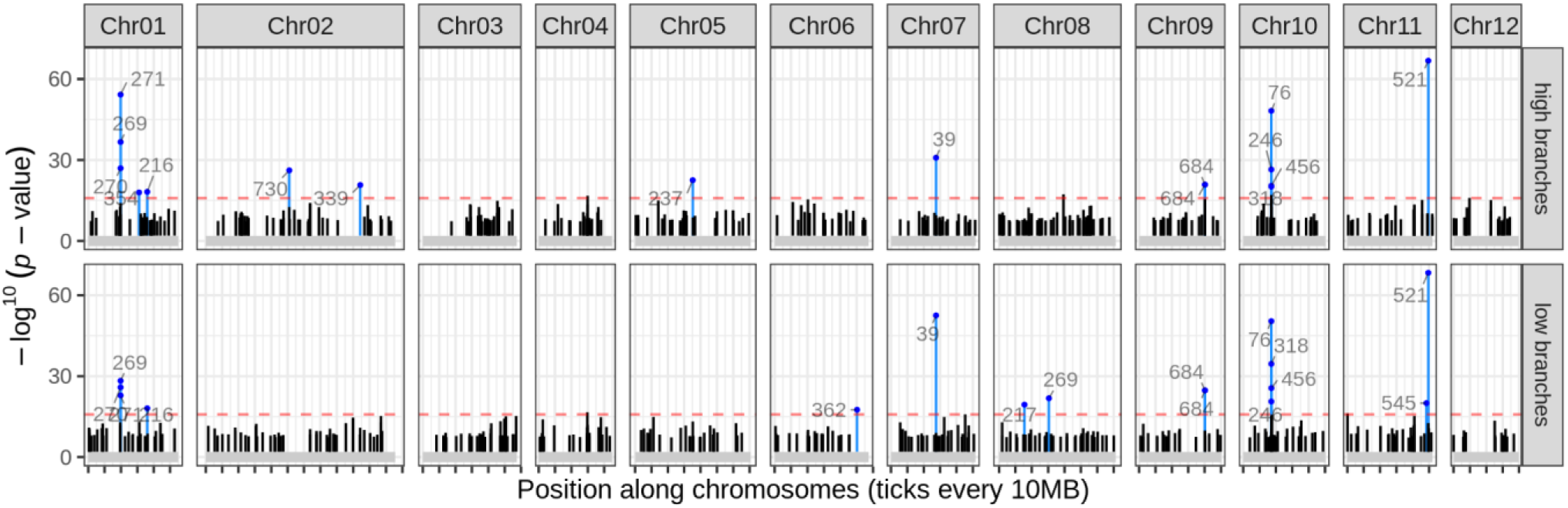
Genome-wide association analyses of oak leaf specialized metabolites. In all panels, the y-axis represents association scores of SNPs obtained by farmCPU and the x-axis positions along oak chromosomes. Each column of two panels corresponds to a chromosome (only chromosomes with significant associations are shown). The top and bottom rows of panels present associations detected using samples collected on high and low branches, respectively. Within each panel, vertical black bars correspond to significant associations detected by farmCPU. Blue bars correspond to the 5% most associated SNPs at both branch heights. These SNPs are labelled with the identifying number of the pseudo-molecule to which it is associated. The horizontal red dashed line corresponds to the 95% quantile of the distribution of association scores (-log^10^(p-*value*)).

Only a handful of pseudo-molecules displayed differentiation among populations (**Figure 2h,i**). To determine if this differentiation was larger than expected because of genetic drift and the demographic history of the species, we compared the F_ST_ of SNPs significantly associated with leaf specialized metabolites variation to the genome-wide F_ST_ distribution ^(^Weir & Cockerham 1984). Fifteen out of 752 markers associated with specialized metabolites variation displayed F_ST_ values above the 99th percentile of the genome-wide F_ST_ distribution (F_ST_ > 0.06). These 15 markers were spread over five different chromosomes and three scaffolds, and were associated with 17 pseudo-molecules, including m181 and m578, which were discriminant between populations (**Figure 2i,j**). The marker associated with specialized metabolite variation with the highest F_ST_ value (F_ST_=0.75) was associated with pseudo-molecule m181 (Qrob_H2.3_Sc0000298:373392), which discriminated the population Grésigne from the others (Figure **S3**).

To investigate the level of pleiotropy of regions associated with specialized metabolites, we clustered the 752 significant associations in 706 SNPs clusters separated by at least 3000 bases, a distance corresponding to the average genome-wide linkage disequilibrium (see Material and Methods). Of these SNPs clusters, 92% included associations with only one molecule. The remaining clusters included associations with two to six molecules. This analysis also allowed investigating the genetic architecture of the pseudo-molecules. We found that 14% of pseudo-molecules (13 out of 93) displayed significant associations in only one SNPs cluster and that on average, pseudo-molecules displayed associations in eight SNPs clusters.

We investigated candidate genes in windows of 3000 bp on each side of significantly associated SNPs. Overall significantly associated SNPs were located within or near 321 candidate genes (Dataset S3) annotated on the oak reference genomes (Plomion *et al*. 2018). Among them, 108 are annotated as long distance duplicated genes, 87 as tandem duplicates and 125 as singletons (Plomion *et al*. 2018, Supplementary Dataset 4).

Table 1 lists the five molecules for which we detected the strongest association score, along with their structural annotation and associated candidate genes. All other molecules with significant associations, their annotations and associated candidate genes when available, are listed in Dataset S2 & Dataset S3.

**Table 1.** The five pseudo-molecules with the strongest association, putative annotations and associated candidate genes.

| ID | Chemical Class | Putative Name | Chemical Formula | PubChem CID | MS-FINDER score | RT (min) | m/z<sup>+</sup> | Identification level | Chr: Position | Asso. score | *Q. robur* candidate gene | *A. thaliana* candidate gene | *A. thaliana* gene name |
| --- | --- | --- | --- | --- | --- | --- | --- | --- | --- | --- | --- | --- | --- |
| m521 |  |  |  |  |  | 5.51 | 598.1420 | Level 5 | Chr11: 49468544 | 68.32 |  | AT4G27570.1 | UGT79B3 |
| m412 |  |  |  |  |  | 6.32 | 477.2073 | Level 5 | Sc0000943: 46383 | 58.14 |  |  |  |
| m271 | Quinic acid and derivatives | Theogallin, 3-Galloylquinic acid | C<sub>14</sub>H<sub>16</sub>O<sub>10</sub> | 442988 | 6.5784 | 3.33 | 345.0803 | Level 2a | Chr01: 19760198 | 54.26 | Qrob_P0270670.2 | AT2G22990.6 | SNG1 |
| m39 | hydroxycinnamic acid like | Ferulic acid-like | C<sub>10</sub>H<sub>8</sub>O<sub>3</sub> | 445858 | NA | 5.05 | 177.0547 | Level 2b | Chr07: 27514866 | 52.49 |  | AT2G36290.1 | alpha/beta-Hydrolases protein superfamily |
| m76 |  |  |  |  |  | 5.46 | 211.0963 | Level 5 | Chr10: 17184779 | 50.38 | Qrob_P0166680.2 | AT5G61150.2 | VIP4 |

Based on structural annotations, the pseudo-molecules were assigned to a chemical class and given a putative name. The RT column gives the retention time in min, the *m/z*<sup>+</sup> column gives the mass to charge ratio (in positive mode) for the peak in the “GWA dataset” chosen as representative for the pseudo-molecule (which doesn’t always correspond to the parent ion). The identification level column gives confidence levels for annotations based on (Schymanski *et al.* 2014). The column “Chr:Position” gives the coordinates, on the *Q. robur* reference genome, of the genetic marker most strongly associated with the variation of each pseudo-molecule. The column “Asso. score” is the highest association score ( $-\log^{10}(p\text{-value})$ ) observed for the genetic marker across both branch heights using farmCPU. The columns “candidate gene” and “gene name” correspond to candidate genes identified within 3 kb of the associated marker, either on the *Quercus robur* reference genome, or based on sequence similarity with *Arabidopsis thaliana* proteins.

For the five molecules with the largest association scores, allelic variation at individual markers explained the bimodal nature of phenotypic distributions, confirming they were major effect loci (**Figure 4a**, Figure **S5**). Three additional observations can be made about these markers. First, alleles at these markers were present at balanced frequencies in all nine populations, hence their low F_ST_ values (**Figure 4a**). Second, allelic variation at these markers explained phenotypic variation similarly well in all nine populations (**Figure 4**). The third observation is that three out five, the homozygous genotype for the reference allele not present, or only represented by one individual, despite balanced allele frequencies (**Figure 4b**).

**Figure 4.**
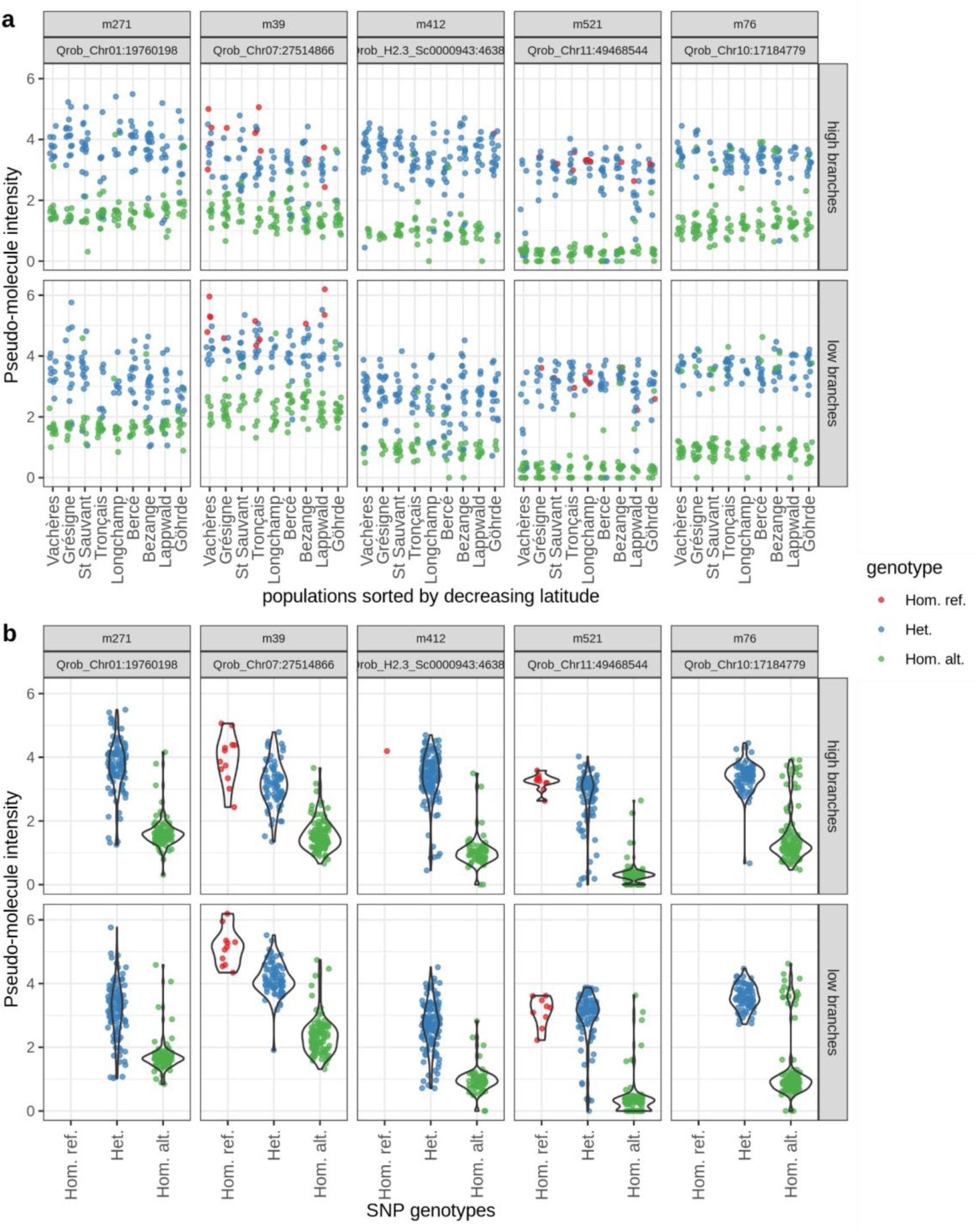
Variation of many leaf specialized metabolites can be explained by genetic variation at a single genetic marker. **a. and b.** In all panels, the plots represent the variation of the peak intensity (y-axis, cube-root transformation) for the five pseudo-molecules displaying to strongest associations with SNPs: m271, m39, m412, m521 and m76. The title of each panel includes the identifier of the pseudo-molecule and the position of the SNP most associated with its variation (Chr or Scaffold: position in bp). Each point corresponds to a leaf sample collected on high branches (top row of panels) or low branches (bottom row). Point colours correspond to the genotype of the tree at each SNP (see legend on the right-hand side). Note that in all panels, a small jitter was added to positions along the x-axis for clarity. In **a.** points are grouped by population of origin along the x-axis. In **b.** points are grouped by genotype at the SNP (all populations together). The labels along the x-axis (and in the legend) are: Hom. ref.: Homozygous for the reference allele; Het: Heterozygous; Hom. alt: Homozygous for the alternative allele. Violin plots were overlaid to help visualise the phenotypic distribution for each genotype.

For the five pseudo-molecules with the highest associations, we sought to check their automatic annotations by comparing MS^n^ spectra with public spectral libraries and by studying known fragmentation patterns (Sumner *et al*. 2007). Putative structures were proposed for only three of these molecules by the automated annotation pipeline (see Material and Methods). Of these, we were able to manually confirm the structures of two. The MS2 spectrum for m271 was consistent with it being a galloylquinic acid, a quinic and gallic acid derivative. Pseudo-molecule m39 was found to be a “ferulic acid-like” molecule, belonging to the ontology class of hydroxycinnamic acid (Table **1**).

SNPs associated with variation of these five molecules were located near candidate genes on the *Quercus robur* genome (Plomion *et al*. 2018). The region around the SNP most associated with the variation of pseudo-molecule m521 (Chr11:49468544), which is also the strongest association detected in our study, showed sequence similarity with the *Arabidopsis thaliana* gene AT4G27570, which encodes a UDP-glycosyltransferase (Li *et al*. 2017). The m521 was also significantly associated with SNPs in sixteen other SNP clusters.

For m271, m270 and m269 the strongest association was located in a region (Chr01:19760198) with a oak candidate gene (Qrob_P0270670.2) encoding a serine carboxypeptidase like protein that display sequence similarity with the *A. thaliana* gene AT2G22990 (SNG1), which encodes a sinapoyl transferase (Lehfeldt *et al*. 2000).

Interestingly, other associations for m271, but also m269 and m270, were located near the neighbouring gene (Chr01:19734409, Qrob_P0270680.2) with a sinapoyl glucose-malate O- sinapoyl transferase function that displayed sequence similarity with the same *A.thaliana* gene AT2G22990 (SNG1) (Dataset **S3**). These three pseudo-molecules were associated with 12 to 17 SNPs clusters, however, only the two associations mentioned above were common to all three pseudo-molecules.

For m39, the *Arabidopsis* candidate gene identified (AT2G36290.1) near the significant association (Chr07:27514866) was part of alpha/beta hydrolase proteins superfamily (Mindrebo, Nartey, Seto, Burkart & Noel 2016). This pseudo-molecule was also associated with SNPs in 11 SNPs clusters.

Finally, for m76 the candidate gene identified (Qrob_P0166680.2, Chr10:17184779) has sequence similarity with the *A. thaliana* gene VIP4 involved in vernalization response. The m76 was also associated with SNPs in 13 other clusters. No candidate genes were found near the strongest association for m412, and this pseudo-molecule was associated with SNPs in 13 other clusters.

We also investigated candidate genes around SNPs associated with the m181 pseudo- molecule. This pseudo-molecule was significantly associated with four markers and the strongest association was detected in scaffold 298 (position 373392 bp, -log^10^(*p-value*) = 11). Interestingly, the Grésigne population displayed homozygous reference genotype compared to the other populations that displayed homozygous alternative genotype which is consistent with the high F_ST_ value (0.75) of this marker (Figure **S4**). This marker was located in a region with sequence similarities with the *A. thaliana* gene ATMG00510, a mitochondrial gene encoding a NADH dehydrogenase. No chemical structural annotation was identified for this molecule.

## Discussion

### Oak populations display low genetic differentiation

We observed low genetic differentiation among the nine sessile oak populations, which is consistent with previously published observations, for the same populations, and in oaks more generally (Leroy *et al*. 2020; Saleh *et al*. 2022). The lack of differentiation among oak populations likely results from large population sizes, limiting genetic drift, the continuous distribution of the species and extensive long distance seed and pollen dispersal (Gerber *et al*. 2014).

Despite very low average genetic differentiation over the genomes, the large number of polymorphisms genotyped in our study allowed grouping individuals by population. Given the very low average differentiation we observe, this pattern is likely driven by a relatively small proportion of the genome (99% of the SNPs had a F_ST_ below 0.06). A previous study, investigated the variation of leaf phenology, growth or hydraulic traits in the same populations and identified phenotypic differentiation for phenology and growth traits. It is possible that differentiated loci detected here contribute to the variation of these traits and may be involved in the local adaptation of the nine populations to their respective environments (Torres-Ruiz *et al*. 2019).

### Many leaf specialized metabolites displayed extensive genetic variation within populations and low differentiation among sessile oak populations

We quantified the variation of 217 pseudo-molecules within the canopies of 206 and 211 for the low and high branches originating from nine populations. Our first observation was a large effect of branch height (**Figure 2a,d**). This effect impacted nearly all pseudo-molecules and explained a large fraction of the overall phenotypic variance across the 217 pseudo- molecules. Such variation was previously reported in pedunculate oak for total phenolics (Valdés-Correcher *et al*. 2020; Volf *et al*. 2022), suggesting that different environmental factors within the canopy such as the exposure to direct sunlight or to a different set of biotic interactions influences the production of leaf specialized metabolites.

A second observation was that oak populations display very little differentiation for leaf specialized metabolites when phenotyped in a common garden (**Figure 2a,b**) and patterns of variation for leaf specialized metabolites didn’t match patterns of genetic structure. Despite the lack of differences among populations, our genome-wide association analyses detected significant associations for 42% of the 217 specialized metabolites we investigated. This means that heritable variation exists for many molecules within oak populations. In addition, we found that these molecules were generally associated with a small number of loci, suggesting simple genetic architectures. This high heritability and the simple architecture of specialized metabolites variation is consistent with observations in other species such as rice (Matsuda *et al*. 2015) or *Arabidopsis thaliana* (Brachi *et al*. 2015). In addition, the lack of differentiation among populations is consistent with observations in natural oak stands (Bertić *et al*. 2021), where, within stand, variation of leaf specialized metabolites determines susceptibility to defoliation by *Tortrix viridana*.

Taken together, our results suggest that the variation of the vast majority of the leaf specialized metabolites we measured were not involved in the local adaptation of populations to their respective environments, at least at the geographical scale considered. This contrasts with the results of previous studies using common gardens to study leaf specialized metabolites variation, where patterns consistent with local adaptation in *Arabidopsis thaliana* (Brachi *et al*. 2015) or in trees (O’Reilly-Wapstra *et al*. 2013; Meijón *et al*. 2016) were reported. While using a randomised common garden trial allowed limiting the environmental effects on leaf specialized metabolites variation, it is possible that phenotypic traits that would display patterns of variation consistent with local adaptation were not expressed in our phenotyping conditions. This could have two plausible reasons. First, we explored the variation of leaf specialized metabolites between populations grown in a common garden at one time point during the summer. Sampling additional time points during the year, or quantifying the variation of leaf in a different location or year, may reveal more phenotypic variation among populations (Meijón *et al*. 2016). Second, the phenotypic variation observed within each population may not be representative of the variation in leaf specialized metabolites among adult trees in natural populations from which acorns were collected (Ducousso *et al*. 2022). In each population, acorns were collected, heat treated to avoid infection by pathogens and then raised in controlled conditions for 3 years. This protocol allowed to maximise seedling survival; however, it may have promoted the development of seedlings that would not have developed under natural conditions.

While most molecules did not display differentiation between populations, a discriminant analysis showed that pseudo-molecule m181 displayed higher signal intensities in the Grésigne population than in other populations (**Figure 2**). This pseudo-molecule was also strongly associated with a genetic marker that displayed the same pattern (Figure **S3**). While this result could be consistent with local adaptation, the associated marker was located on a short scaffold and the closest gene was similar to a mitochondrial gene in Arabidopsis, suggesting this region is part of the mitochondrial genome. A previous study of chloroplast and mitochondrial haplotypes in white oaks observed chloroplast or mitochondrial haplotypes exclusive populations (Dumolin, Demesure & Petit 1995). Hence it is possible that trees from the Grésigne population have a different mitochondrial haplotype than the other populations and that this variation is associated with phenotypic variation. Whether this variation is locally adaptive remains an open question.

Consistent with the lack of phenotypic differentiation among populations, the vast majority of markers associated with leaf specialized metabolites did not display high F_ST_ values (except for 17 molecules, including the m181 pseudo-molecule). One interpretation could be that the variation of many leaf specialized metabolites evolves neutrally. This is unlikely in our opinion as drift would likely generate differentiation between populations, as visible in our analysis of population genetic variation, and leaf specialized metabolites are known to impact fitness related traits such as stress resistance and defence against pests (De-la-Cruz, Merilä, Valverde, Flores-Ortiz & Núñez-Farfán 2020). At least two scenarios could be formulated to explain the very high level of variation within provenances for leaf specialized metabolites, or even the balanced frequency of phenotype classes we observed for certain compounds across all oak populations (see Moore, Andrew, Külheim & Foley 2014 for a review). A first scenario could be that phenotypic variation among adult trees within populations of origin is geographically structured at a fine scale, due to adaptation to micro-local conditions. Locally this would result in a diverse pollen cloud and extensive phenotypic variation among offsprings. This scenario would likely require strong and recurrent selection on seedling populations that vary over relatively small distances (maybe in order of a few kilometres, Gerber *et al*. 2014) to geographically structure the phenotypic variation in neighbouring adult populations. The second scenario would be analogous to the first one, but instead of varying over the landscape, selection would vary in time, favouring different phenotypes depending on the year. Investigating allele frequencies at associated markers in trees of different ages could be a way to test this scenario (Saleh *et al*. 2022). Moreover, selective trade-off may act in combination with these two scenarios to maintain balanced frequencies of phenotypic classes and alleles (Troth, Puzey, Kim, Willis & Kelly 2018). This was observed in a recent study exploring the mechanisms that maintain high levels of variation in leaf glucosinolates in a perennial wildflower (*Boechera stricta*). The authors showed that the chemical profile of plants had contrasting fitness effects depending on the presence of herbivores and water availability. Because these selective pressures varied over time and small geographical scales, selection by herbivores and exposure to drought have maintained genetic variation for glucosinolate profiles for a very long time (Carley *et al*. 2021). We suspect similar mechanisms may be involved in maintaining high levels of within populations variation for leaf specialized metabolites in oaks, as individual specialized metabolites often play a key role in the response to multiple stresses. Like in *Boechera stricta*, it is possible that selective pressures have opposite effects on leaf specialized metabolites variation in oak populations and vary across space or time, preventing the fixation of a beneficial phenotypic form. In the next section, we discuss molecules that displayed high levels of within populations genetic variation and the associated candidate genes, with a focus on potential pleiotropic effects.

### Potential pleiotropic effect of oak leaf specialized metabolites associated genes

In oaks, previous studies on leaf specialized metabolites abundance variation showed that variation of different specialized metabolites, such as quinic acids or quercitols, were shaped by both biotic stresses (Sardans *et al*. 2014; Bertić *et al*. 2021) and abiotic stresses (Passarinho, Lamosa, Baeta, Santos & Ricardo 2006; Aranda, Cadahía & Fernández De Simón 2021). As hypothesised in the previous section, selective pressures shaping leaf specialized metabolite variation can vary over time and space, maintaining genetic variation at the underlying biosynthetic genes or transcription factors (Carley *et al*. 2021).

Here, we only discuss possible pleiotropic effects of two metabolites for which we have good confidence in their putative annotations, identified a relevant candidate gene, and for which literature searches revealed potential pleiotropic effects on the tolerance or resistance to both abiotic and biotic stresses.

The first pseudo-molecule of interest, m39, was annotated as a ferulic acid like metabolite, derived from the phenylpropanoid pathway, and associated with a single major-effect locus (chromosome 7; position 27,514,866 bp) with a sequence similarity in *Arabidopsis* with an alpha/beta-Hydrolases protein superfamily. The alpha/beta-Hydrolases protein superfamily was previously described to be involved in specialized metabolism in plants (reviewed in Mindrebo *et al*. 2016). Ferulic acids play a key role in plant cell wall biosynthesis and may be involved in drought tolerance in cereals (Hura, Hura & Grzesiak 2009). In addition, ferulic acids may contribute to herbivore resistance in oaks. Their absolute quantity in leaves, quantified as lignin equivalent, were positively correlated with the growth rate of caterpillars of a generalist herbivore, *Lymantria dispar* (Damestoy *et al*. 2019).

The second pseudo-molecule, m271, is part of a group of three also including m269 and m270. The three pseudo-molecules m269, m270, and m271 did not have the same retention time, but had identical *m/z*, which together suggests they were isomers. Our automatic annotations and inspection of fragmentation patterns suggests they were likely galloylquinic acid, also known as theogallin.

In oaks, galloylquinic acids were previously identified within leaves of 12 species of black and white oaks (Yarnes, Boecklen, Tuominen & Salminen 2006), but, to our knowledge, no study investigated the biological function of this molecule in oaks. Galloylquinic acid is composed of a quinic acid and a gallic acid moiety and is a precursor of hydrolysable tannins. Previous studies have investigated the properties of quinic acid and gallic acid in oaks, however separately. Quinic acid concentrations were shown to increase in response to wounding (Sardans *et al*. 2014), temperature elevation (Passarinho *et al*. 2006) and water deprivation (Aranda *et al*. 2021). In addition, the production of gallic acid was frequently associated with herbivore resistance, and its concentration appears to vary seasonally in oak leaves (Salminen *et al*. 2004).

All three pseudo-molecules were associated with partially different sets of markers across the genomes but all displayed associations in a region of chromosome 1, including a marker to which they were all very strongly associated, at position 19,760,198 bp. This marker is located in a gene (Qrob_P0270670.2) annotated as a serine carboxypeptidase like (SCPL) on the oak reference genome. SCPL genes were previously shown to play a key role in metabolomic biosynthesis pathways of flavonoids in grapevine *Vitis vinifera* (Bontpart *et al*. 2018) and galloylated catechins in *Camellia sinensis* (Ahmad *et al*. 2020). Other associations with these three pseudo-molecules in this region were located near the neighbouring gene (Qrob_P0270680.2) encoding a sinapoyl glucose-malate-O-sinapoyl transferase, involved in phenylpropanoid biosynthesis pathway (Vogt 2010).

Interestingly, both candidate genes, Qrob_P0270670.2 and Qrob_P0270680.2, were identified as tandem duplicates (Plomion *et al*. 2018), and matched with the same *Arabidopsis thaliana* gene: sinapoyl glucose 1 (SNG1). The fact that these two genes were identified as tandem duplicates may explain why the reference homozygous genotype was absent for the SNP with the strongest association to the variation of these pseudo-molecules (Qrob_Chr1:19760198). The SNP may tag a copy number polymorphism in this region. Further investigations will be needed to confirm copy number variation and its role in the variation of these molecules in oaks (Lichman, Godden & Buell 2020).

In conclusion, we observed an extensive variation of leaf specialized metabolites within sessile oak populations sampled over a latitudinal cline and established in a common garden. Very few molecules displayed differentiation among populations, suggesting little population effects, however we found significant associations for 42% of the leaf specialized metabolites investigated suggesting that the variation of many molecules within populations is genetically determined. The lack of population differentiation and high heritability within all populations suggest that most leaf specialized metabolites were not involved in the local adaptation of populations to their respective environment. Instead, we found very high levels of genetic variation for leaf specialized metabolites present in all nine European populations investigated. While this variation could be mostly selectively neutral, it could also be maintained by a form of selective trade-off where the production of a molecule is favourable under certain environmental conditions, but deleterious in others. The annotation of pseudo- molecules and candidate genes detected by our genome-wide association analyses were discussed in the light of this hypothesis.

## Supporting information

Supplementary informations

## Acknowledgments

This work was funded by successive grants to C.P. and B.B. between (2016-2019) from the Idex of the university of Bordeaux, Action Thématique Transversale: <u>METAB-OAK</u> “Natural variation of secondary metabolite production in forest trees: the case of European white oaks”. B.B. has received the support of the European Union in the framework of the Marie-Curie FP7 COFUND People Programme, through the award of an AgreenSkills/AgreenSkills+ fellowship (under Grant Agreement 267196). This work was also made possible by MetaboHUB (ANR-11-INBS-0010). D.CE. received funding for a PhD from the INRAE department ECODIV and Clermont Auvergne Métropole. Computations and data storage for this work were provided by the Bordeaux Bioinformatics Center at the University of Bordeaux (CBIB, https://www.cbib.u-bordeaux.fr), the Genotoul bioinformatics platform in Toulouse, Occitanie, France (Bioinfo Genotoul, http://bioinfo.genotoul.fr), and Le Mésocentre de Calcul Intensif Aquitain (MCIA), France.

## Author contributions

B.B. and C.P. designed the study, G.LP., F.B., D.CE. and B.B. collected samples, B.B., D.CE., and C.L. extracted DNA and performed quality controls. B.B., S.B. and A.M. designed and optimised LC-MS methods. B.B., S.B. and D.CE. performed LC-MS analysis. A.K. and A.D. installed the oak common garden and contributed information about provenances and its experimental design. B.B. and D.CE. performed the analyses with guidance from S.B., wrote the manuscript. C.P., A.M., S.B., A.K. provided comments on the manuscript.

## Data Accessibility and Benefit-Sharing

Codes and pipelines used for this study will be available upon publication at:

1. https://forgemia.inra.fr/domitille.coq-etchegaray/gw-oak-snp-pipeline.git
2. https://forgemia.inra.fr/domitille.coq-etchegaray/gwas-gwoak-metaboak-pipeline.git
3. https://forgemia.inra.fr/domitille.coq-etchegaray/metab-oak-pipeline.git

Raw metabolomics data and SNPs matrix will be available upon publication in the Dataverse INRAE Biogeco: doi:10.57745/3CSJZ9

Raw WGS sequencing used for this study will be available upon publication in the European Nucleotide Archive (ENA) at EMBL-EBI under accession number PRJEB60751. (https://www.ebi.ac.uk/ena/browser/view/PRJEB60751).

## Supporting Information

**Table S1** Geographic origin of the populations and number of oak trees analysed per population.

**Table S2** Tables of oak trees and sample counts for each analysis in this study.

**Table S3** Optimised parameters used for peak picking and peak alignment for metabolomics data treatment with the R package XCMS.

**Figure S1** Relationships between the datasets produced

**Figure S2** Potential confounding effects on specialized metabolites variation in our study.

**Figure S3** Decay of linkage disequilibrium in our sample of 225 *Quercus petraea* from 9 European populations.

**Figure S4** The variation of leaf specialized metabolite m181 among populations is largely explained by genetic variation at a single genetic marker.

**Figure S5** Phenotypic variation explained by genetic variation at the most associated marker for each pseudo-molecule.

**Dataset S1** Pseudo-molecules intensities, sampling and peak descriptions.

**Dataset S2** Formula and automatic structural annotation of pseudo-molecules based on MS2 acquisitions.

**Dataset S3** GWAs associations results

## References

1. Ahmad M.Z., Li P., She G., Xia E., Benedito V.A., Wan X.C. & Zhao J. (2020) Genome- Wide Analysis of Serine Carboxypeptidase-Like Acyltransferase Gene Family for Evolution and Characterization of Enzymes Involved in the Biosynthesis of Galloylated Catechins in the Tea Plant (*Camellia sinensis*). Frontiers in plant science 11, 848.

2. Aranda I., Cadahía E. & Fernández De Simón B. (2021) Specific leaf metabolic changes that underlie adjustment of osmotic potential in response to drought by four *Quercus* species. Tree physiology 41, 728–743.

3. Bailey J.K., Deckert R., Schweitzer J.A., Rehill B.J., Lindroth R.L., Gehring C. & Whitham T.G. (2005) Host plant genetics affect hidden ecological players: links among *Populus*, condensed tannins, and fungal endophyte infection. Canadian journal of botany. 83, 356–361.

4. Bartholomé J., Brachi B., Marçais B., Mougou-Hamdane A., Bodénès C., Plomion C., … Desprez-Loustau M.L. (2020) The genetics of exapted resistance to two exotic pathogens in pedunculate oak. The New phytologist 226, 1088–1103.

5. Bednarek P. (2012) Chemical warfare or modulators of defence responses-the function of secondary metabolites in plant immunity This review comes from a themed issue on Biotic interactions. Current opinion in plant biology 15, 407–414.

6. Berardini T.Z., Reiser L., Li D., Mezheritsky Y., Muller R., Strait E. & Huala E. (2015) The arabidopsis information resource: Making and mining the “gold standard” annotated reference plant genome. Genesis 53, 474–485.

7. Bertić M., Schroeder H., Kersten B., Fladung M., Orgel F., Buegger F., … Ghirardo A. (2021) European oak chemical diversity – from ecotypes to herbivore resistance. The New phytologist 232, 818–834.

8. Bhatia G., Patterson N., Sankararaman S. & Price A.L. (2013) Estimating and interpreting FST: The impact of rare variants. Genome research 23, 1514.

9. Bontpart T., Ferrero M., Khater F., Marlin T., Vialet S., Vallverdù-Queralt A., … Terrier N. (2018) Focus on putative serine carboxypeptidase-like acyltransferases in grapevine. Plant physiology and biochemistry 130, 356–366.

10. Brachi B., Meyer C.G., Villoutreix R., Platt A., Morton T.C., Roux F. & Bergelson J. (2015) Coselected genes determine adaptive variation in herbivore resistance throughout the native range of *Arabidopsis thaliana*. Proceedings of the National Academy of Sciences of the United States of America 112, 201421416.

11. Camacho C., Coulouris G., Avagyan V., Ma N., Papadopoulos J., Bealer K. & Madden T.L. (2009) BLAST+: Architecture and applications. BMC bioinformatics 10, 421.

12. Carley L.N., Mojica J.P., Wang B., Chen C.-Y., Lin Y.-P., Prasad K.V.S.K., … Mitchell- Olds T. (2021) Ecological factors influence balancing selection on leaf chemical profiles of a wildflower. Nature ecology & evolution 5, 1135–1144.

13. Chambers M.C., MacLean B., Burke R., Amodei D., Ruderman D.L., Neumann S., … Mallick P. (2012) A cross-platform toolkit for mass spectrometry and proteomics. Nature biotechnology 30, 918–920.

14. Chang C.C., Chow C.C., Tellier L.C.A.M., Vattikuti S., Purcell S.M. & Lee J.J. (2015) Second-generation PLINK: Rising to the challenge of larger and richer datasets. GigaScience 4, 7.

15. Chong J. & Xia J. (2018) MetaboAnalystR: an R package for flexible and reproducible analysis of metabolomics data. Bioinformatics 34, 4313–4314.

16. Damestoy T., Brachi B., Moreira X., Jactel H., Plomion C. & Castagneyrol B. (2019) Oak genotype and phenolic compounds differently affect the performance of two insect herbivores with contrasting diet breadth. Tree physiology 39, 615–627.

17. Danecek P., Bonfield J.K., Liddle J., Marshall J., Ohan V., Pollard M.O., … Li H. (2021) Twelve years of SAMtools and BCFtools. GigaScience 10, 1–4.

18. Dearing M.D., Foley W.J. & McLean S. (2005) The Influence of Plant Secondary Metabolites on the Nutritional Ecology of Herbivorous Terrestrial Vertebrates. Annual review of ecology, evolution, and systematics 36, 169–189.

19. De-la-Cruz I.M., Merilä J., Valverde P.L., Flores-Ortiz C.M. & Núñez-Farfán J. (2020) Genomic and chemical evidence for local adaptation in resistance to different herbivores in *Datura stramonium*. Evolution; international journal of organic evolution 74, 2629– 2643.

20. De Luca V. & St Pierre B. (2000) The cell and developmental biology of alkaloid biosynthesis. Trends in plant science 5, 168–173.

21. Ducousso A., Ehrenmann F., Girard Q., Lamy J.B., Louvet J.M., Reynet P., … Kremer A. (2022) Long-term and large-scale *Quercus petraea* population survey conducted in provenance tests installed in France. Annals of forest science 79, 1–10.

22. Dumolin S., Demesure B. & Petit R.J. (1995) Inheritance of chloroplast and mitochondrial genomes in pedunculate oak investigated with an efficient PCR method. *TAG*. Theoretical and applied genetics 91, 1253–1256.

23. Forkner R.E., Marquis R.J. & Lill J.T. (2004) Feeny revisited: condensed tannins as anti-herbivore defences in leaf-chewing herbivore communities of *Quercus*. Ecological entomology 29, 174–187.

24. Fraisier-Vannier O., Chervin J., Cabanac G., Puech V., Fournier S., Durand V., … Marti G. (2020) MS-CleanR: A Feature-Filtering Workflow for Untargeted LC-MS Based Metabolomics. Analytical chemistry 92, 9971–9981.

25. Galeotti F., Barile E., Curir P., Dolci M. & Lanzotti V. (2008) Flavonoids from carnation (*Dianthus caryophyllus*) and their antifungal activity. Phytochemistry letters 1, 44–48.

26. Gerber S., Chadoeuf J., Gugerli F., Lascoux M., Buiteveld J., Cottrell J., … Kremer A. (2014) High rates of gene flow by pollen and seed in oak populations across Europe. PloS one 9.

27. Hura T., Hura K. & Grzesiak S. (2009) Possible contribution of cell-wall-bound ferulic acid in drought resistance and recovery in triticale seedlings. Journal of plant physiology 166, 1720–1733.

28. Joshi N.A. & Fass J.N. (2011) Sickle: a sliding-window, adaptive, quality-based trimming tool for FastQ files.

29. Jun G., Wing M.K., Abecasis G.R. & Kang H.M. (2015) An efficient and scalable analysis framework for variant extraction and refinement from population scale DNA sequence data. Genome research 25, gr.176552.114.

30. Köster J., Mölder F., Jablonski K.P., Letcher B., Hall M.B., Tomkins-Tinch C.H., … Nahnsen S. (2021) Sustainable data analysis with Snakemake. F1000Research 2021 10:33 10, 33.

31. Kuhl C., Tautenhahn R., Böttcher C., Larson T.R. & Neumann S. (2012) CAMERA: An Integrated Strategy for Compound Spectra Extraction and Annotation of Liquid Chromatography/Mass Spectrometry Data Sets. Analytical chemistry 84, 283–289.

32. Kurtzer G.M., Sochat V. & Bauer M.W. (2017) Singularity: Scientific containers for mobility of compute. PloS one 12, e0177459.

33. Lehfeldt C., Shirley A.M., Meyer K., Ruegger M.O., Cusumano J.C., Viitanen P.V., … Chapple C. (2000) Cloning of the SNG1 gene of Arabidopsis reveals a role for a serine carboxypeptidase-like protein as an acyltransferase in secondary metabolism. The Plant cell 12, 1295–1306.

34. Leroy T., Louvet J.M., Lalanne C., Le Provost G., Labadie K., Aury J.M., … Kremer A. (2020) Adaptive introgression as a driver of local adaptation to climate in European white oaks. The New phytologist 226, 1171–1182.

35. Libiseller G., Dvorzak M., Kleb U., Gander E., Eisenberg T., Madeo F., … Magnes C. (2015) IPO: A tool for automated optimization of XCMS parameters. BMC bioinformatics 16, 1–10.

36. Lichman B.R., Godden G.T. & Buell C.R. (2020) Gene and genome duplications in the evolution of chemodiversity: perspectives from studies of Lamiaceae. Current Opinion in Plant Biology 55, 74–83.

37. Li H. (2013) Aligning sequence reads, clone sequences and assembly contigs with BWA-MEM. arXiv preprint arXiv 00, 3.

38. Li P., Li Y.-J., Zhang F.-J., Zhang G.-Z., Jiang X.-Y., Yu H.-M. & Hou B.-K. (2017) The Arabidopsis UDP-glycosyltransferases UGT79B2 and UGT79B3, contribute to cold, salt and drought stress tolerance via modulating anthocyanin accumulation. The Plant journal: for cell and molecular biology 89, 85–103.

39. Liu X., Huang M., Fan B., Buckler E.S. & Zhang Z. (2016) Iterative Usage of Fixed and Random Effect Models for Powerful and Efficient Genome-Wide Association Studies. PLoS genetics 12, e1005767.

40. Martin M. (2011) Cutadapt removes adapter sequences from high-throughput sequencing reads. *EMBnet*. journal 17, 10–12.

41. Maruki T. & Lynch M. (2015) Genotype-frequency estimation from high-throughput sequencing data. Genetics 201, 473–486.

42. Maruki T. & Lynch M. (2017) Genotype calling from population-genomic sequencing data. G3: Genes, Genomes, Genetics 7, 1393–1404.

43. Matsuda F., Nakabayashi R., Yang Z., Okazaki Y., Yonemaru J.-I., Ebana K., … Saito K. (2015) Metabolome-genome-wide association study dissects genetic architecture for generating natural variation in rice secondary metabolism. The Plant journal: for cell and molecular biology 81, 13–23.

44. McCormick A.C., Unsicker S.B. & Gershenzon J. (2012) The specificity of herbivore-induced plant volatiles in attracting herbivore enemies. Trends in plant science 17, 303– 310.

45. Meijón M., Feito I., Oravec M., Delatorre C., Weckwerth W., Majada J. & Valledor L. (2016) Exploring natural variation of *Pinus pinaster* Aiton using metabolomics: Is it possible to identify the region of origin of a pine from its metabolites? Molecular ecology 25, 959–976.

46. Mindrebo J.T., Nartey C.M., Seto Y., Burkart M.D. & Noel J.P. (2016) Unveiling the functional diversity of the alpha/beta hydrolase superfamily in the plant kingdom. Current opinion in structural biology 41, 233–246.

47. Mitchell R.J., Bellamy P.E., Ellis C.J., Hewison R.L., Hodgetts N.G., Iason G.R., … Taylor A.F.S. (2019) Collapsing foundations: The ecology of the British oak, implications of its decline and mitigation options. Biological conservation 233, 316–327.

48. Moore B.D., Andrew R.L., Külheim C. & Foley W.J. (2014) Explaining intraspecific diversity in plant secondary metabolites in an ecological context. The New phytologist 201, 733–750.

49. Nakabayashi R., Yonekura-Sakakibara K., Urano K., Suzuki M., Yamada Y., Nishizawa T., … Saito K. (2014) Enhancement of oxidative and drought tolerance in Arabidopsis by overaccumulation of antioxidant flavonoids. The Plant journal: for cell and molecular biology 77, 367–379.

50. Newcombe G. (1998) A review of exapted resistance to diseases of *Populus*. European journal of forest pathology. 28, 209–216.

51. Obata T., Witt S., Lisec J., Palacios-Rojas N., Florez-Sarasa I., Yousfi S., … Fernie A.R. (2015) Metabolite Profiles of Maize Leaves in Drought, Heat, and Combined Stress Field Trials Reveal the Relationship between Metabolism and Grain Yield. Plant physiology 169, 2665–2683.

52. O’Reilly-Wapstra J.M., Miller A.M., Hamilton M.G., Williams D., Glancy-Dean N. & Potts B.M. (2013) Chemical Variation in a Dominant Tree Species: Population Divergence, Selection and Genetic Stability across Environments. PloS one 8, 58416.

53. Passarinho J.A.P., Lamosa P., Baeta J.P., Santos H. & Ricardo C.P.P. (2006) Annual changes in the concentration of minerals and organic compounds of *Quercus suber* leaves. Physiologia plantarum 127, 100–110.

54. Plomion C., Aury J.-M., Amselem J., Leroy T., Murat F., Duplessis S., … Salse J. (2018) Oak genome reveals facets of long lifespan. Nature Plants 4, 440–452.

55. R Core Team (2021) R: A Language and Environment for Statistical Computing. R Foundation for Statistical Computing, Vienna, Austria.

56. Rozema J., Björn L.O., Bornman J.F., Gaberščik A., Häder D.P., Trošt T., … Meijkamp B.B. (2002) The role of UV-B radiation in aquatic and terrestrial ecosystems-An experimental and functional analysis of the evolution of UV-absorbing compounds. Journal of photochemistry and photobiology. B, Biology 66, 2–12.

57. Sáenz-Romero C., Lamy J.B., Ducousso A., Musch B., Ehrenmann F., Delzon S., … Kremer A. (2017) Adaptive and plastic responses of *Quercus petraea* populations to climate across Europe. Global change biology 23, 2831–2847.

58. Saleh D., Chen J., Leplé J.C., Leroy T., Truffaut L., Dencausse B., … Kremer A. (2022) Genome-wide evolutionary response of European oaks during the Anthropocene. Evolution Letters 6, 4–20.

59. Salminen J.-P., Roslin T., Karonen M., Sinkkonen J., Pihlaja K. & Pulkkinen P. (2004) Seasonal variation in the content of hydrolyzable tannins, flavonoid glycosides, and proanthocyanidins in oak leaves. Journal of chemical ecology 30, 1693–1711.

60. Sardans J., Gargallo-Garriga A., Pérez-Trujillo M., Parella T.J., Seco R., Filella I. & Peñuelas J. (2014) Metabolic responses of *Quercus ilex* seedlings to wounding analysed with nuclear magnetic resonance profiling. Plant biology 16, 395–403.

61. Sardans J., Peñuelas J. & Rivas-Ubach A. (2011) Ecological metabolomics: Overview of current developments and future challenges. Chemoecology 21, 191–225.

62. Schweitzer J.A., Bailey J.K., Fischer D.G., LeRoy C.J., Lonsdorf E.V., Whitham T.G. & Hart S.C. (2008) Plant–Soil–Microorganism Interactions: Heritable Relationship Between Plant Genotype and Associated Soil Microorganisms. Ecology 89, 773–781.

63. Schymanski E.L., Jeon J., Gulde R., Fenner K., Ruff M., Singer H.P. & Hollender J. (2014) Identifying small molecules via high resolution mass spectrometry: communicating confidence. Environmental science & technology 48, 2097–2098.

64. Sebastiana M., Gargallo-Garriga A., Sardans J., Pérez-Trujillo M., Monteiro F., Figueiredo A., … Peñuelas &. J. (2021) Metabolomics and transcriptomics to decipher molecular mechanisms underlying ectomycorrhizal root colonization of an oak tree. Scientific reports 11, 8576.

65. Smith C. a., Want E.J., O’Maille G., Abagyan R. & Siuzdak G. (2006) XCMS: Processing Mass Spectrometry Data for Metabolite Profiling Using Nonlinear Peak Alignment, Matching, and Identification. Analytical chemistry 78, 779–787.

66. Sumner L.W., Amberg A., Barrett D., Beale M.H., Beger R., Daykin C.A., … Viant M.R. (2007) Proposed minimum reporting standards for chemical analysis Chemical Analysis Working Group (CAWG) Metabolomics Standards Initiative (MSI). Metabolomics: Official journal of the Metabolomic Society 3, 211–221.

67. Torres-Ruiz J.M., Kremer A., Carins Murphy M.R., Brodribb T., Lamarque L.J., Truffaut L., … Delzon S. (2019) Genetic differentiation in functional traits among European sessile oak populations. Tree physiology 39, 1736–1749.

68. Troth A., Puzey J.R., Kim R.S., Willis J.H. & Kelly J.K. (2018) Selective trade-offs maintain alleles underpinning complex trait variation in plants. Science 361, 475–478.

69. Tsugawa H., Cajka T., Kind T., Ma Y., Higgins B., Ikeda K., … Arita M. (2015) MS-DIAL: data-independent MS/MS deconvolution for comprehensive metabolome analysis. Nature Methods 2015 *12*: 6 12, 523–526.

70. Tsugawa H., Kind T., Nakabayashi R., Yukihira D., Tanaka W., Cajka T., … Arita M. (2016) Hydrogen Rearrangement Rules: Computational MS/MS Fragmentation and Structure Elucidation Using MS-FINDER Software. Analytical chemistry 88, 7946–7958.

71. Valdés-Correcher E., Bourdin A., González-Martínez S.C., Moreira X., Galmán A., Castagneyrol B. & Hampe A. (2020) Leaf chemical defences and insect herbivory in oak: accounting for canopy position unravels marked genetic relatedness effects. Annals of botany 126, 865–872.

72. Van der Auwera G.A., Carneiro M.O., Hartl C., Poplin R., del Angel G., Levy-Moonshine A., … DePristo M.A. (2013) From fastQ data to high-confidence variant calls: The genome analysis toolkit best practices pipeline. Current protocols in bioinformatics 11, 11.10.1.

73. Vargas P., Farias G. a., Nogales J., Prada H., Carvajal V., Barón M., … Gallegos M.-T. (2013) Plant flavonoids target *Pseudomonas syringae pv.* tomato DC3000 flagella and type III secretion system. Environmental microbiology reports 5, 841–850.

74. Vogt T. (2010) Phenylpropanoid biosynthesis. Molecular plant 3, 2–20.

75. Volf M., Volfová T., Seifert C.L., Ludwig A., Engelmann R.A., Jorge L.R. é., … van Dam N.M. (2022) A mosaic of induced and non-induced branches promotes variation in leaf traits, predation and insect herbivore assemblages in canopy trees. Ecology letters 25, 729–739.

76. Walker T.S., Bais H.P., Grotewold E. & Vivanco J.M. (2003) Root exudation and rhizosphere biology. Plant physiology 132, 44–51.

77. Weir B.S. & Cockerham C.C. (1984) Estimating F-Statistics for the Analysis of Population Structure. Evolution; international journal of organic evolution 38, 1358–1370.

78. Wink M. (2018) Plant Secondary Metabolites Modulate Insect Behavior-Steps Toward Addiction? Frontiers in physiology 9.

79. Yarnes C.T., Boecklen W.J., Tuominen K. & Salminen J.P. (2006) Defining phytochemical phenotypes: Size and shape analysis of phenolic compounds in oaks (Fagaceae, Quercus) of the Chihuahuan Desert. Canadian journal of botany 84, 1233–1248.

80. Yin L., Zhang H., Tang Z., Xu J., Yin D., Zhang Z., … Liu X. (2021) rMVP: A Memory-efficient, Visualization-enhanced, and Parallel-accelerated Tool for Genome-wide Association Study. Genomics, proteomics & bioinformatics 19, 619–628.

81. Züst T., Heichinger C., Grossniklaus U., Harrington R., Kliebenstein D.J. & Turnbull L.A. (2012) Natural Enemies Drive Geographic Variation in Plant Defenses. Science 338, 116–119.

