## Supplementary informations for "Extensive genetic variation of leaf specialized metabolites in sessile oak (*Quercus petraea*) populations"

Short running title: Genetic bases of leaf specialized metabolites in oak

Domitille Coq--Etchegaray[1*], Stéphane Bernillon[2, 3], Grégoire Le-Provost[1], Antoine Kremer[1], Alexis Ducousso[1], Céline Lalanne[1], Fabrice Bonne[4], Annick Moing[3], Christophe Plomion[1], Benjamin Brachi[1*]

[1] Univ. Bordeaux, INRAE, BIOGECO, UMR 1202, F-33610 Cestas, France

[2] INRAE, MycSA, F-33140 Villenave d'Ornon, France

[3] Univ. Bordeaux, INRAE, BFP, UMR 1332, Bordeaux Metabolome, MetaboHUB, F-33140 Villenave d'Ornon, France

[4] Univ. Lorraine, AgroParisTech, INRAE, UMR SILVA, F-54280 Champenoux, France

**Corresponding authors:**

Benjamin Brachi, INRAE, 69 route d'Arcachon, Cestas, France

### Supplementary Materials and Methods

#### S1. Samples lyophilization

We ground frozen leaves to a thin powder by adding two 8-mm stainless steel ball bearings to each sample and shaking the vials with a GenoGrinder 2010 (SPEX SamplePrep, Metuchen, NJ, U.S.) equipped with a custom tube holder and set to 1750 rpm for 2*45 seconds. We then lyophilised the leaf powder using a Cryotec freeze dryer (Cryotec, Saint-Gély-du-Fesc, France) and a two stages lyophilisation program:

- primary lyophilisation: pressure set to 0.6 mBar (6e-5 MPa) with shelves temperature set to -60ºC for 24 h, -50ºC for 24 h, -40ºC for 24 h, and -20ºC for 18 h
- secondary lyophilisation: shelves at -20ºC, 0.1 mBar (1e-5 MPa) 30 min followed by 0.001 mBar for 1 h, repeated 4 times.

2

Lyophilized samples were sealed and stored in sealed bags on silica-gel at -20°C until further processing.

### S2. DNA extraction

We extracted DNA from 10 mg lyophilized leaf powder, from each low and high sample per tree, for 225 individuals sampled in 2016 using a custom CTAB extraction protocol (Larue *et al.*, 2021). We estimated DNA quality and concentration using a NanoDrop™ 8000 (Thermo Fisher, Waltham, Massachusetts, U.S.) and Qubit (Thermo Fisher, Waltham, Massachusetts, U.S.). For sequencing, samples were organised in 3 batches of 80 samples, including 2 to 5 technical replicates for 8 samples. Library preparation and sequencing was performed at the GeT-PlaGe core facility located in Toulouse (https://get.genotoul.fr). Libraries were constructed using the TruSeq DNA Nano kit from Illumina® (San Diego, California, U.S.) according to the manufacturer's instructions. Sequencing was performed on the Illumina® NovaSeq system, producing 150-base pairs (bp) paired-end reads. Each of the three batches of 80 libraries was sequenced on one lane. By sequencing 80 individual libraries per lane, we aimed to obtain an average sequencing depth of ~10X, considering a reference genome of 800 Mb (Plomion *et al.*, 2018).

### S3. Methanolic extraction and liquid-chromatography

**Methanolic extraction:** We performed a methanolic extraction and weighed 20 mg (± 2 mg) of lyophilized leaf powder samples in 1.2 mL Micronic® tubes in 96 tube-plates.

For each plate, we added 500 µl of extraction solvent composed of 70%v acidified methanol with 0.1%v acid formic, 30%v $H_20$ and 0.1 g/L quercetin internal standard in each tube. Note that quercetin was chosen as an internal standard after checking it did not obscure the signal from other molecules. While quercetin is abundant in oaks, all abundant forms are linked to sugars. The unglycosylated form we used as an internal standard had a longer retention time than other glycosylated quercetin-based flavonoids. Plates were vortexed for 5 min at medium speed and then sonicated in a bath of iced water for 15 min and vortexed again for 5 min. For each sample, we filtered supernatant on 0.22 µm membranes using a Millipore vacuum manifold (Merck Millipore, Burlington, Massachusetts, United States) and a Fisher vacuum pump during two min. We prepared “Quality control” (QC) samples by mixing 2 µl of each sample, excluding blanks and technical replicates. We sealed plates with peaceable push-caps and stored them at -20°C until analysis.

**Liquid chromatography:** For all methanolic extracts, the separation of extracts was achieved using reverse phase liquid chromatography with a Gemini 3-µm C18 column (150 x 2 mm, Phenomenex, Torrance, CA, USA) on a Ultimate 3000 liquid chromatograph (ThermoFisher Scientific, Sunnyvale, CA, USA) equipped for passing 96-well plates. The flow was set to 300 µL/min and the injection volume to 5 µL. Ultrapure water acidified with 0.1% formic acid and LC-MS grade acetonitrile were used as solvent A and B, respectively, with the following 10-min gradient: 0 min, 3%B; 1 min, 10%B; 6.5 min, 45%B; 7 min, 100%B; 7.5 min, 100%B; 8 min, 3%B; 10 min, 3%B. Each injection sequence was composed of an initialization phase with 10 injections of water and 10 injections of QC samples followed by injection of samples or blanks with injection of QC samples at regular intervals each 12 injections. The samples were maintained at a temperature of 6°C in the autosampler.

### S4. LTQ-ORBITRAP parameters

HESI electrospray ionisation probes operating in the positive and negative modes were respectively (50-1500 m/z range, 30k resolving power, 300°C capillary temperature, 300°C source heater temperature, 40 a.u. sheath gas flow, 20 a.u. auxiliary gas flow, 5 a.u. sweep gas flow, 3.2 (pos) or 2.5 (neg) kV source voltage, 55% S-lens RF level, HCD activation mode, 35% (pos) or 60% (neg) NCE).

### References

Larue, C. *et al.* (2021) ‘Development of highly validated SNP markers for genetic analyses of chestnut species’, *Conservation genetics resources*, 13(4), pp. 383–388.

Plomion, C. *et al.* (2018) ‘Oak genome reveals facets of long lifespan’, *Nature Plants*, 4(7), pp. 440–452.

**Table S1 Geographic origin of the populations and number of oak trees analysed per population.**

| Populations | Code | Latitude (°) | Longitude (°) | Phenotyped (n) | Sequenced (n) | Genotyped (n) |
|---|---|---|---|---|---|---|
| **Saint-Sauvant** | 9 | 46.38 | 0.12 | 25 | 25 | 25 |
| **Grésigne** | 97 | 44.04 | 1.75 | 25 | 25 | 25 |
| **Bézanges** | 204 | 48.76 | 6.49 | 28 | 28 | 28 |
| **Bercé** | 217 | 47.81 | 0.39 | 25 | 25 | 24 |
| **Longchamp** | 218 | 47.26 | 5.31 | 22 | 21 | 20 |
| **Tronçais** | 219 | 46.68 | 2.83 | 25 | 25 | 25 |
| **Vachères** | 233 | 43.98 | 5.63 | 25 | 25 | 25 |
| **Göhrde** | 253 | 53.10 | 10.86 | 25 | 25 | 23 |
| **Lappwald** | 256 | 52.26 | 10.99 | 25 | 25 | 23 |

Each of the nine populations included in the study is identified by a name (column "Populations") and a code (given by column "Code"). Latitude and longitude of the population sampled are given in degree decimal (datum WGS84). The columns "Phenotyped", "Sequenced", and "Genotyped" give the numbers of trees that were phenotyped with untargeted LC-MS analyses, sequenced at a depth of 10X, and successfully genotyped, respectively.

5

**Table S2 Tables of oak trees and sample counts for each analysis in this study.**

**a.** SNPs and genotyping dataset

| | **Oak trees** | **Commentary** |
|---|---|---|
| Sampling (2016) | 225 | |
| DNA extraction and sequencing | 224 | After extraction, DNA for one individual didn’t pass quality check before sequencing |
| SNPs & genotypes matrix | 218 | Six individuals removed (high sequence similarity) |

**b.** Annotation dataset

| | | **Oak trees** | **Leaf samples high branches** | **Leaf samples low branches** | **Technical replicate samples** | **Total number of leaf samples** | **Extraction blanks** | **Total counts** | **Commentary** |
|---|---|---|---|---|---|---|---|---|---|
| Sampling | 2016 | 54 | 54 | 53 | 0 | 107 | 0 | 215 | Samples from 2021 not used in the present study |
| | 2021 | 54 | 54 | 54 | 0 | 108 | | | |
| Methanolic extractions | 2016 | 54 | 54 | 53 | 25 | 132 | 29 | 285 | Remove one sample to keep pairs of samples between 2016 and 2021 samples |
| | 2021 | 54 | 54 | 53 | 20 | 127 | | | |

6

**Table S2 continued**

**c.** GWA dataset

| | Oak trees | Leaf samples high branches | Leaf samples low branches | Biological replicates | | | Time serie samples | Technical replicate samples | Total number of leaf samples | Extraction blanks | Total counts | Commentary |
|---|---|---|---|---|---|---|---|---|---|---|---|---|
| | | | | **H** | **L** | **U** | | | | | | |
| Sampling | 225 | 225 | 225 | 42 | 46 | 20 | 24 | 0 | 582 | 0 | 582 | |
| Methanolic extractions | 225 | 225 | 223 | 41 | 48 | 0 | 24 | 99 | 661 | 107 | 768 | eight 96-well plates |
| Dataset Figure S1 | 225 | 215 | 209 | 42 | 40 | 0 | 0 | 93 | 599 | 91 | 712 | outlier filtering |
| Dataset for PCA, PLS-DA and GWAs | 225 | 211 | 206 | 0 | 0 | 0 | 0 | 0 | 417 | 0 | 417 | 192 trees are in common between high and low branches |

In the column "biological replicates", the sub-columns H and L refer to samples harvested from high and low branches, respectively. The sub-column U refers to samples for which branch height was undetermined.

7

**Table S2 continued**

**d.** GWAs analysis

| | Oak trees | Leaf samples high branches | Leaf samples low branches | Commentary |
|---|---|---|---|---|
| SNPs & genotyping | 218 | 218 | 218 | |
| GWA dataset | 225 | 211 | 206 | 192 individuals with data for both branch heights |
| GWA analysis | 217 | 203 | 200 | Removed individuals due to missing values in metabolomics and/or genotypes. Overall, 187 genotyped individuals had phenotypes for both high and low branches. |

8

**Table S3** Optimised parameters used for peak picking and peak alignment in the processing of the data obtained from the LC-QTOF-MS (GWA dataset)

| IPO parameters | Value |
|---|---|
| **min_peakwidth** | 6.75 |
| **max_peakwidth** | 10.6 |
| **ppm** | 75 |
| **bw** | 0.25 |
| **snthres** | 5 |
| **mzdiff** | -0.0065 |
| **prefilter** | 3 |
| **value_of_prefilter** | 100 |
| **gapInit** | 0.352 |
| **gapExtend** | 2.208 |
| **minsamp** | 1 |
| **max** | 50 |

These values were chosen based on the result of the IPO R package v1.18, designed to optimize the parameters of the functions “PeakDensityParam”, “adjustRtime” and “CentWaveParam” from the R package XCMS.

**Figure S1** Relationships between the datasets produced

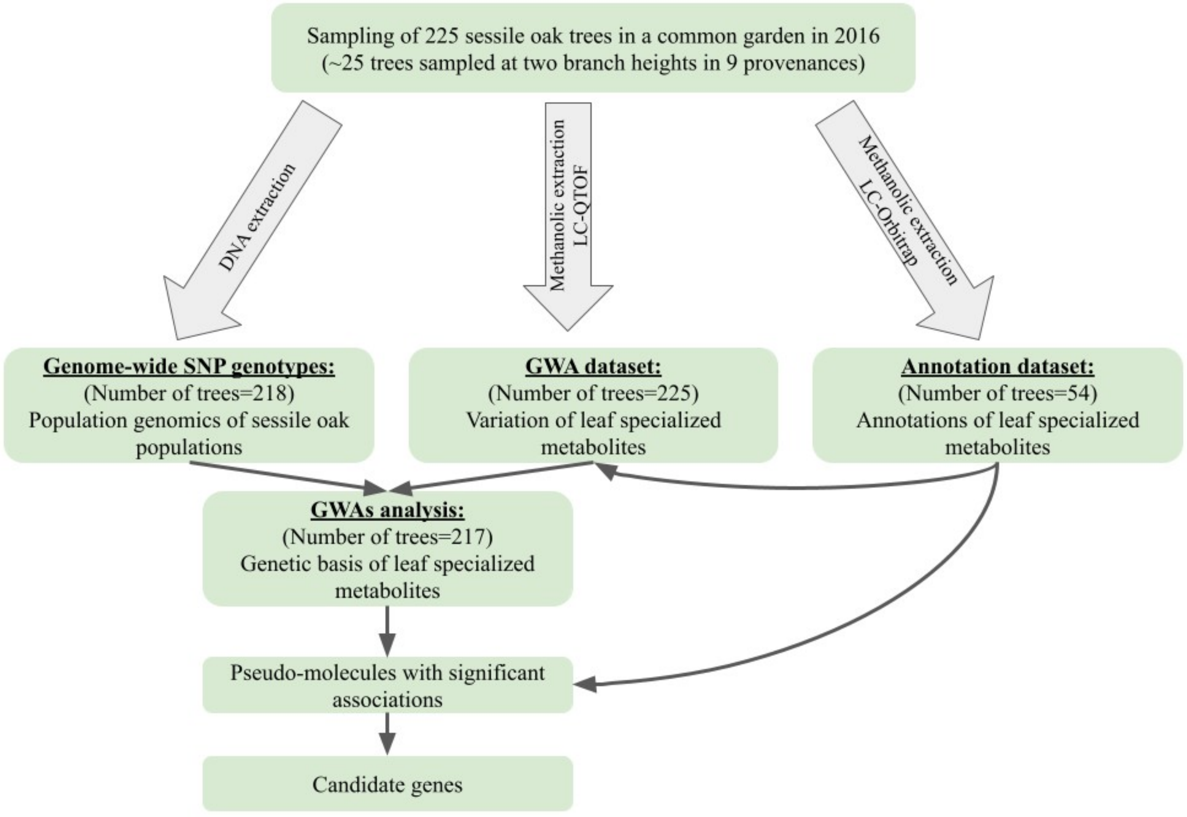

**Figure S2** Potential confounding effects on specialized metabolites variation in our study.

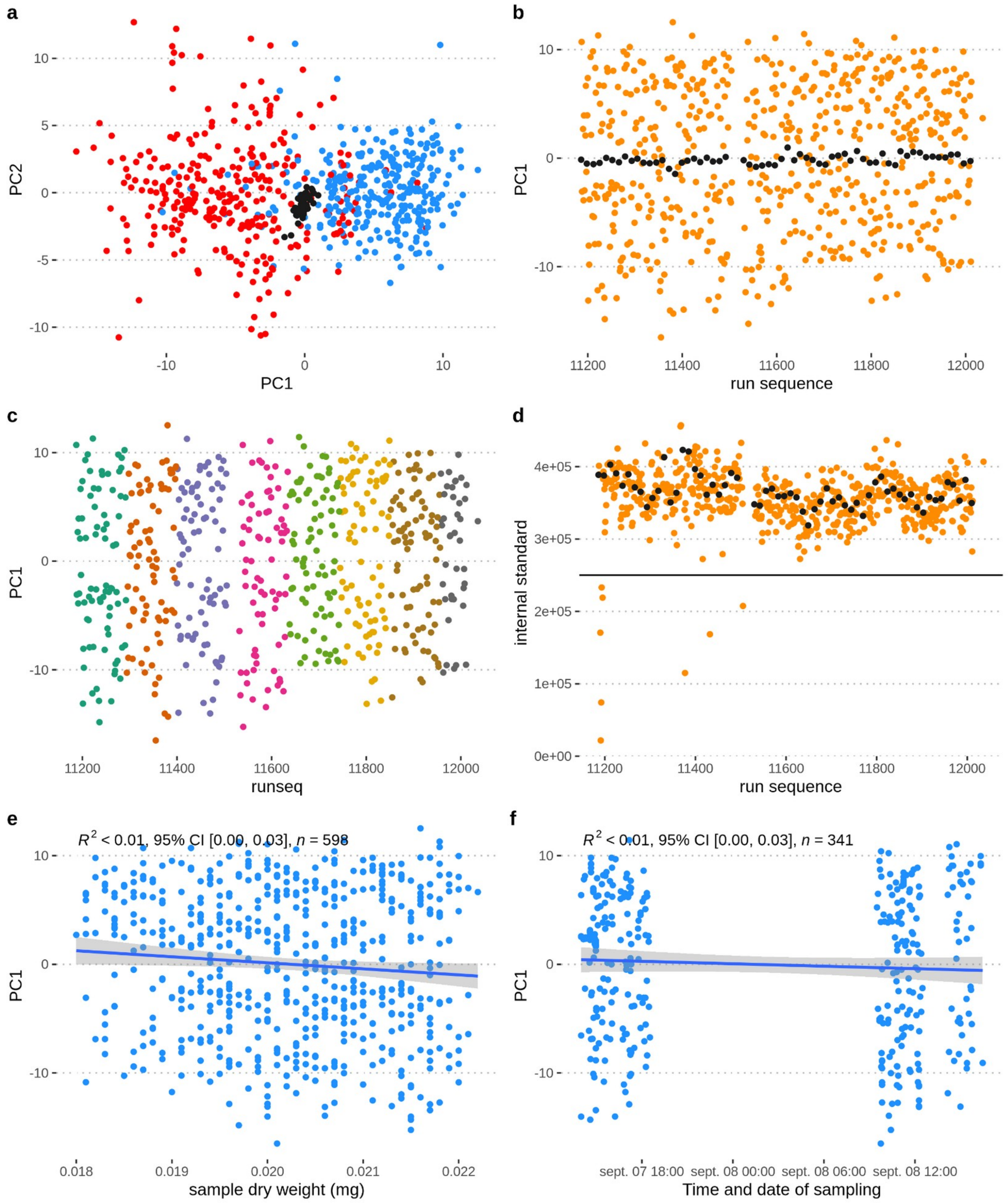

**a.** Principal component analysis of samples variance (N=599 samples + 61 QCs) including leaf samples, technical replicates and QCs samples. The points are coloured according to height (high and low) at which leaf samples were collected in the trees. Black points correspond to QCs samples (N=61). **b.** Effect of sample run sequence on specialized metabolite variation. Each point corresponds to an injection, black dots correspond to QCs and orange dots to all other sample types, including replicates. The x-axis is the run sequence and the y-axis (PC1) reports the coordinates of each sample along the first component of the PCA in **a**. **c.** Same as **b**, except QCs are not represented and points are coloured according to the 96-well plate used for the methanolic extraction. **d.** Intensity of the quercetin internal standard in samples is represented on the y-axis along the injection sequence on the x-axis. The horizontal black line represents a threshold below which samples were removed **e.** Effect of samples dry weight on specialized metabolite variation. Each point corresponds to a sample, the x-axis is the sample dry weight and the y-axis the coordinates of samples on the PC1 of the PCA plotted in **a. f.** Effect of sampling time of leaf samples on specialized metabolites variation. Each point corresponds to a leaf sample, the x-axis represents the sampling date and time of the sample and the y-axis the coordinates of the sample along the PC1 of the PCA plotted in **a.**

10

**Figure S3** Decay of linkage disequilibrium in the sample of 225 *Quercus petraea* genotypes from 9 European populations.

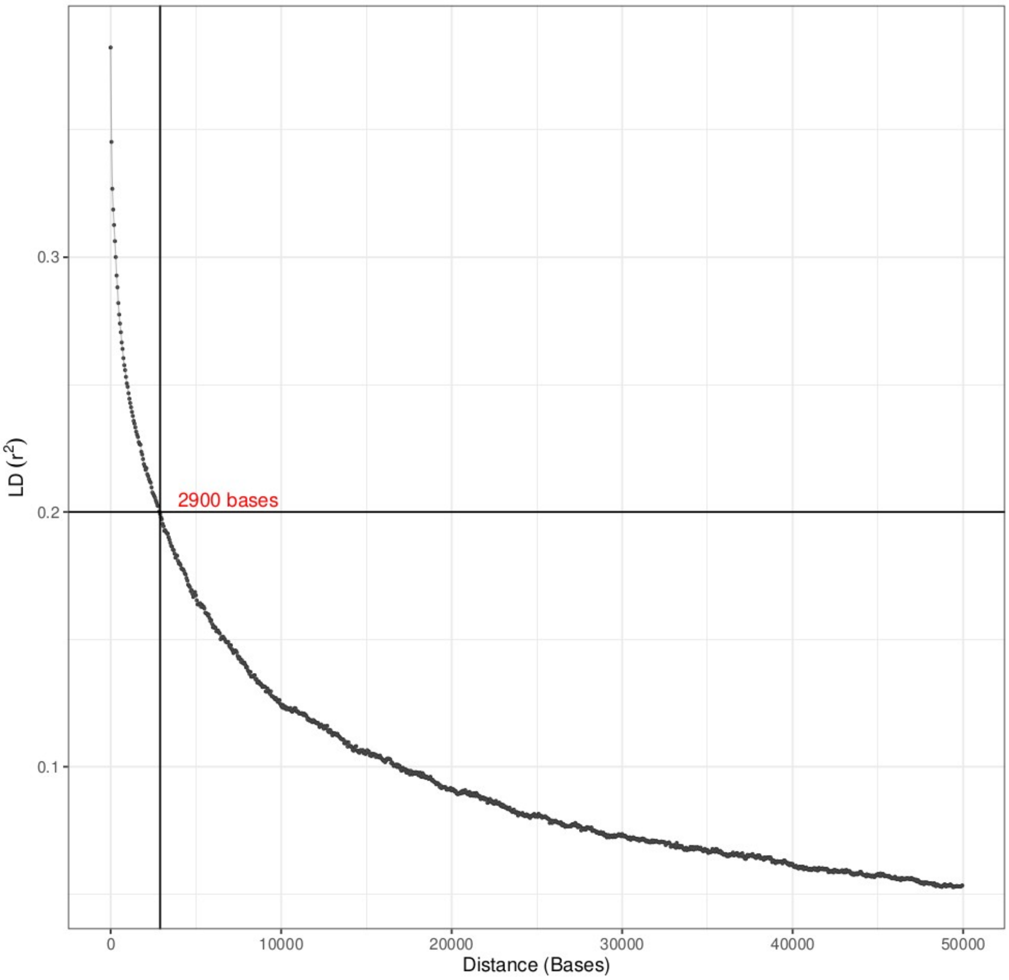

The x-axis represents the distance between pairs of markers (in bases). The y-axis represents linkage disequilibrium ($r^2$) between markers. Each point represents the mean $r^2$ of all markers with pairwise physical distances within windows of 500 bp. Mean r2 values were computed for all 500bp windows, from 0 to 50kb, taking steps of 50 bp. The vertical and horizontal black line represent $r^2 < 0.2$ at 2,900 base pairs.

11

**Figure S4** The variation of leaf specialized metabolite m181 among populations is largely explained by genetic variation at a single genetic marker.

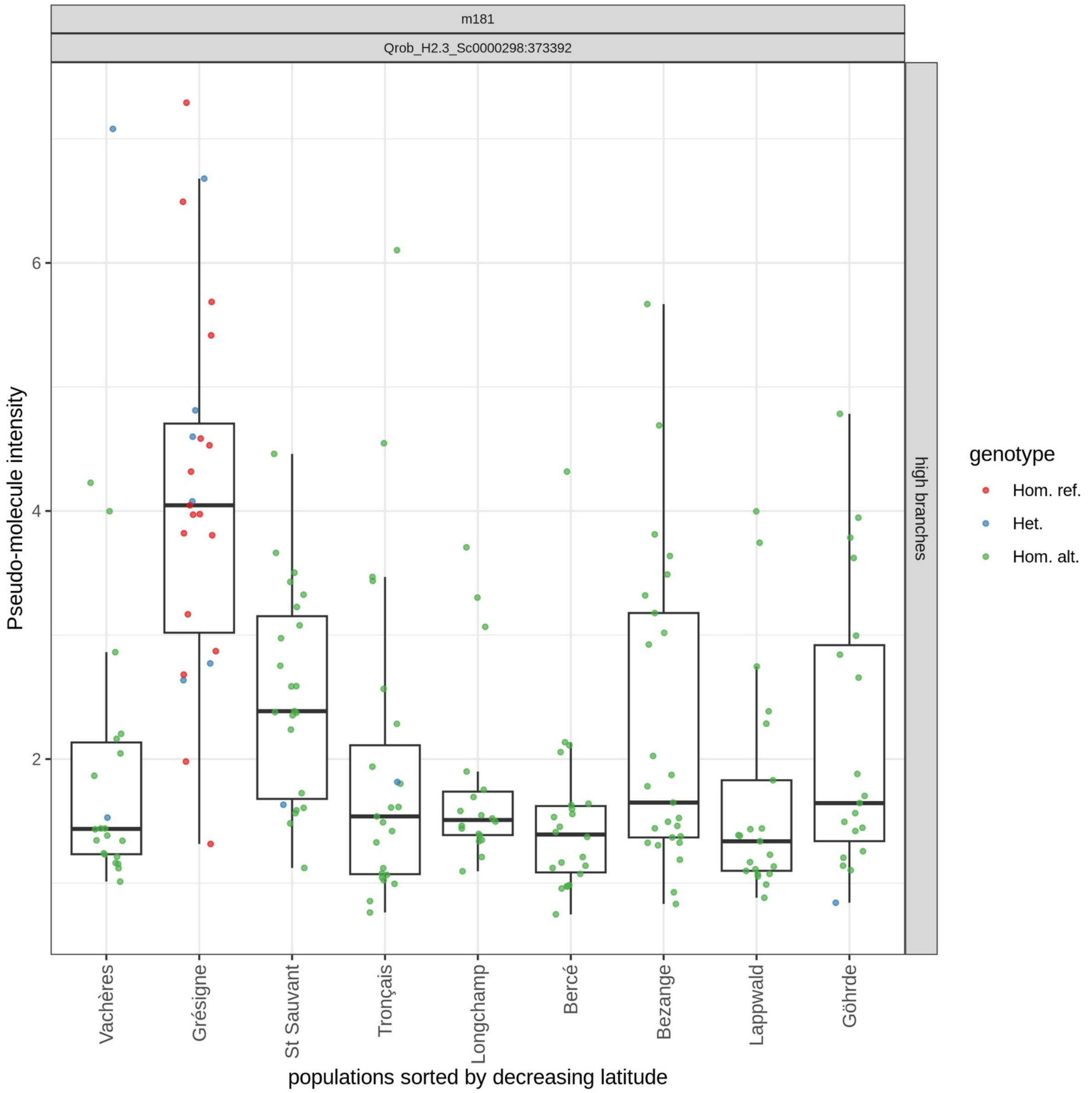

The plot represents the variation of the peak intensity (y-axis, cube-root transformation) explained by the most associated marker with the m181 pseudo-molecule. The title of the panel includes the identifier of the pseudo-molecule and the position of the most associated SNP to its variation (Scaffold: position in bp). Each point corresponds to a leaf sample collected on high branches. Point colours correspond to the genotype of the tree at the associated SNP (see legend on the right-hand side). “Hom. ref.” (red points) stands for “homozygous for the reference allele”, “Het.” (blue points) stands for “heterozygous” and “Hom. alt.” (green points) stands for “homozygous for the alternative allele”. Note that a small jitter was added to positions along the x-axis for clarity. Points are grouped by population of origin along the x-axis. The horizontal bars in the boxes represent the median for each population. The boxes include 50% of the samples between the third and third quartile of the phenotypic distribution and the whiskers extend to the largest or smallest value observed, but no further than 1.5 times the distance between the first and third quartiles.

12

**Figure S5** Phenotypic variation explained by genetic variation at the most associated marker for each pseudo-molecule.

In all panels, the plots represent the variation of the peak intensity (y-axis, cube root transformation and Pareto scaling) for 93 pseudo-molecules for leaves collected on high and low branches. Points correspond to leaves of individual samples in high branches and low branches, as indicated in the title of each panel. Points are grouped by population along the x-axis and populations are ordered by increasing latitude of origin. Note that a jitter was applied to the location of points along the x-axis for clarity. Points are coloured according to the genotype at the SNP most strongly associated with each pseudo-molecule as described in the legend on the left. "Hom. ref." (red points) stands for "homozygous for the reference allele", "Het." (blue points) stands for "heterozygous" and "Hom. alt." (green points) stands for "homozygous for the alternative allele". The position of the SNP associated with each pseudo-molecule is given in the panel title.

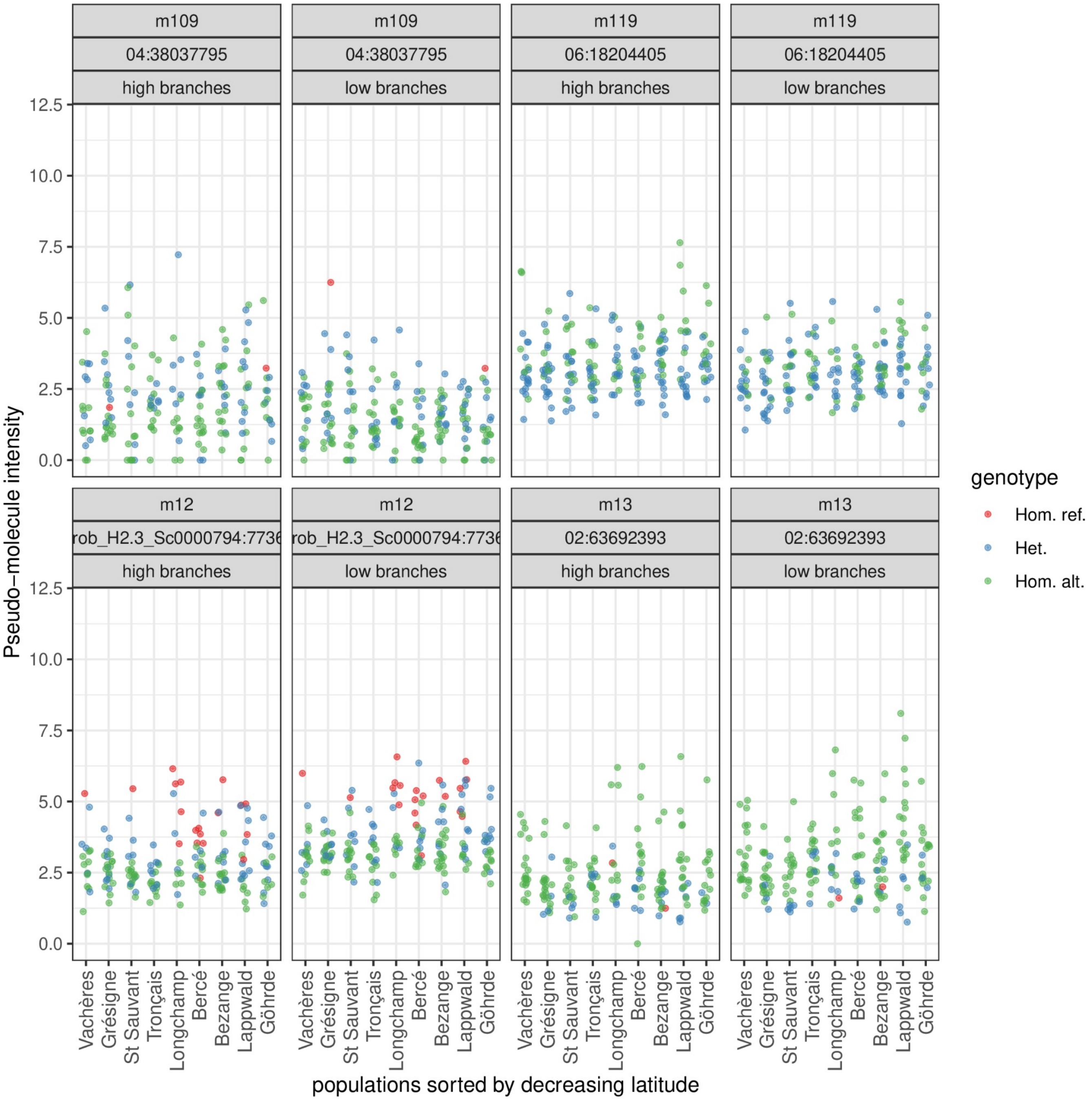

 13

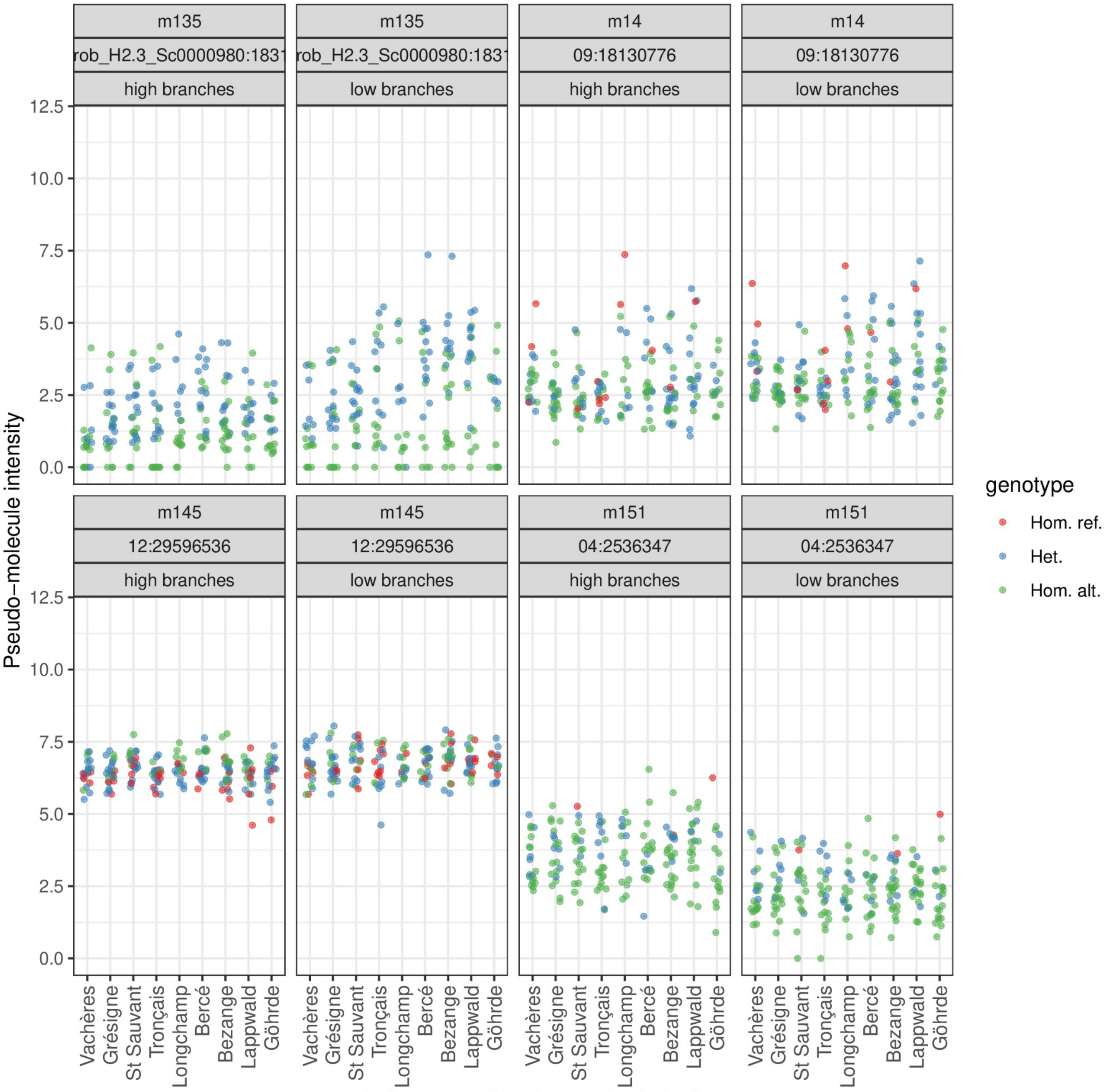
m135
rob_H2.3_Sc0000980:1831
high branches
low branches
m14
09:18130776
m145
12:29596536
m151
04:2536347
Pseudo−molecule intensity
12.5
10.0
7.5
5.0
2.5
0.0
genotype
Hom. ref.
Het.
Hom. alt.
Vachères
Grésigne
St Sauvant
Tronçais
Longchamp
Bercé
Bezange
Lappwald
Göhrde
populations sorted by decreasing latitude

14

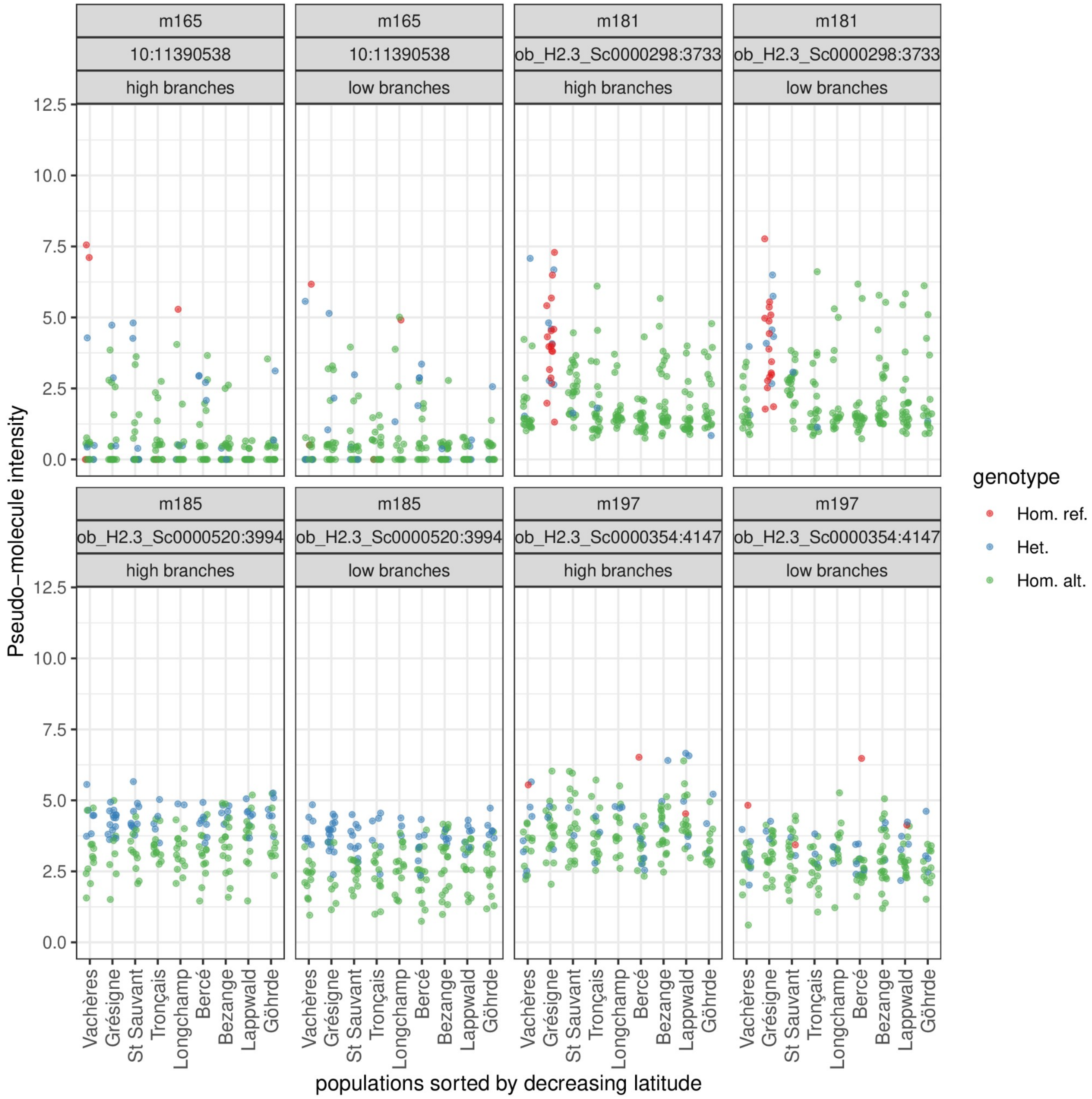
m165
10:11390538
high branches
low branches
m181
ob_H2.3_Sc0000298:3733
m185
ob_H2.3_Sc0000520:3994
m197
ob_H2.3_Sc0000354:4147
Pseudo−molecule intensity
0.0
2.5
5.0
7.5
10.0
12.5
genotype
Hom. ref.
Het.
Hom. alt.
Vachères
Grésigne
St Sauvant
Tronçais
Longchamp
Bercé
Bezange
Lappwald
Göhrde
populations sorted by decreasing latitude

 15

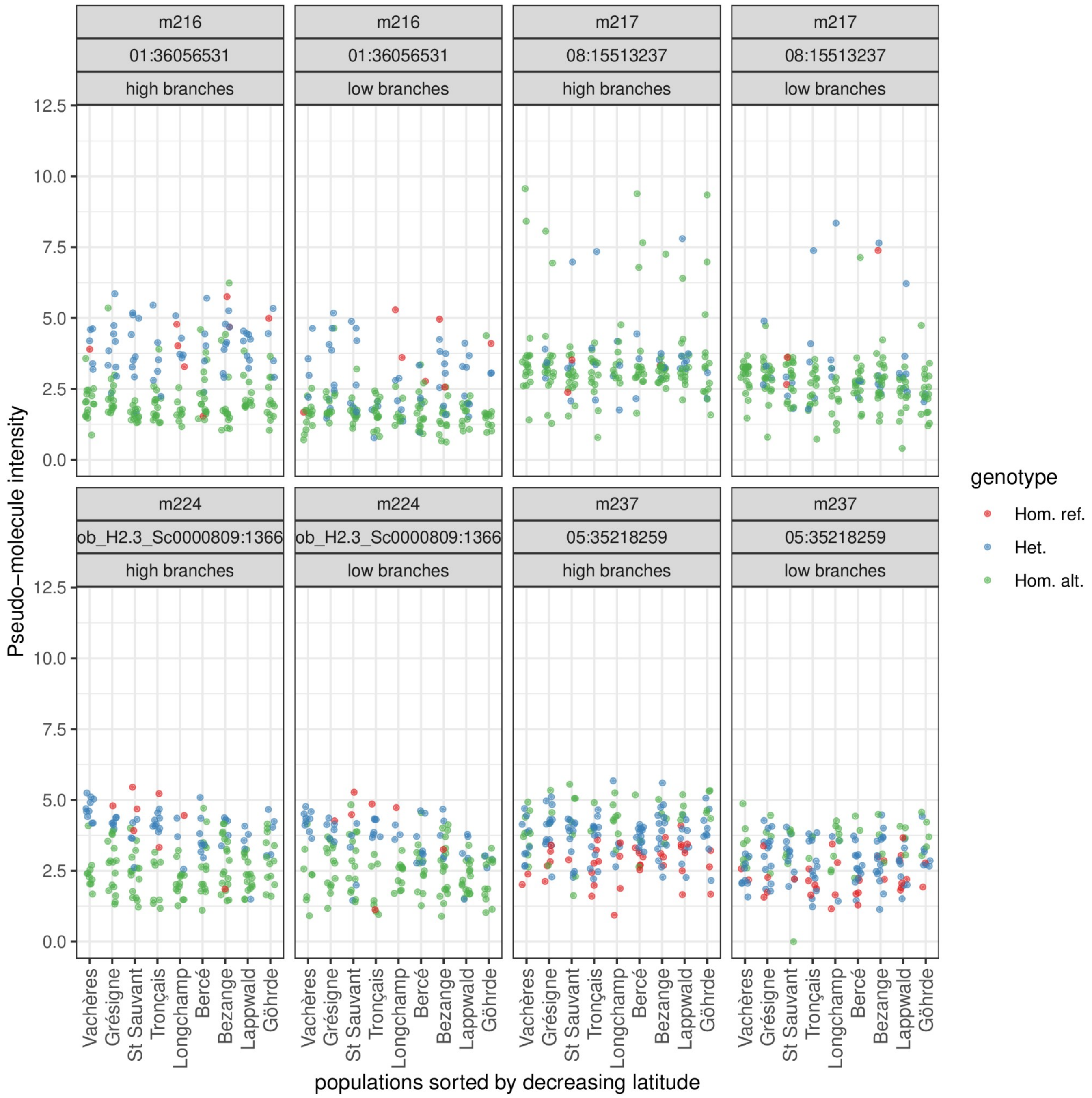

m216
01:36056531
high branches
low branches
m217
08:15513237
m224
ob_H2.3_Sc0000809:1366
m237
05:35218259
Pseudo−molecule intensity
0.0
2.5
5.0
7.5
10.0
12.5
Vachères
Grésigne
St Sauvant
Tronçais
Longchamp
Bercé
Bezange
Lappwald
Göhrde
populations sorted by decreasing latitude
genotype
Hom. ref.
Het.
Hom. alt.

16

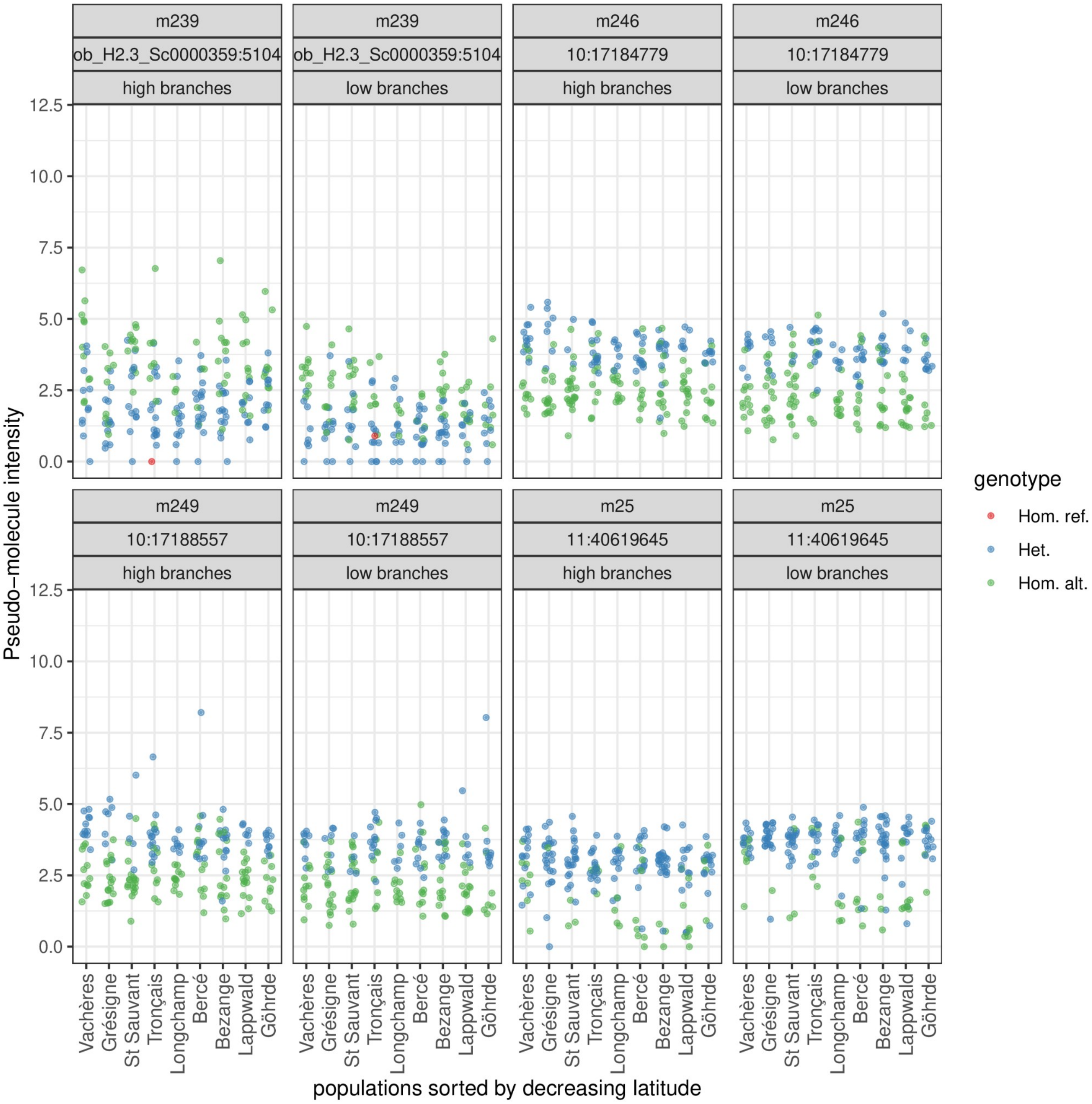
m239
ob_H2.3_Sc0000359:5104
high branches
low branches
m246
10:17184779
m249
10:17188557
m25
11:40619645
Pseudo−molecule intensity
0.0
2.5
5.0
7.5
10.0
12.5
Vachères
Grésigne
St Sauvant
Tronçais
Longchamp
Bercé
Bezange
Lappwald
Göhrde
populations sorted by decreasing latitude
genotype
Hom. ref.
Het.
Hom. alt.

17

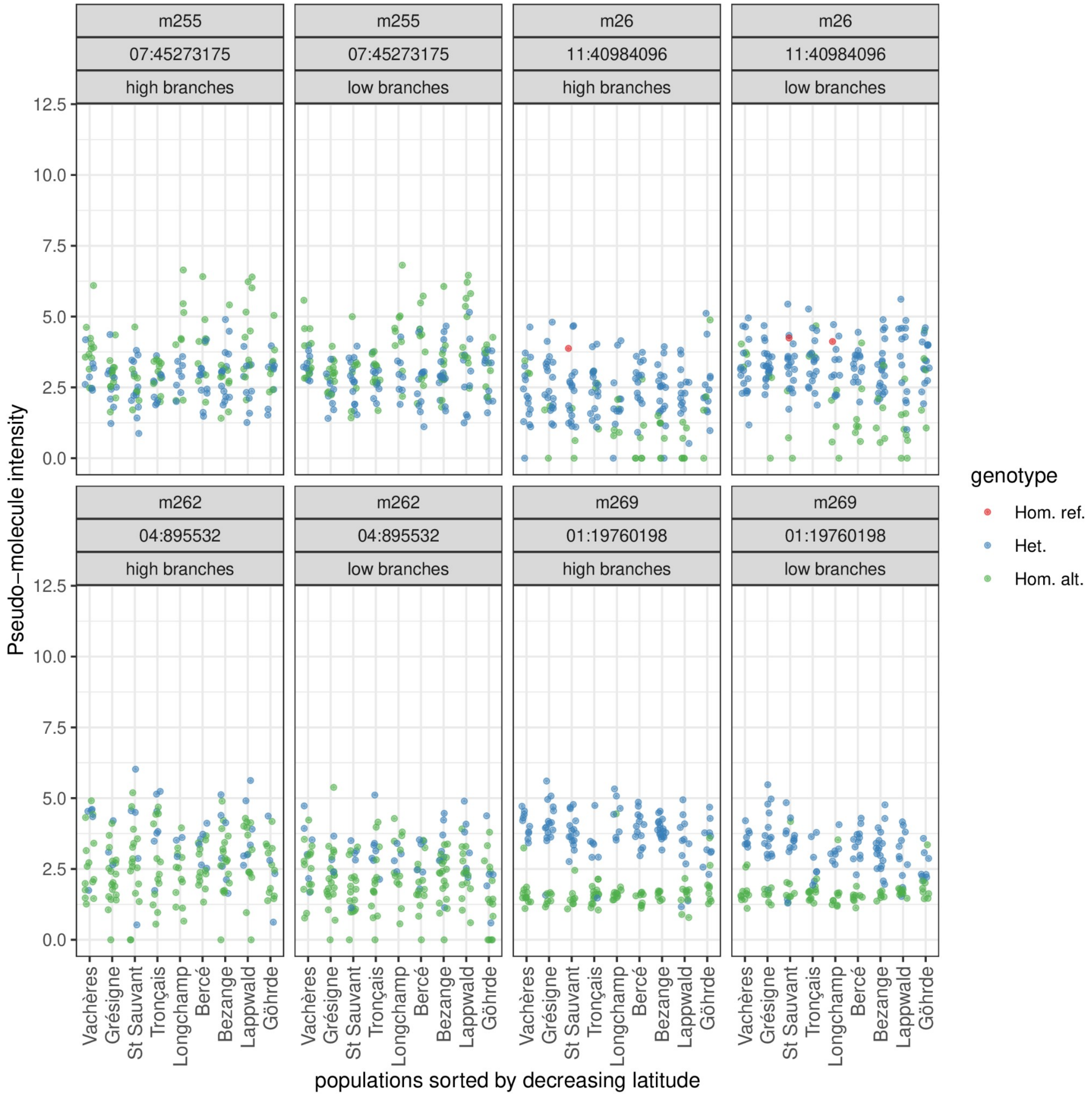
m255
07:45273175
high branches
low branches
m26
11:40984096
m262
04:895532
m269
01:19760198
Pseudo−molecule intensity
0.0
2.5
5.0
7.5
10.0
12.5
Vachères
Grésigne
St Sauvant
Tronçais
Longchamp
Bercé
Bezange
Lappwald
Göhrde
populations sorted by decreasing latitude
genotype
Hom. ref.
Het.
Hom. alt.

 18

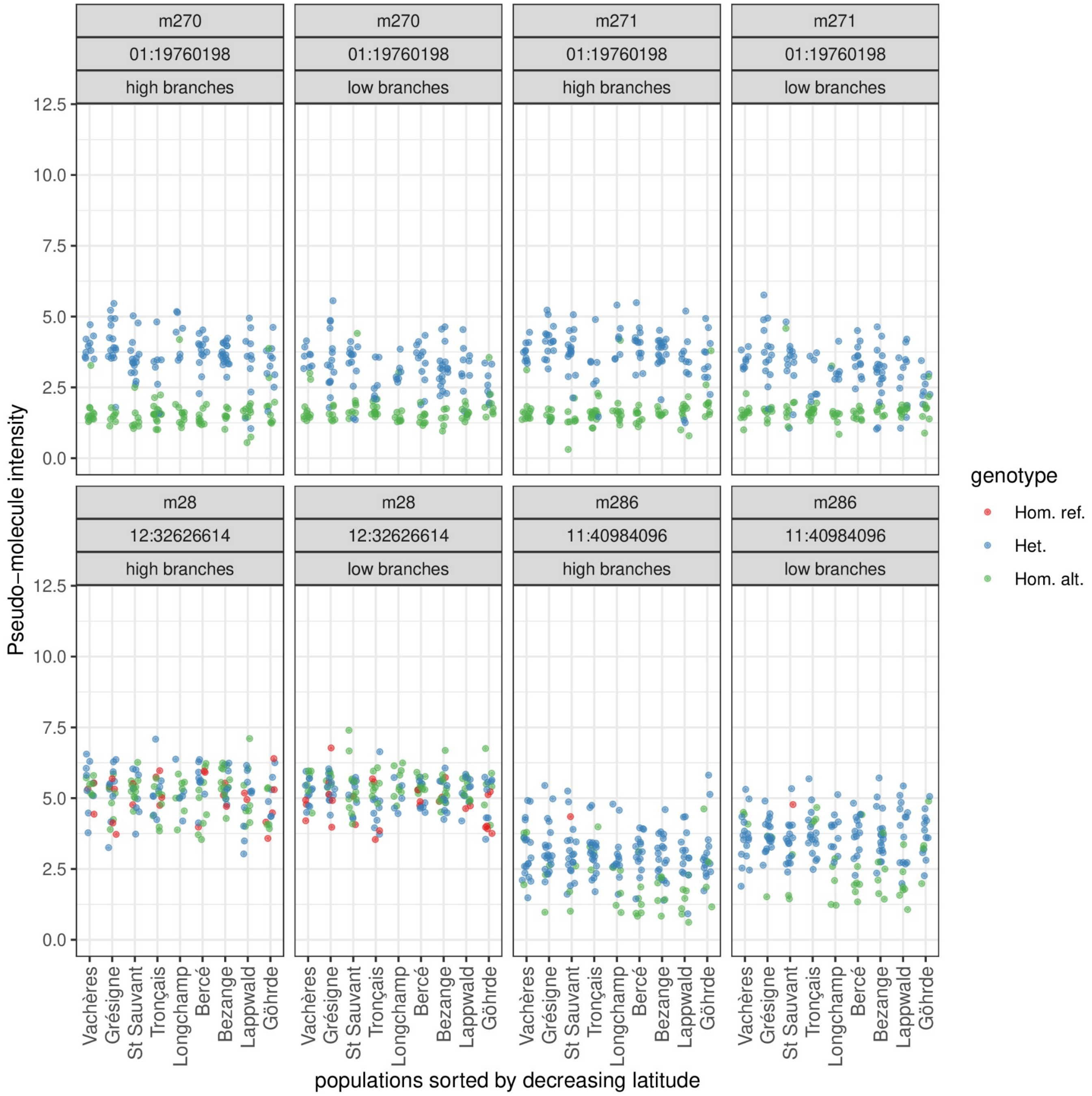
m270
01:19760198
high branches
m270
01:19760198
low branches
m271
01:19760198
high branches
m271
01:19760198
low branches
m28
12:32626614
high branches
m28
12:32626614
low branches
m286
11:40984096
high branches
m286
11:40984096
low branches
Pseudo−molecule intensity
0.0
2.5
5.0
7.5
10.0
12.5
Vachères
Grésigne
St Sauvant
Tronçais
Longchamp
Bercé
Bezange
Lappwald
Göhrde
populations sorted by decreasing latitude
genotype
Hom. ref.
Het.
Hom. alt.

 19

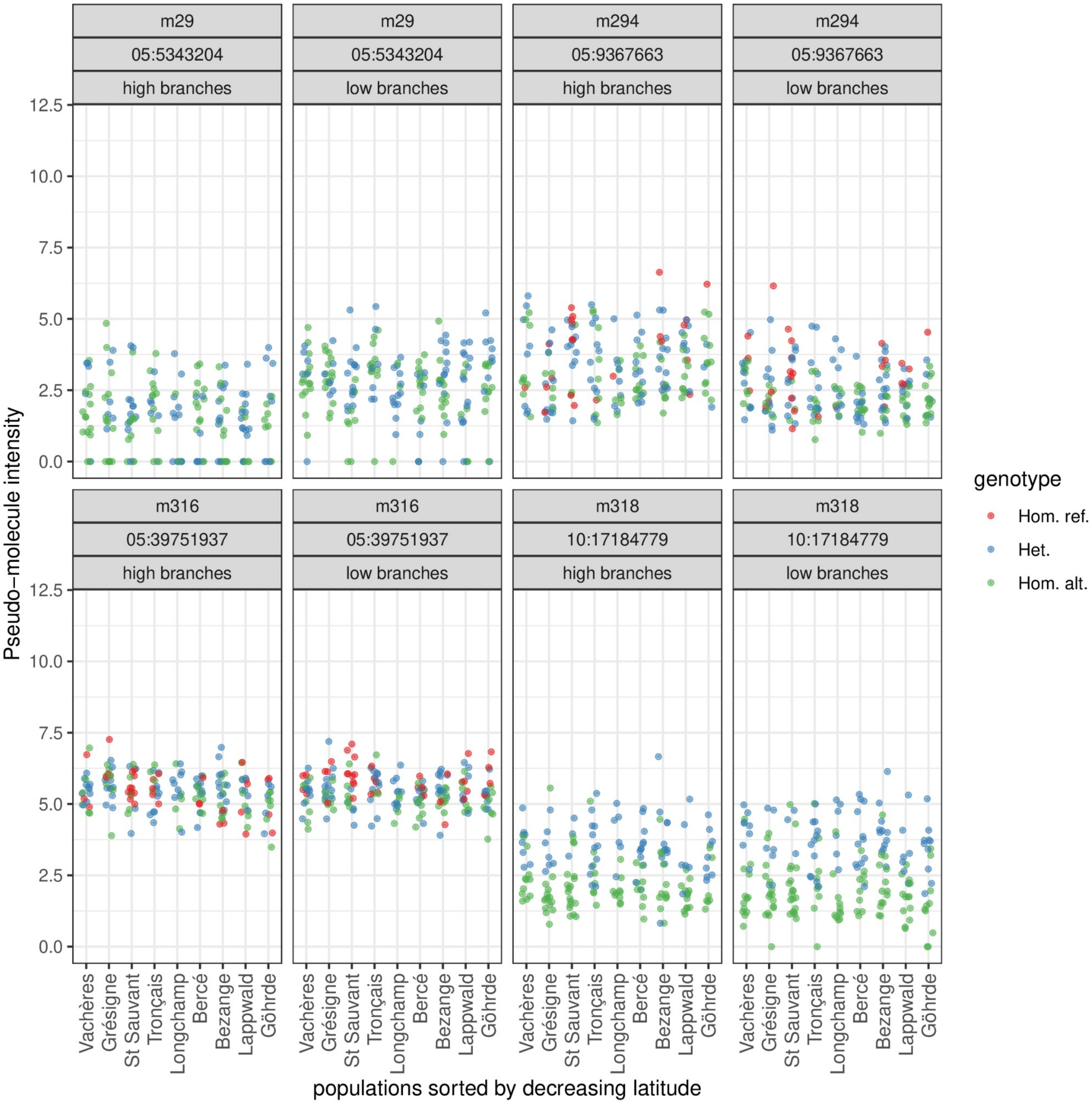
m29
05:5343204
high branches
low branches
m294
05:9367663
m316
05:39751937
m318
10:17184779
Pseudo−molecule intensity
0.0
2.5
5.0
7.5
10.0
12.5
Vachères
Grésigne
St Sauvant
Tronçais
Longchamp
Bercé
Bezange
Lappwald
Göhrde
populations sorted by decreasing latitude
genotype
Hom. ref.
Het.
Hom. alt.

20

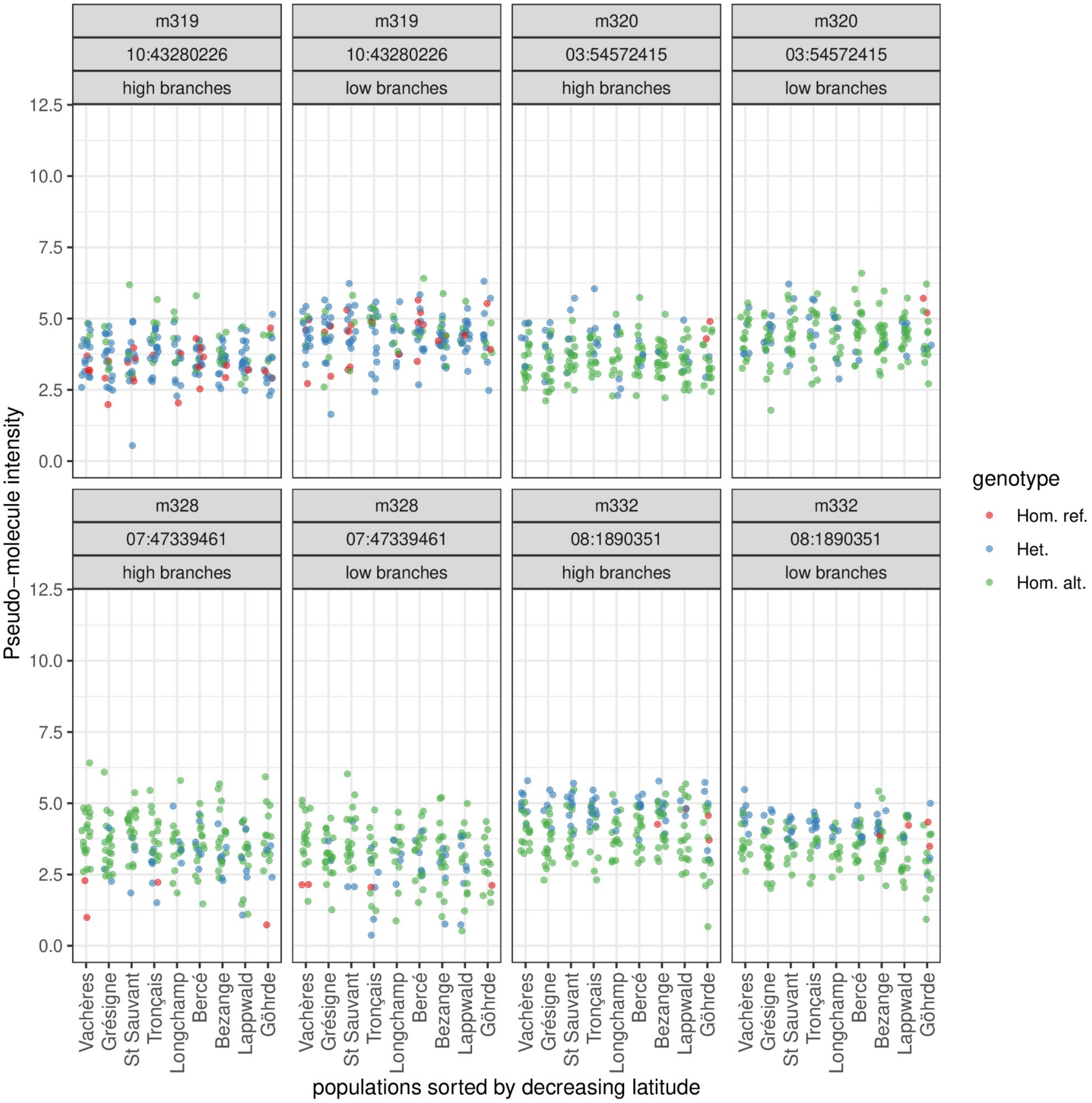
m319
10:43280226
high branches
m319
10:43280226
low branches
m320
03:54572415
high branches
m320
03:54572415
low branches
m328
07:47339461
high branches
m328
07:47339461
low branches
m332
08:1890351
high branches
m332
08:1890351
low branches
Pseudo−molecule intensity
0.0
2.5
5.0
7.5
10.0
12.5
Vachères
Grésigne
St Sauvant
Tronçais
Longchamp
Bercé
Bezange
Lappwald
Göhrde
populations sorted by decreasing latitude
genotype
Hom. ref.
Het.
Hom. alt.

21

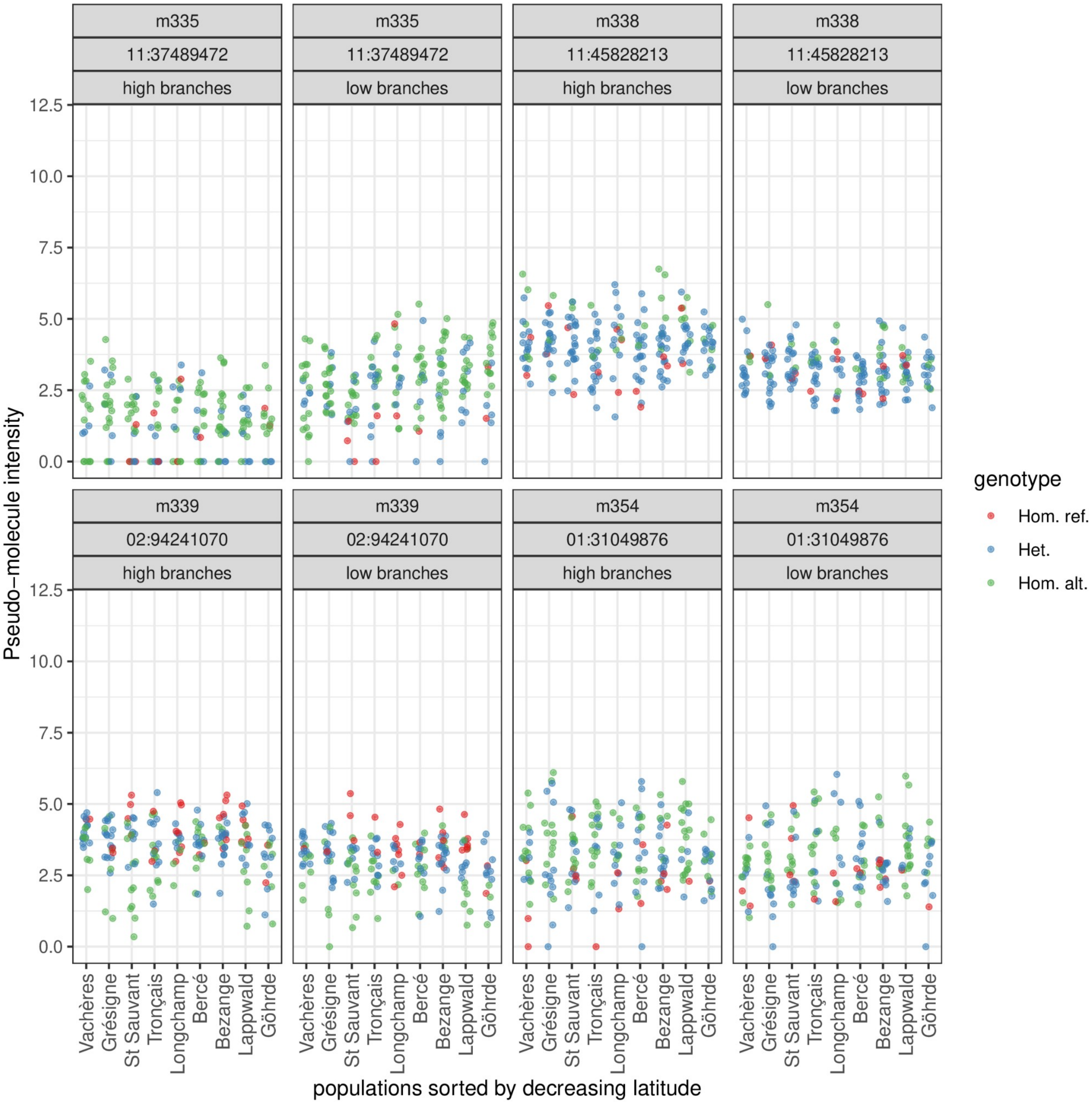
m335
11:37489472
high branches
low branches
m338
11:45828213
m339
02:94241070
m354
01:31049876
Pseudo−molecule intensity
0.0
2.5
5.0
7.5
10.0
12.5
genotype
Hom. ref.
Het.
Hom. alt.
Vachères
Grésigne
St Sauvant
Tronçais
Longchamp
Bercé
Bezange
Lappwald
Göhrde
populations sorted by decreasing latitude

 22

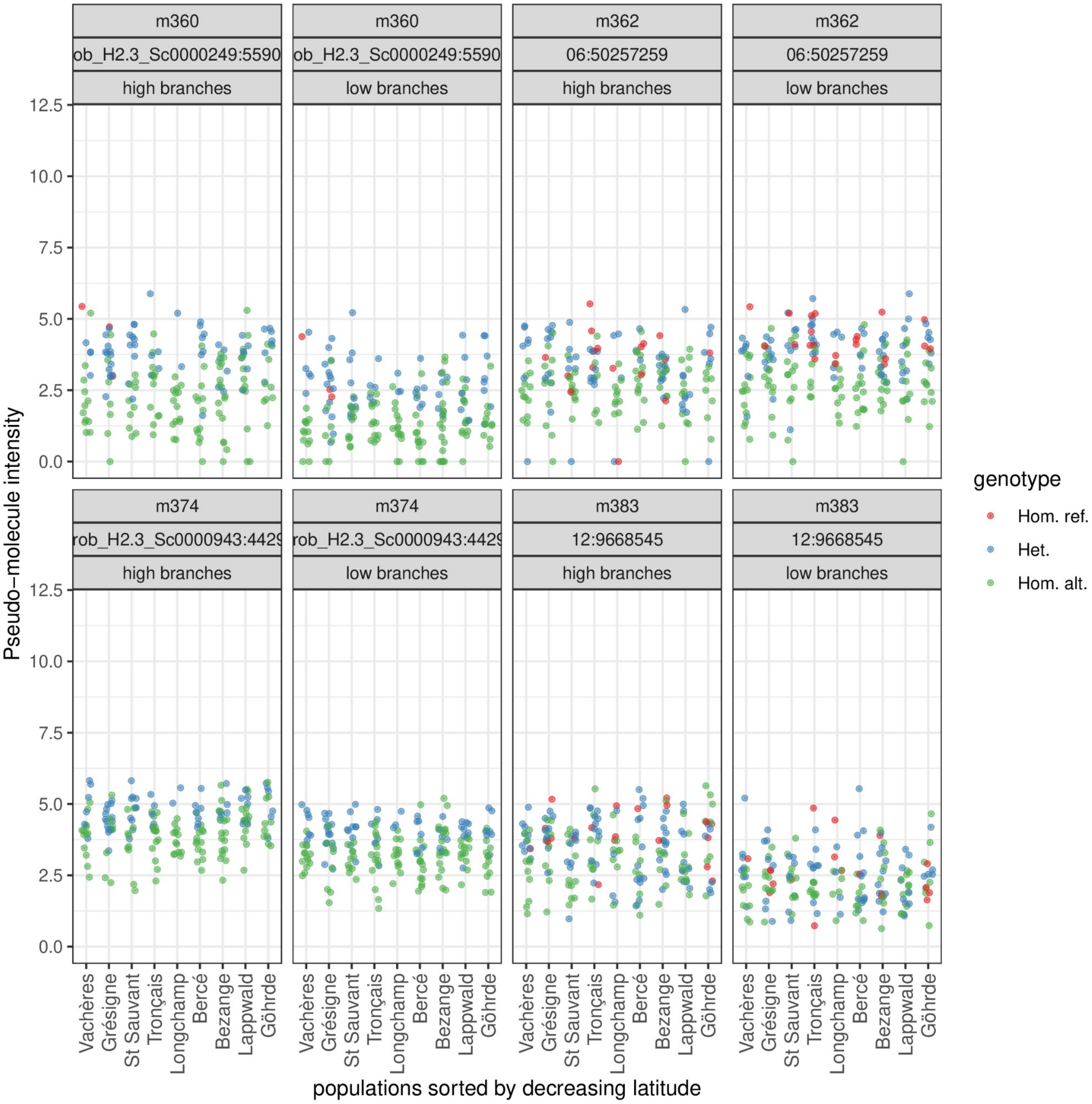
m360
ob_H2.3_Sc0000249:5590
high branches
low branches
m362
06:50257259
m374
rob_H2.3_Sc0000943:4429
m383
12:9668545
Pseudo−molecule intensity
0.0
2.5
5.0
7.5
10.0
12.5
Vachères
Grésigne
St Sauvant
Tronçais
Longchamp
Bercé
Bezange
Lappwald
Göhrde
populations sorted by decreasing latitude
genotype
Hom. ref.
Het.
Hom. alt.

23

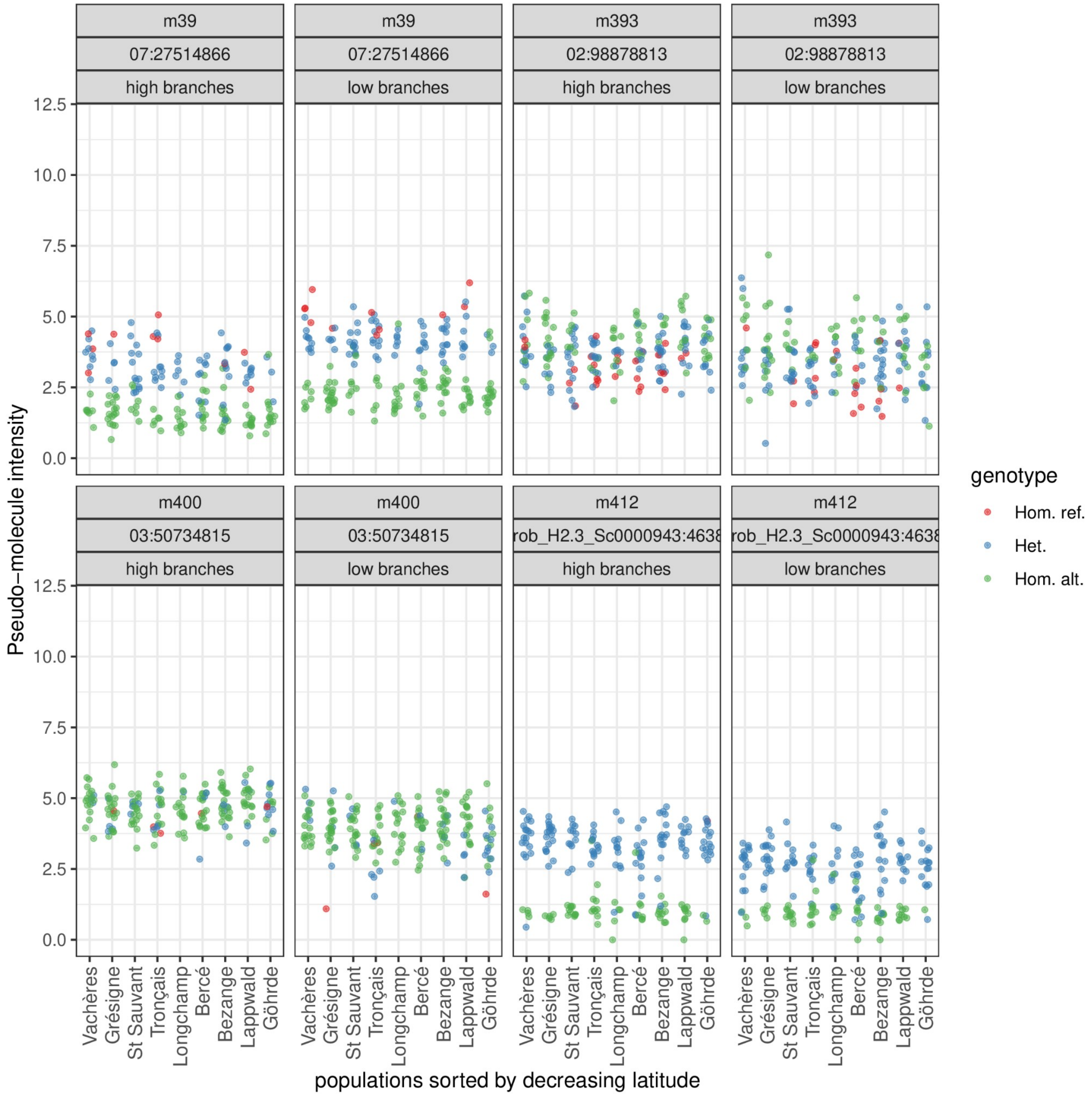
m39
07:27514866
high branches
low branches
m393
02:98878813
m400
03:50734815
m412
rob_H2.3_Sc0000943:4638
Pseudo−molecule intensity
0.0
2.5
5.0
7.5
10.0
12.5
Vachères
Grésigne
St Sauvant
Tronçais
Longchamp
Bercé
Bezange
Lappwald
Göhrde
populations sorted by decreasing latitude
genotype
Hom. ref.
Het.
Hom. alt.

198

199

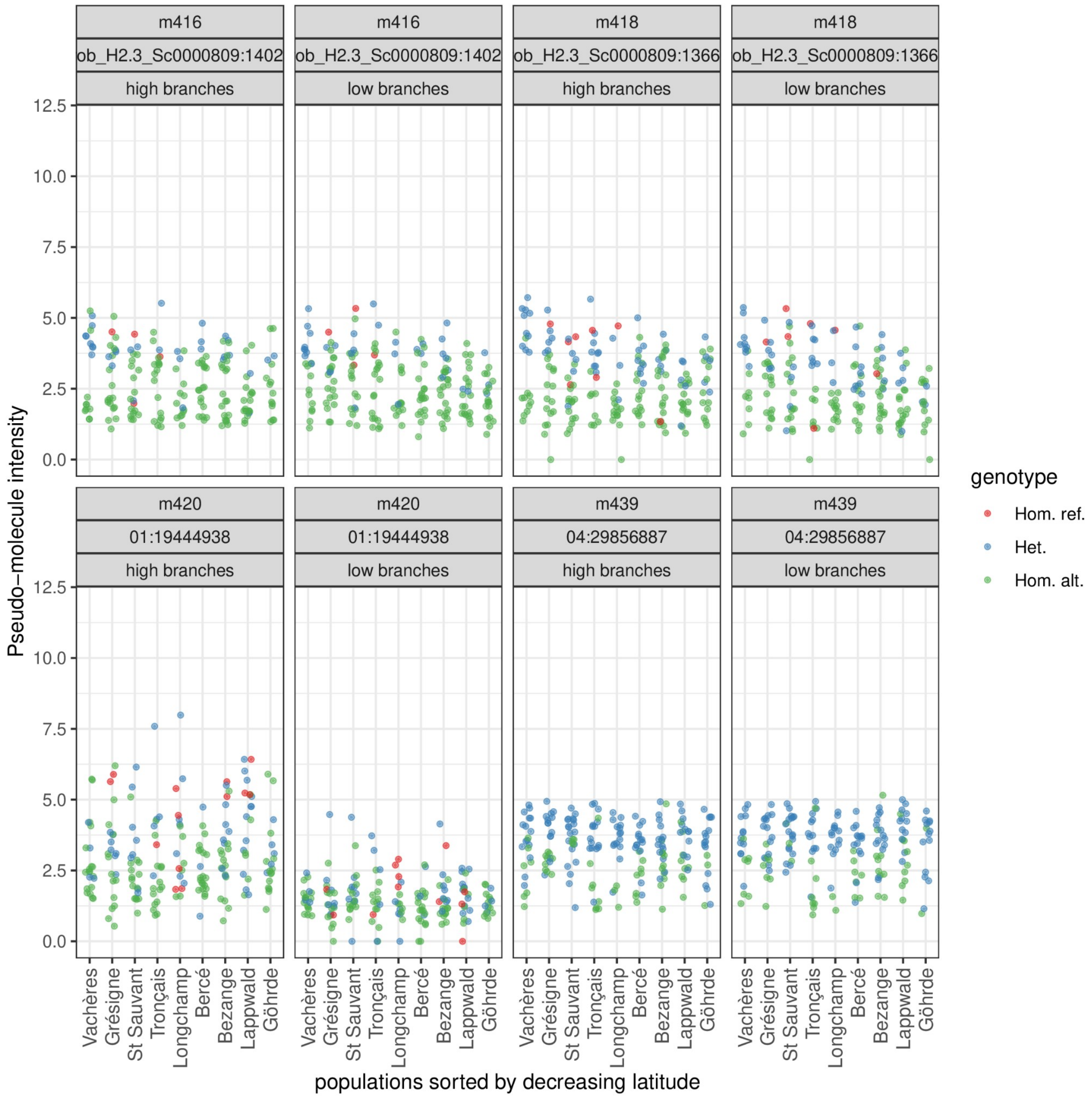

m416
ob_H2.3_Sc0000809:1402
high branches
low branches
m418
ob_H2.3_Sc0000809:1366
m420
01:19444938
m439
04:29856887
Pseudo−molecule intensity
0.0
2.5
5.0
7.5
10.0
12.5
Vachères
Grésigne
St Sauvant
Tronçais
Longchamp
Bercé
Bezange
Lappwald
Göhrde
populations sorted by decreasing latitude
genotype
Hom. ref.
Het.
Hom. alt.

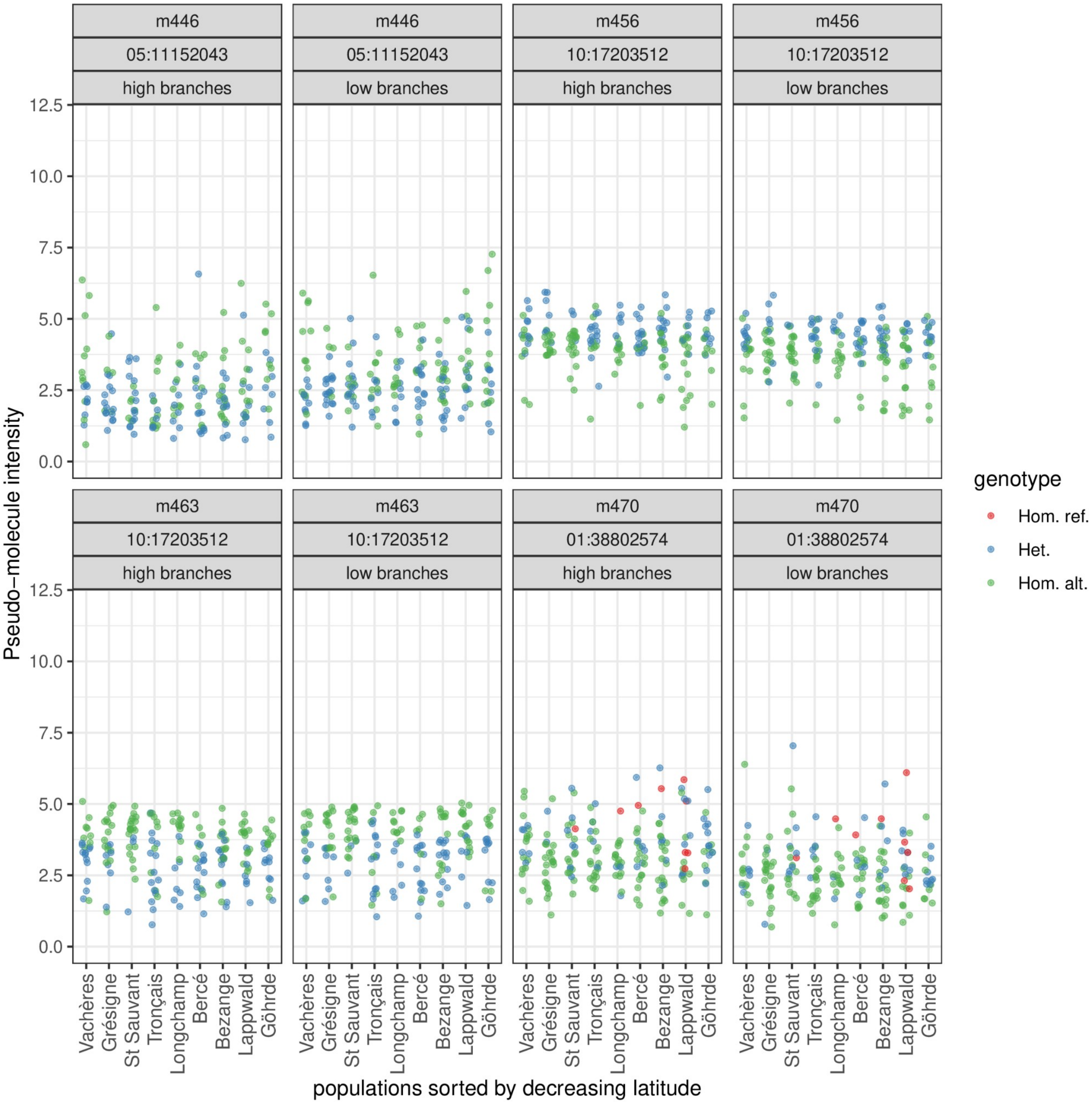
m446
05:11152043
high branches
m446
05:11152043
low branches
m456
10:17203512
high branches
m456
10:17203512
low branches
m463
10:17203512
high branches
m463
10:17203512
low branches
m470
01:38802574
high branches
m470
01:38802574
low branches
Pseudo−molecule intensity
0.0
2.5
5.0
7.5
10.0
12.5
Vachères
Grésigne
St Sauvant
Tronçais
Longchamp
Bercé
Bezange
Lappwald
Göhrde
populations sorted by decreasing latitude
genotype
Hom. ref.
Het.
Hom. alt.

26

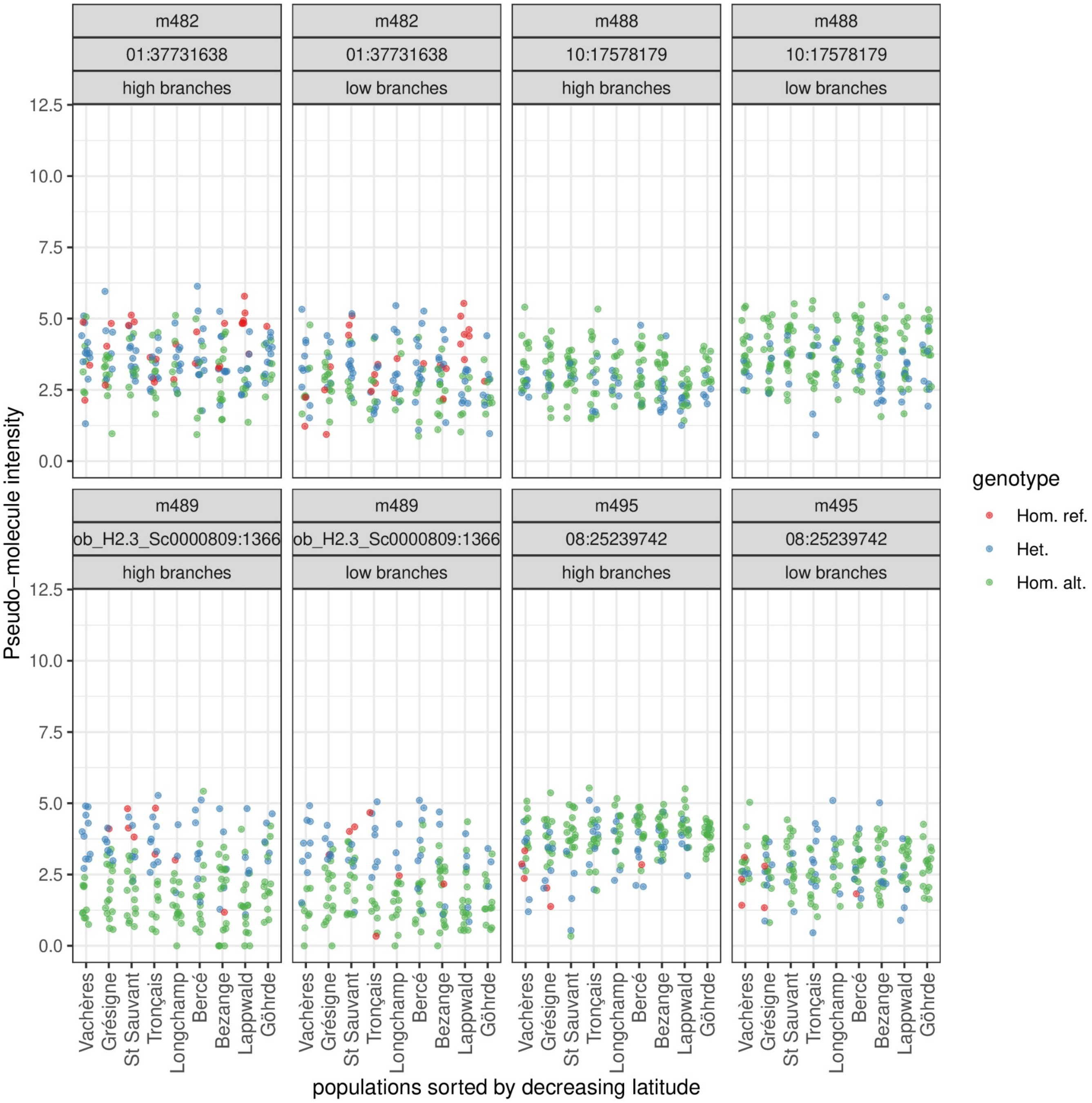
m482
01:37731638
high branches
low branches
m488
10:17578179
m489
ob_H2.3_Sc0000809:1366
m495
08:25239742
Pseudo−molecule intensity
12.5
10.0
7.5
5.0
2.5
0.0
Vachères
Grésigne
St Sauvant
Tronçais
Longchamp
Bercé
Bezange
Lappwald
Göhrde
populations sorted by decreasing latitude
genotype
Hom. ref.
Het.
Hom. alt.

27

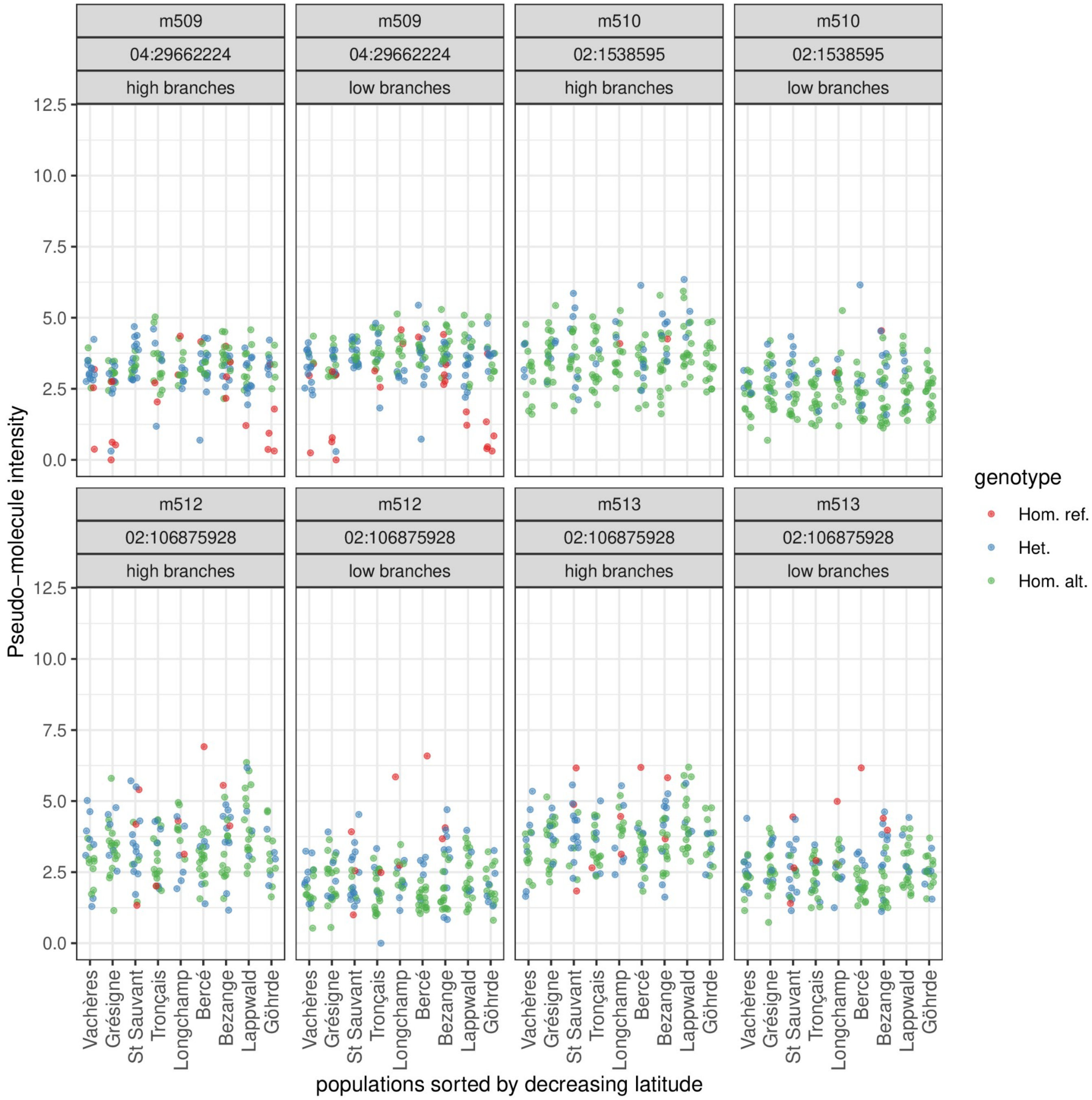

m509
04:29662224
high branches
m509
04:29662224
low branches
m510
02:1538595
high branches
m510
02:1538595
low branches
m512
02:106875928
high branches
m512
02:106875928
low branches
m513
02:106875928
high branches
m513
02:106875928
low branches
Pseudo−molecule intensity
0.0
2.5
5.0
7.5
10.0
12.5
genotype
Hom. ref.
Het.
Hom. alt.
Vachères
Grésigne
St Sauvant
Tronçais
Longchamp
Bercé
Bezange
Lappwald
Göhrde
populations sorted by decreasing latitude

202

28

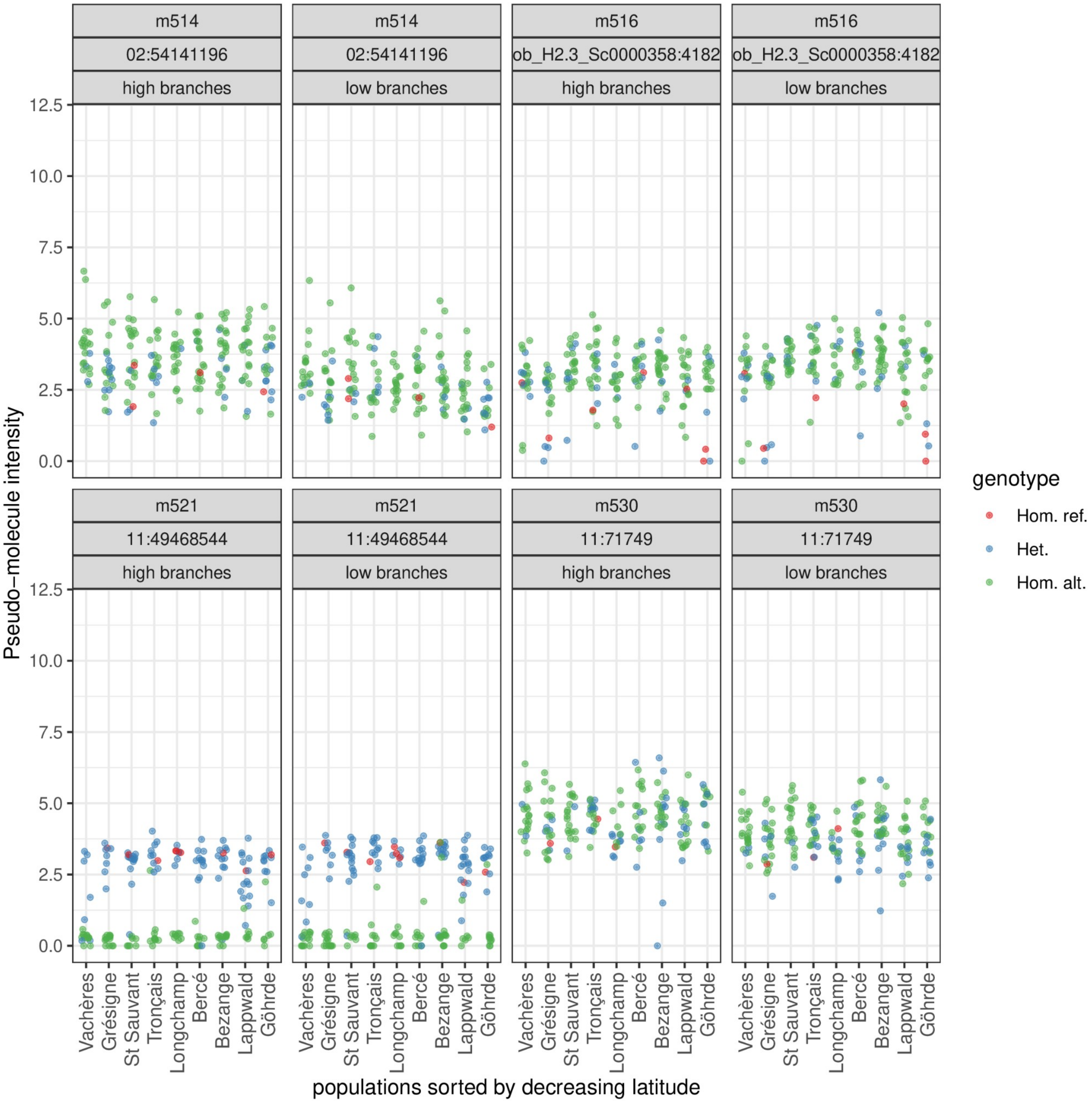
m514
02:54141196
high branches
low branches
m516
ob_H2.3_Sc0000358:4182
m521
11:49468544
m530
11:71749
Pseudo−molecule intensity
0.0
2.5
5.0
7.5
10.0
12.5
Vachères
Grésigne
St Sauvant
Tronçais
Longchamp
Bercé
Bezange
Lappwald
Göhrde
populations sorted by decreasing latitude
genotype
Hom. ref.
Het.
Hom. alt.

29

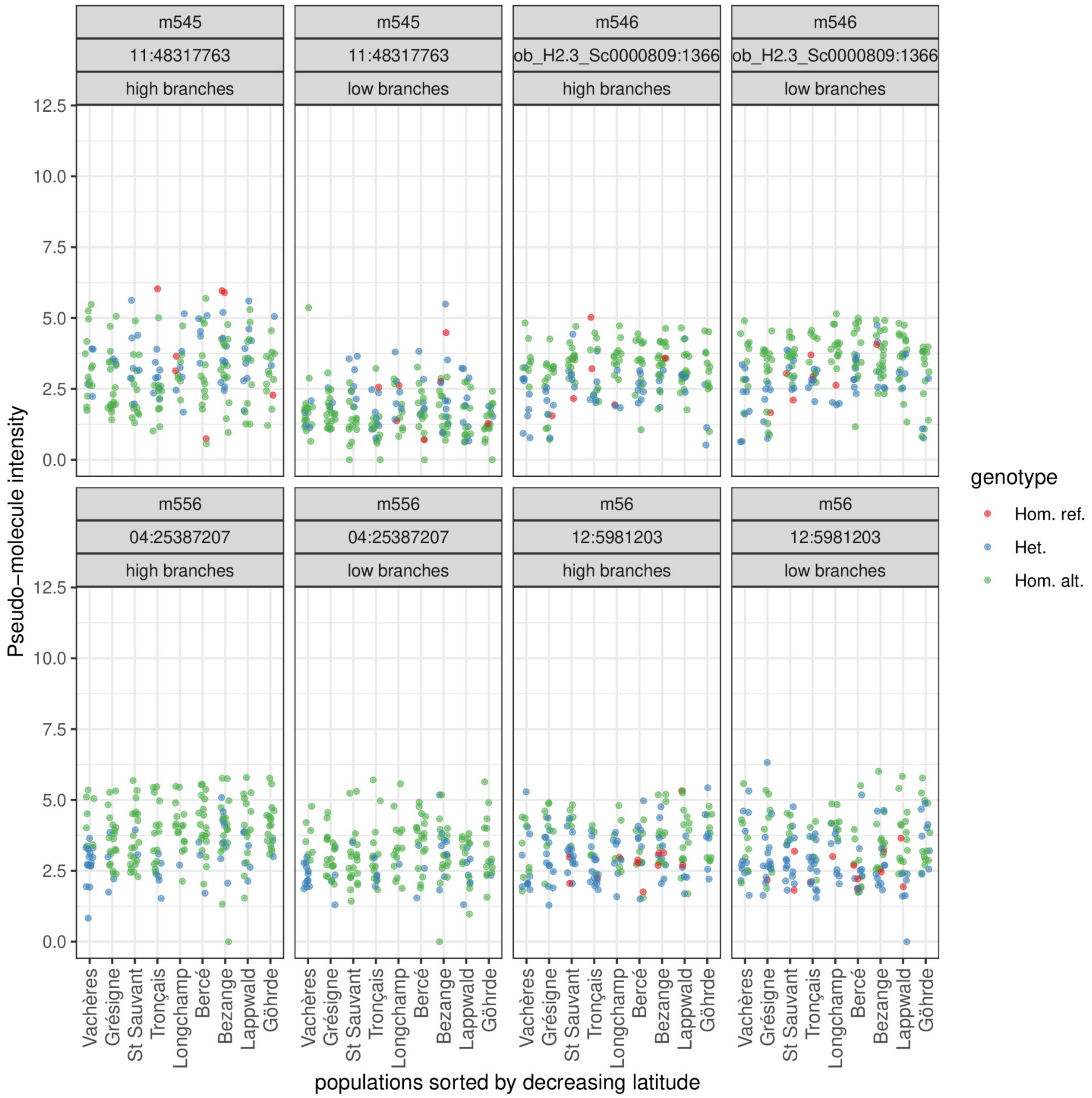
m545
11:48317763
high branches
m545
11:48317763
low branches
m546
ob_H2.3_Sc0000809:1366
high branches
m546
ob_H2.3_Sc0000809:1366
low branches
m556
04:25387207
high branches
m556
04:25387207
low branches
m56
12:5981203
high branches
m56
12:5981203
low branches
0.0
2.5
5.0
7.5
10.0
12.5
Pseudo−molecule intensity
Vachères
Grésigne
St Sauvant
Tronçais
Longchamp
Bercé
Bezange
Lappwald
Göhrde
populations sorted by decreasing latitude
genotype
Hom. ref.
Het.
Hom. alt.

30

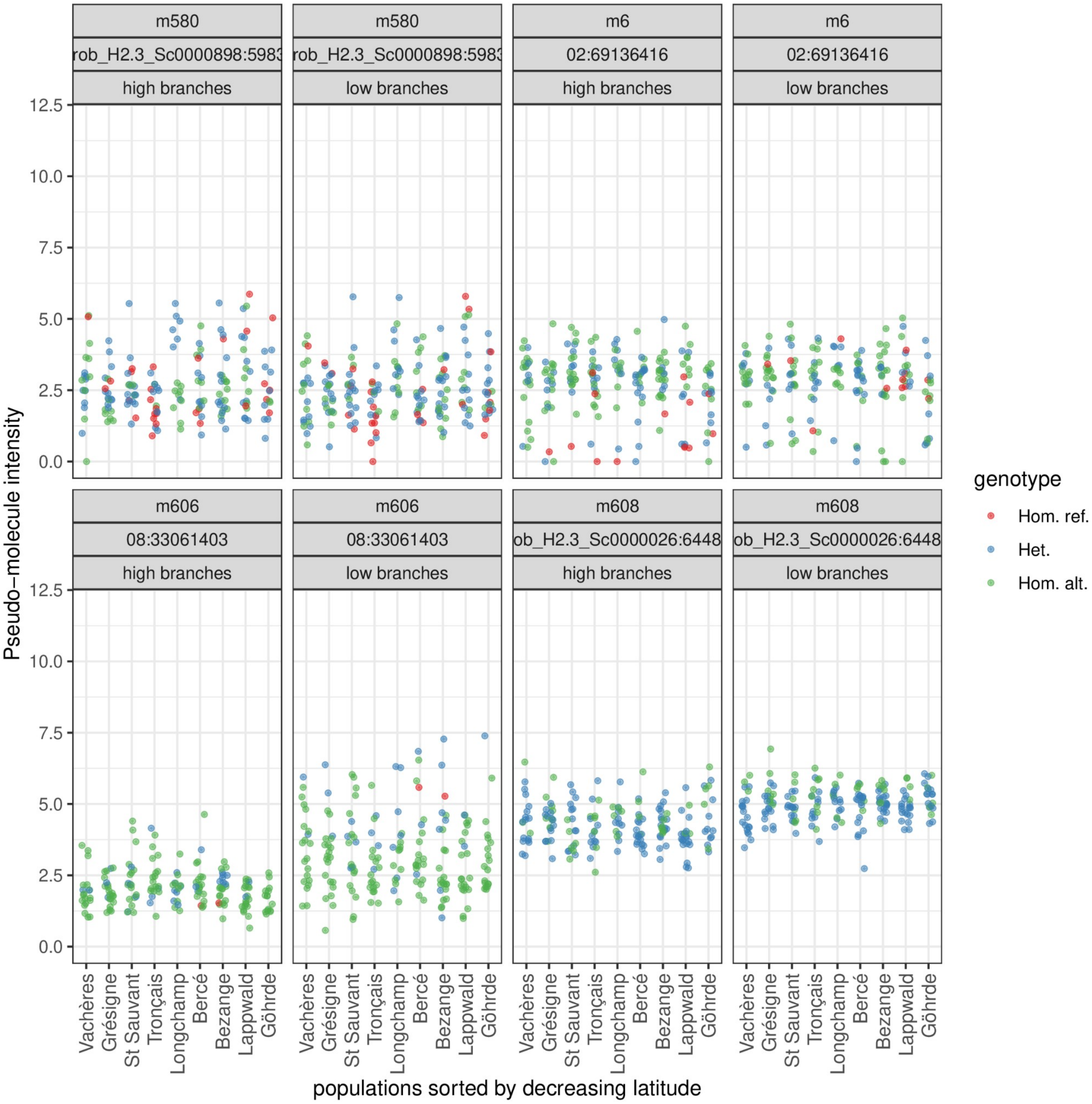
m580
rob_H2.3_Sc0000898:5983
high branches
low branches
m6
02:69136416
m606
08:33061403
m608
ob_H2.3_Sc0000026:6448
Pseudo−molecule intensity
12.5
10.0
7.5
5.0
2.5
0.0
Vachères
Grésigne
St Sauvant
Tronçais
Longchamp
Bercé
Bezange
Lappwald
Göhrde
populations sorted by decreasing latitude
genotype
Hom. ref.
Het.
Hom. alt.

 31

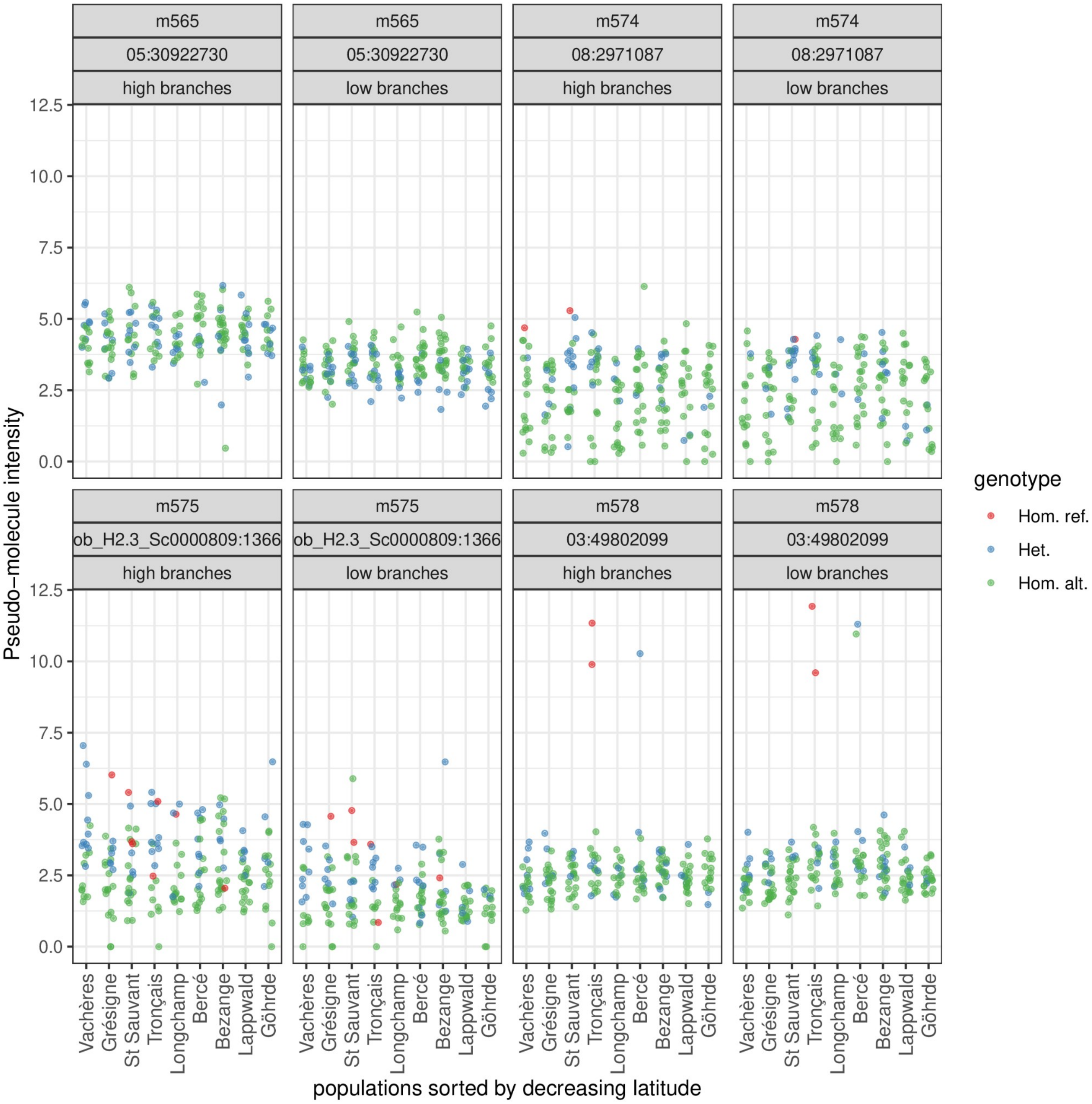
m565
05:30922730
high branches
low branches
m574
08:2971087
m575
ob_H2.3_Sc0000809:1366
m578
03:49802099
Pseudo−molecule intensity
0.0
2.5
5.0
7.5
10.0
12.5
Vachères
Grésigne
St Sauvant
Tronçais
Longchamp
Bercé
Bezange
Lappwald
Göhrde
populations sorted by decreasing latitude
genotype
Hom. ref.
Het.
Hom. alt.

32

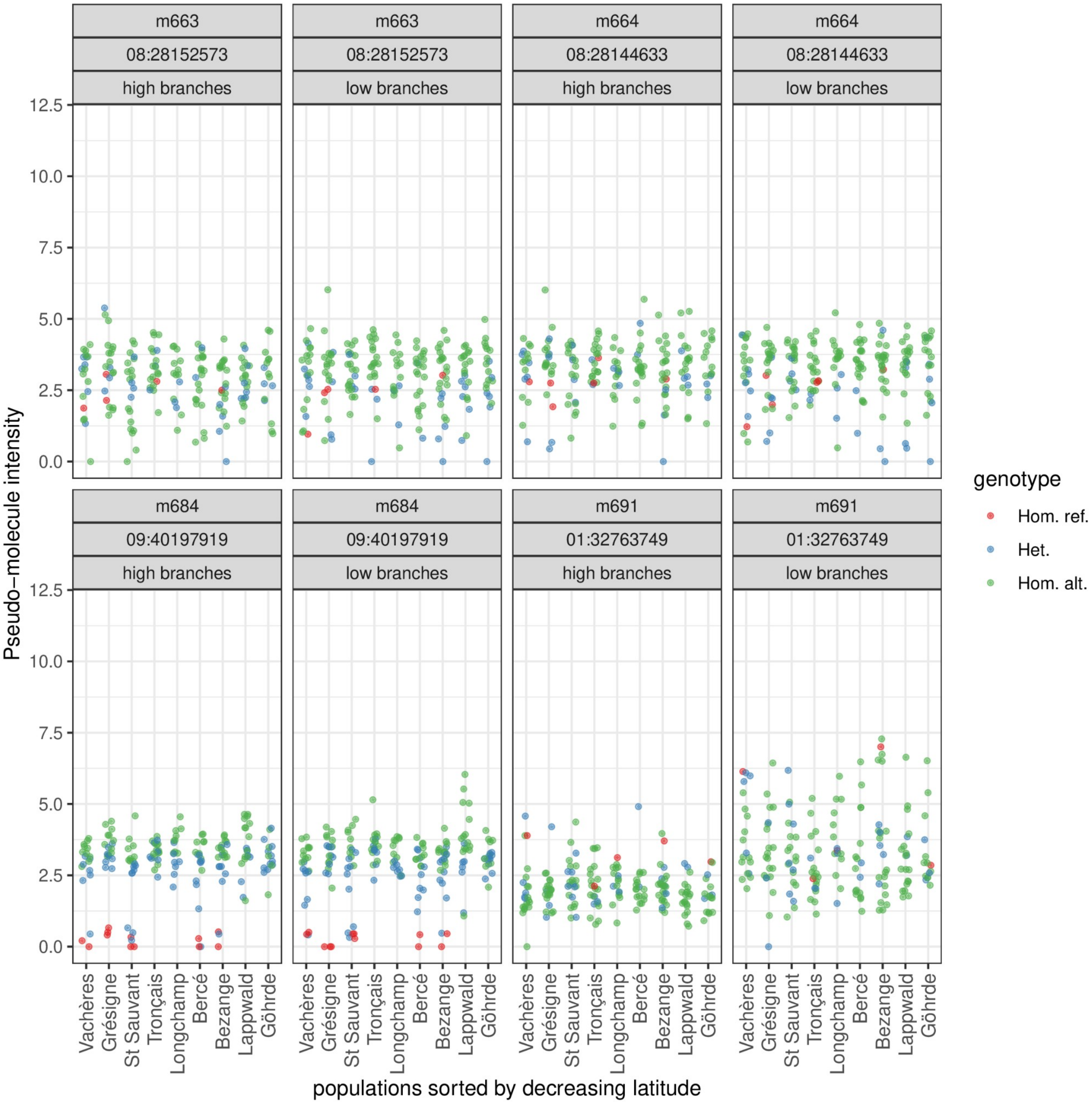
m663
08:28152573
high branches
m663
08:28152573
low branches
m664
08:28144633
high branches
m664
08:28144633
low branches
m684
09:40197919
high branches
m684
09:40197919
low branches
m691
01:32763749
high branches
m691
01:32763749
low branches
Pseudo−molecule intensity
12.5
10.0
7.5
5.0
2.5
0.0
Vachères
Grésigne
St Sauvant
Tronçais
Longchamp
Bercé
Bezange
Lappwald
Göhrde
populations sorted by decreasing latitude
genotype
Hom. ref.
Het.
Hom. alt.

207

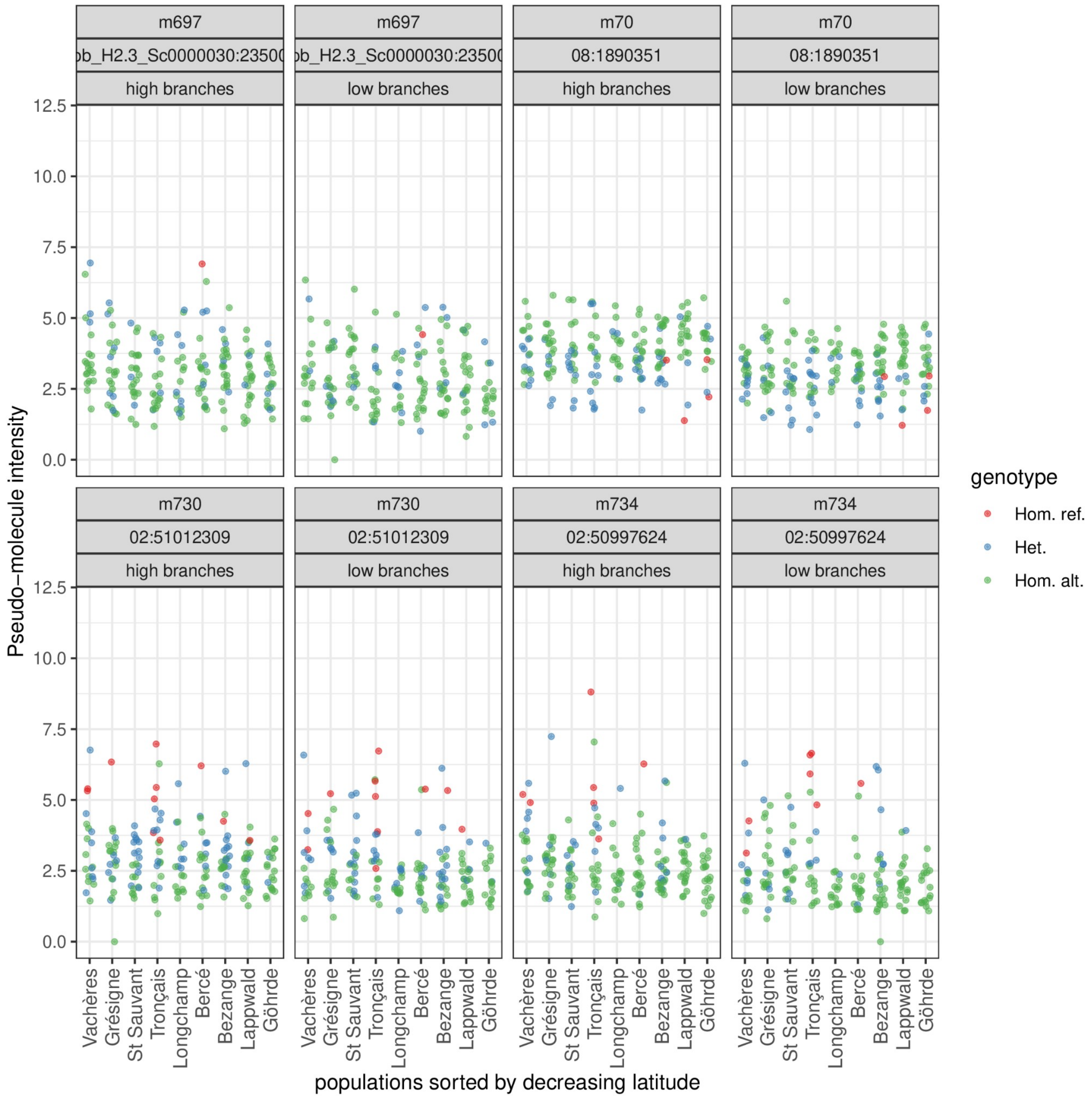
m697
b_H2.3_Sc0000030:23500
high branches
low branches
m70
08:1890351
m730
02:51012309
m734
02:50997624
12.5
10.0
7.5
5.0
2.5
0.0
Pseudo−molecule intensity
genotype
Hom. ref.
Het.
Hom. alt.
Vachères
Grésigne
St Sauvant
Tronçais
Longchamp
Bercé
Bezange
Lappwald
Göhrde
populations sorted by decreasing latitude

208

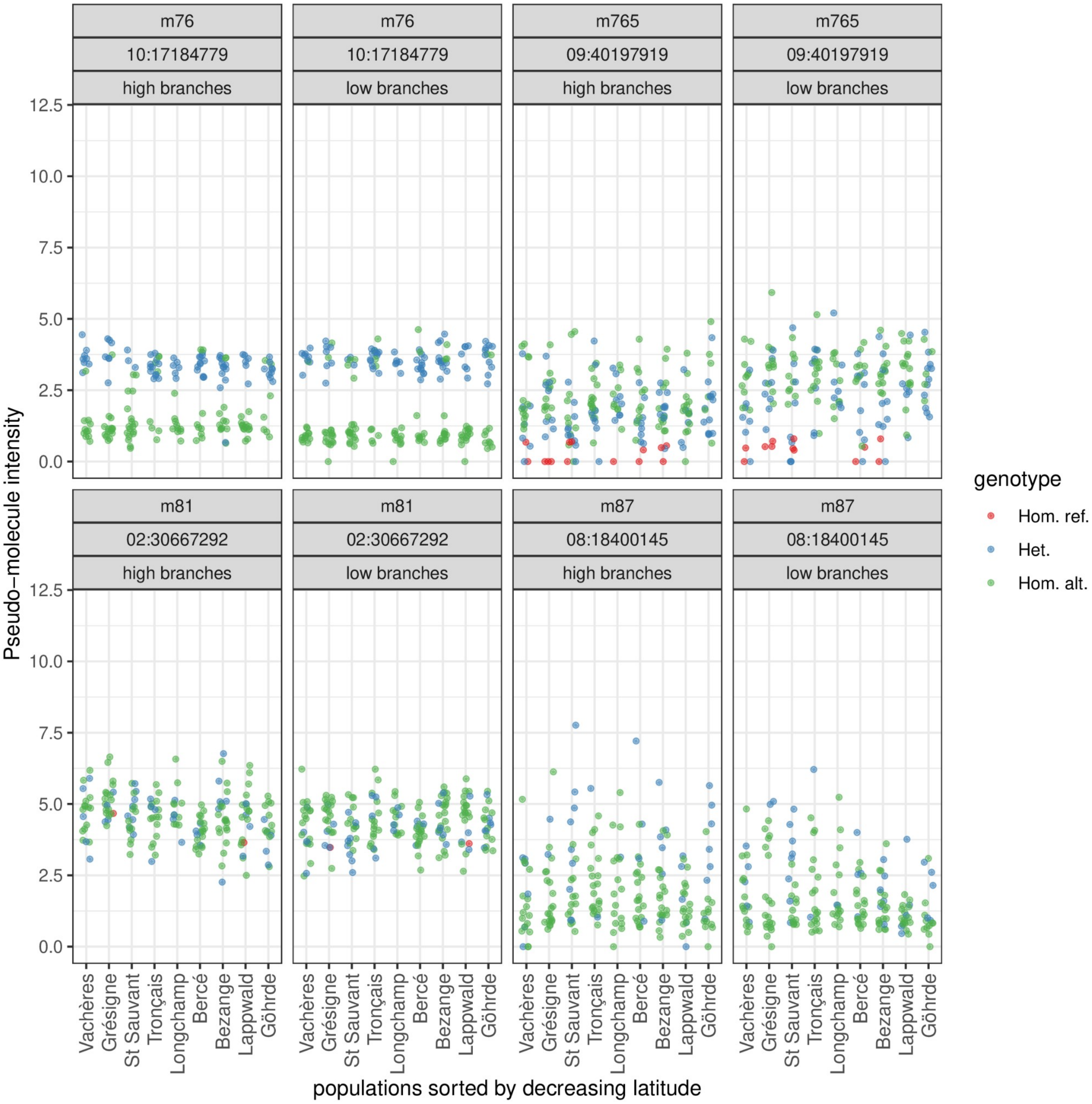
m76
10:17184779
high branches
low branches
m765
09:40197919
m81
02:30667292
m87
08:18400145
Pseudo−molecule intensity
0.0
2.5
5.0
7.5
10.0
12.5
genotype
Hom. ref.
Het.
Hom. alt.
Vachères
Grésigne
St Sauvant
Tronçais
Longchamp
Bercé
Bezange
Lappwald
Göhrde
populations sorted by decreasing latitude

35

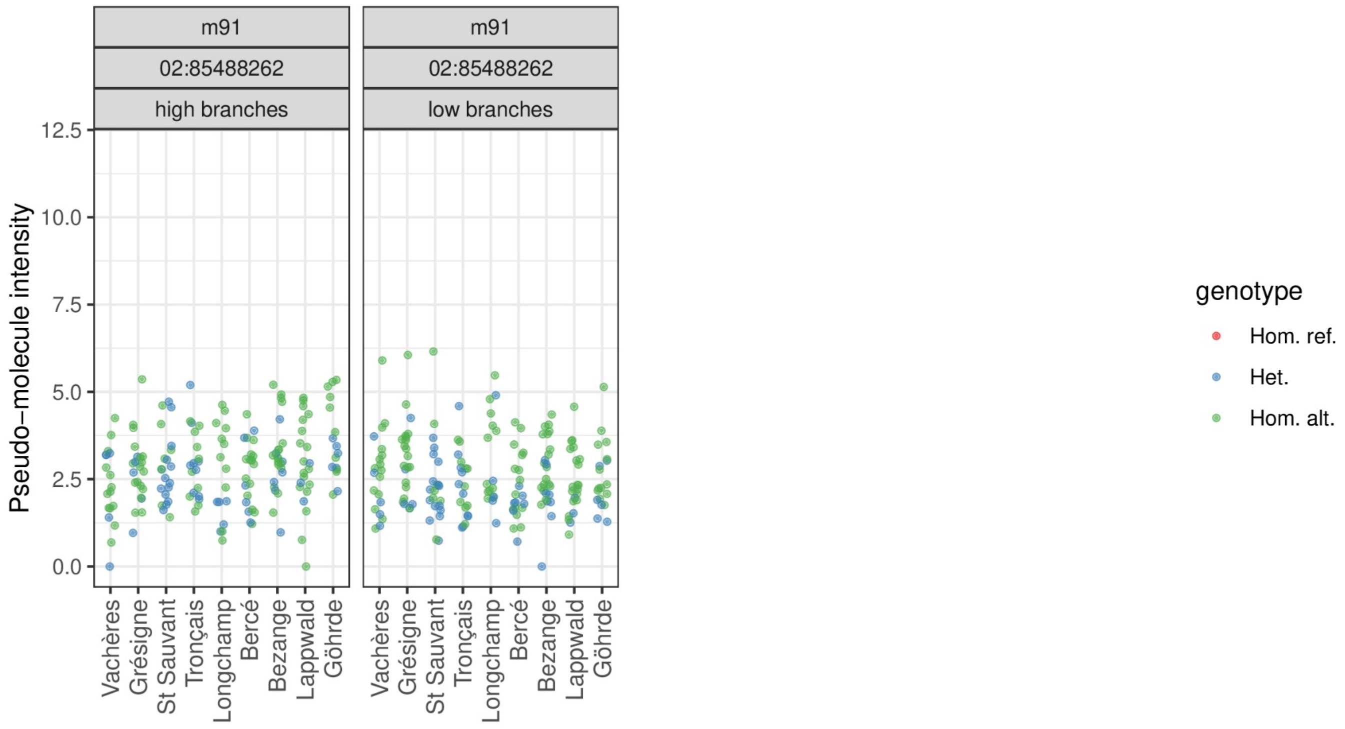
m91
02:85488262
high branches
low branches
Pseudo−molecule intensity
12.5
10.0
7.5
5.0
2.5
0.0
Vachères
Grésigne
St Sauvant
Tronçais
Longchamp
Bercé
Bezange
Lappwald
Göhrde
genotype
Hom. ref.
Het.
Hom. alt.

36
